# Endophytic pyrroloquinoline quinone enhances banana growth and immunity against *Fusarium* wilt for plant-microbe mutualisms

**DOI:** 10.1101/2024.08.20.608638

**Authors:** Shih-Hsun Walter Hung, Man-Yun Yu, Chia-Ho Liu, Tsai-Ching Huang, Jian-Hau Peng, Nai-Yun Jang, Chih-Horng Kuo, Yu-Liang Yang, Ying-Ning Ho, En-Pei Isabel Chiang, Hau-Hsuan Hwang, Chieh-Chen Huang

## Abstract

*Fusarium* wilt has a substantial impact on global banana production, posing a threat to food security worldwide. However, breeding new *Fusarium*-resistant cultivars is difficult and time-consuming. Alternatively, endophytic biostimulants that could combat such pervasive plant diseases provide possible novel solutions. Our prior research demonstrated that a pyrroloquinoline quinone (PQQ)–producing endophytic bacterium, *Burkholderia seminalis* 869T2, can enhance the growth of various plant species and protect bananas from *Fusarium* wilt in the field. PQQ is a peptide-derived redox cofactor known to stimulate mitochondrial biogenesis and metabolism in animals, but its molecular roles, especially in plants, remain to be elucidated. In this study, multi-omics approaches were employed to explore the potential mechanisms through which PQQ influences banana plants. The result of *in situ* imaging mass spectrometry revealed that the endophytic metabolite PQQ does not function through direct antagonism against *Fusarium*. The follow-up transcriptomic profiling shows it could regulate plant respiration, TCA cycle, oxidative phosphorylation, NAD/NADP–dependent dehydrogenases, MAPK signalling, and various phytohormone signalling pathways. Furthermore, PQQ appeared to trigger plant systemic immunity, thereby enhancing plant health and resistance to biotic stress. Beyond that, the complete genome of 869T2 was determined for follow-up comparative genomics analyses, revealing its genetic contexts, potential evolutionary events of PQQ operons among the *Burkholderia* species, and the absence of human virulence-facilitating genes within those PQQ-producing agricultural isolates. In summary, this study facilitates our understanding of PQQ in plant-microbe mutualisms and provides scientific evidence for its future application in agriculture.

**Significance Statement:** *Fusarium* wilt is caused by *Fusarium oxysporum* f. sp. *cubense* tropical race 4 (*Foc* TR4), a notorious soil-borne pathogen that attacks bananas’ vascular system, which critically threatens global banana production and food security. A potential PQQ-producing endophytic strain has been confirmed to protect bananas through *in planta* biocontrol, reducing the morbidity of Banana *Fusarium* Wilt (BFW) disease in the field and promoting the growth of banana plants simultaneously. Our results revealed that the endophytic metabolite PQQ does not function through direct antagonism but triggers plant systemic immunity and coordinates energetic metabolisms, thereby improving the overall health of host plants and enhancing their resistance against *Fusarium* wilt.

## Introduction

Pyrroloquinoline quinone (PQQ), also known as methoxatin, was initially identified as a unique quinone-like cofactor that partners with bacterial glucose and alcohol dehydrogenases, as reported in early studies (1, 2). Subsequent research uncovered a variety of PQQ-dependent dehydrogenases in prokaryotes, which process substances like methanol, alcohol, aldehydes, and sugars (3–5). Initially thought to be exclusive to prokaryotes, PQQ was later discovered in eukaryotic organisms, including animals and plants (6). Serving as a redox cofactor for several dehydrogenases, PQQ is integral to vital metabolic processes across different life forms of kingdoms (7, 8). Its diverse benefits, both in vivo and in vitro, have been demonstrated in numerous studies focusing on microbes and animals. For example, PQQ acts as a powerful antioxidant by neutralizing reactive oxygen species (ROS) (9), it helps maintain the balance of nicotinamide adenine dinucleotide levels (9), and it stimulates the creation of mitochondria (10, 11), thereby enhancing energy-related metabolisms such as oxidative phosphorylation and ATP production (12, 13). However, the impact of PQQ on plant molecular biology is not well understood. Recent findings suggest that PQQ can eliminate ROS in plants similarly to its function in animals (14). Additionally, PQQ appears to enhance the glucose dehydrogenase activity and phosphate-solubilizing abilities of plant growth-promoting bacteria (15–19), and it may also bolster the antifungal defence of plants by supporting plant-associated microbes (20, 21). Nevertheless, the precise molecular mechanisms underlying these effects are still to be elucidated.

On the other hand, *Fusarium oxysporum* f. sp. *cubense* tropical race 4 (*Foc* TR4) is a notorious soil-borne pathogen that attacks the vascular system of bananas, critically threatening banana production and having severe consequences on global food security and the livelihoods of around 400 million people (22, 23). The most effective approach to combat this highly destructive and aggressive fungus is to prevent its initial spread, as it can cause complete yield loss once established in a field (24). Currently, over 80% of global banana production relies on Foc TR4-susceptible germplasm (25). Given the challenge of breeding new varieties or cultivars that are both suitable for the value chain and resistant to disease, new technologies for accelerating precise breeding and biological agents for in-field protection are urgently needed. Some plant symbionts have been reported to protect banana plants against *Foc* TR4 (26–29), and some microbial metabolites function as biostimulants, resulting in plant disease protection through exogenous supplementation (30, 31).

Our previous works isolated and characterised a potential PQQ-producing endophyte, *Burkholderia seminalis* 869T2, which confers strong resistance to *Fusarium oxysporum* f. sp. *cubense* tropical race 4 (Foc TR4) (28, 32). This endophytic strain has been confirmed to protect bananas in field tests through *in planta* biocontrol, reducing the morbidity of Banana *Fusarium* Wilt disease from 24.5% to 3.4% while also promoting the growth of banana plants (28). Besides those positive effects brought on bananas, strain 869T2 and its metabolites have been established as plant-beneficial or stress-resilient endophytic biostimulants that promote the growth of various crops, including *Brassica*, *Lactuca*, *Amaranthus*, *Capsicum*, and *Abelmoschus* (33, 34). The strain has also been established as a versatile endophytic biostimulant for leafy vegetables. It significantly promotes plant growth and enhances crop quality by increasing harvested head weight, size and nutrient composition. Additionally, a two-week acceleration in harvest time and 50% savings on fertilising costs result in reducing costs and increasing product value simultaneously, which provides a model for in-field endophytic biostimulants-assisted smart agricultural management for lettuce production (34). On the other hand, the bioremediation availability of 869T2 for the degradation of persistent organic pollutants (POPs), such as dioxins, under an aerobic environment through 2-haloacid dehalogenase has also been demonstrated (35). These results emphasize the importance of endophytic bacteria, warranting further investigation due to their enigmatic symbiotic relationships with hosts and potential applications in the agro-industry.

In this study, our goal was to better understand the molecular mechanisms of endophytic PQQ *in planta*, particularly in the context of plant-microbe mutualistic symbiosis and the defence against pathogens. Our result revealed that endophytic PQQ has no direct antagonism activity to inhibit the growth of *Fusarium* species. Instead, it promotes the overall health of host plants by triggering plant immunity through the modulation of multiple phytohormones and MAPK signalling pathways. Additionally, it coordinates their energetic metabolisms, including the TCA cycle, oxidative phosphorylation, and the biosynthesis and activity of NAD/NADP-dependent enzymes. Meanwhile, the results of comparative genomics shed light on the evolution of the pqq operon and other genetic contents, thus offering clues about the co-evolution and mutualistic relationships among those plant-associated bacteria with their hosts.

## Results

### Species reclassification of strain 869T2 to *Burkholderia seminalis*

Strain 869T2 was initially identified as *Burkholderia cenocepacia* with its draft genome sequence (32). However, updated analysis of molecular phylogeny and gene content indicated that this strain is more similar to *B. seminalis* (36). Therefore, complete genome sequencing and comparative analyses were performed in this work to determine its species assignment.

For hybrid *de novo* assembly, 2 x 3,178,899 (∼1.9 Gb) of Illumina and 46,153 (∼0.5 Gb) of untrimmed ONT raw reads were generated, which provided ∼238– and ∼65-fold coverages for this genome, respectively. The assembly result indicated that 869T2 has three circular chromosomes totalling 8,022,336 bp and an average GC content of 67.03%. The comparison against a reference Burkholderiales dataset identified 687 complete BUSCOs (99.9%), one fragmented BUSCOs (0.1%), and no missing BUSCOs within this genome assembly, supporting that the genome assembly is complete. The annotation showed that this genome contains six complete sets of rRNA genes, 67 tRNA genes, four ncRNAs, 7,090 protein-coding genes and 112 pseudogenes (Table S1).

For the species assignment, the newly generated complete genome sequence was utilised for comparative analysis with key representatives of *B. seminalis* and *B. cenocepacia* (Table S2) based on genome-wide average nucleotide identity (ANI) and core-genome phylogeny. The result showed that 869T2 shares ∼93-95% of its genome with other *B. seminalis* strains and has > 99% ANI (Fig. 1A). In contrast, 869T2 shares only ∼79-82% of its genome and has < 93% ANI when compared to *B. cenocepacia* strains. Additionally, 3,018 single-copy genes shared by all strains were identified and used to produce a concatenated alignment containing 987,390 aligned amino acid sites for maximum likelihood phylogeny inference (Fig. 1A). Based on 1,000 resamplings, the resulting tree had 100% bootstrap support value for all internal branches and 869T2 is a member of the *B. seminalis* clade. Based on these results, strain 869T2 can be confidently re-classified as a member of *B. seminalis*.

**Fig. 1.**
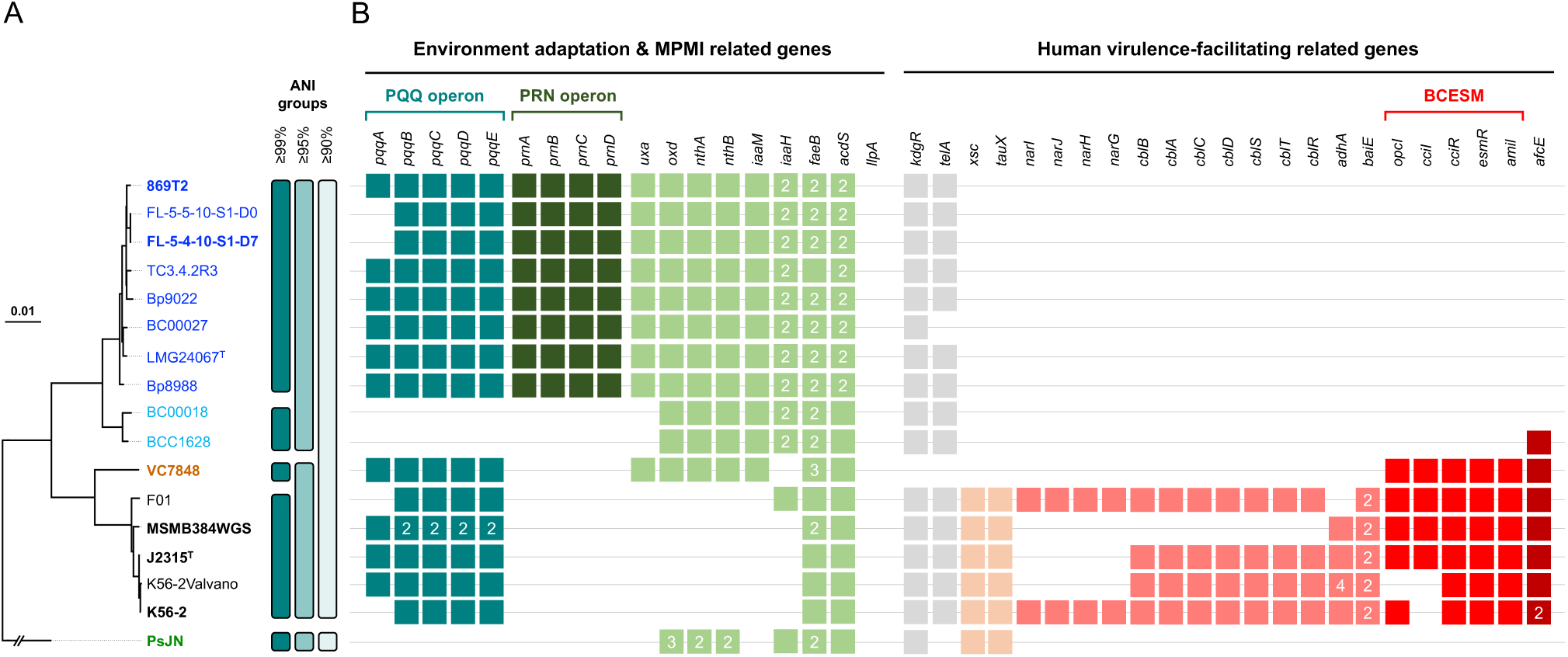
Evolutionary and comparative genomics among strain 869T2 and its relatives. **A** Maximum likelihood phylogeny based on a concatenated alignment of 3,018 single-copy genes shared by all genomes (987,390 aligned amino acid sites). All internal nodes received 100% bootstrap support based on 1,000 resamplings. *Paraburkholderia phytofirmans* PsJN^T^ is included as the outgroup. The superscripts ‘T’ following the strain names indicate type strains and the complete genomes are presented in bold. Grouping based on different genome-wide average nucleotide identity (ANI) thresholds is plotted to the right of the phylogenic tree. **B** Gene content analysis. The distribution of key genes involved in environmental adaptation, molecular plant-microbe interactions (MPMI), and human virulence-facilitating (VIR) are plotted. The colour boxes indicate gene presence, and the numbers within the boxes indicate copy numbers of multi-copy genes. Abbreviations: PQQ, pyrroloquinoline quinone. PRN, pyrrolnitrin. BCESM, *Burkholderia cepacia* epidemic strain marker. Detailed information regarding the strains and target genes are provided in Table S2 and Table S3, respectively.

### Gene content analysis between *B. cenocepacia* and *B. seminalis*

For more in-depth analysis, the gene content of representative *B. seminalis* and *B. cenocepacia* strains was compared. Specifically, genes known to be related to environmental adaptation and molecular plant-microbe interactions (MPMI) were the main targets (Fig. 1B and Table S3). A distinct differentiation was observed for genes encode for nitrile hydratase cluster (*nthAB*), phenylacetaldoxime dehydratase (*oxd*), galacturonate metabolism (*uxa*), indoleacetic acid (IAA) biosynthesis (*iaaHM*), and pyrrolnitrin biosynthesis (*prnABCD*); these genes were conserved among *B. seminalis* strains but mostly absent among *B. cenocepacia* strains (Fig. 1B). Additionally, *B. seminalis* strains mostly have two copies of genes for feruloyl esterase (*faeB*) and ACC deaminase (*acdS*). On the other hand, genes involved in virulence to human (*esmR*, *amiI*, *cciI*, *cciR*, and *opcI*) (37) were found only in *B. cenocepacia* strains, but absent in all of the *B. seminalis* strains examined (Fig. 1B), which mitigate the risk of using *B. seminalis* strains such as 869T2 for agricultural application. Finally, the *pqq* operon, encoding essential genes responsible for the biosynthesis of PQQ, is conserved in most of the strains examined.

### PQQ cannot antagonise *Fusarium in vitro*

Given that 869T2 has various plant-growth-promoting (PGP) traits and abundant MPMI-related genetic contents, we used the ultra-high performance liquid chromatography-tandem mass spectrometry (UHPLC-MS/MS) technique to determine its metabolome and target the specific chemicals responsible for the PGP or pathogen resistance potential. The results showed that 869T2 produces multiple phytohormones, including auxin, cytokinin, gibberellic acid, ethylene and salicylic acid, as well as the corresponding regulatory factors were confirmed (Table S4). Besides, 31 cyclic dipeptides (CDPs) were identified; some CDPs were found as antimicrobial substances and antibiotics (Table S5). Furthermore, along with a biosynthesis operon of pyrrolnitrin, an antibiotic that suppresses *Fusarium* species (38), four of its antifungal derivates: iso-, oxy– and fluoro-pyrrolnitrin and 3-chloro-4-(2-nitrophenyl)-1H-pyrrole were also detected. These results suggested that 869T2 could inhibit the growth of fungi, *e.g.*, *Foc* TR4, via specific antifungal compounds.

A follow-up *in situ* metabolic investigation using the matrix-assisted laser desorption ionization-time of flight (MALDI-TOF) imaging mass spectrometry (IMS) technique also confirmed that 869T2 produced pyrrolnitrin and its derivatives, oxypyrrolnitrin or 3-Chloro-4-(3-chloro-4-fluoro-2-nitrophenyl) pyrrole, to inhibit the growth *Foc* TR4 *in vitro* (Fig. 2A). Strikingly, the signal of endophytic biostimulants PQQ was found and was only detected when incubating without *Foc* TR4 (Fig. 2B); the endophytic PQQ production was further confirmed through gas chromatography-mass spectrometry (GC-MS) by comparing to the synthetic PQQ standard (>98% purity) (Fig. S1). Previous studies suggested mutagenesis on *pqqC* and *pqqE*, essential genes for PQQ biosynthesis, affects the bacterial antimicrobial activity *in vitro* or *in planta* (20, 21, 39). However, our imaging mass results suggested that the production of PQQ seems irrelevant to the *Fusarium* inhibition. To explain this paradoxical relationship, an antagonism assay was performed using levels of synthetic PQQ standards as treatments and *B. seminalis* 869T2 as a control. The results showed that PQQ has no positive effects on inhibiting the growth of *Foc* TR4 but improves it instead (Fig. 2C). Based on this evidence, we raise another possibility that PQQ may induce plant immunity, resulting in antimicrobial activity in plants. To examine our hypothesis, synthetic PQQ was exogenously supplemented to banana plantlets, and the morbidity to *Fusarium* wilt and plant transcriptomics were investigated to unveil the obscure molecular mechanisms among them.

**Fig. 2.**
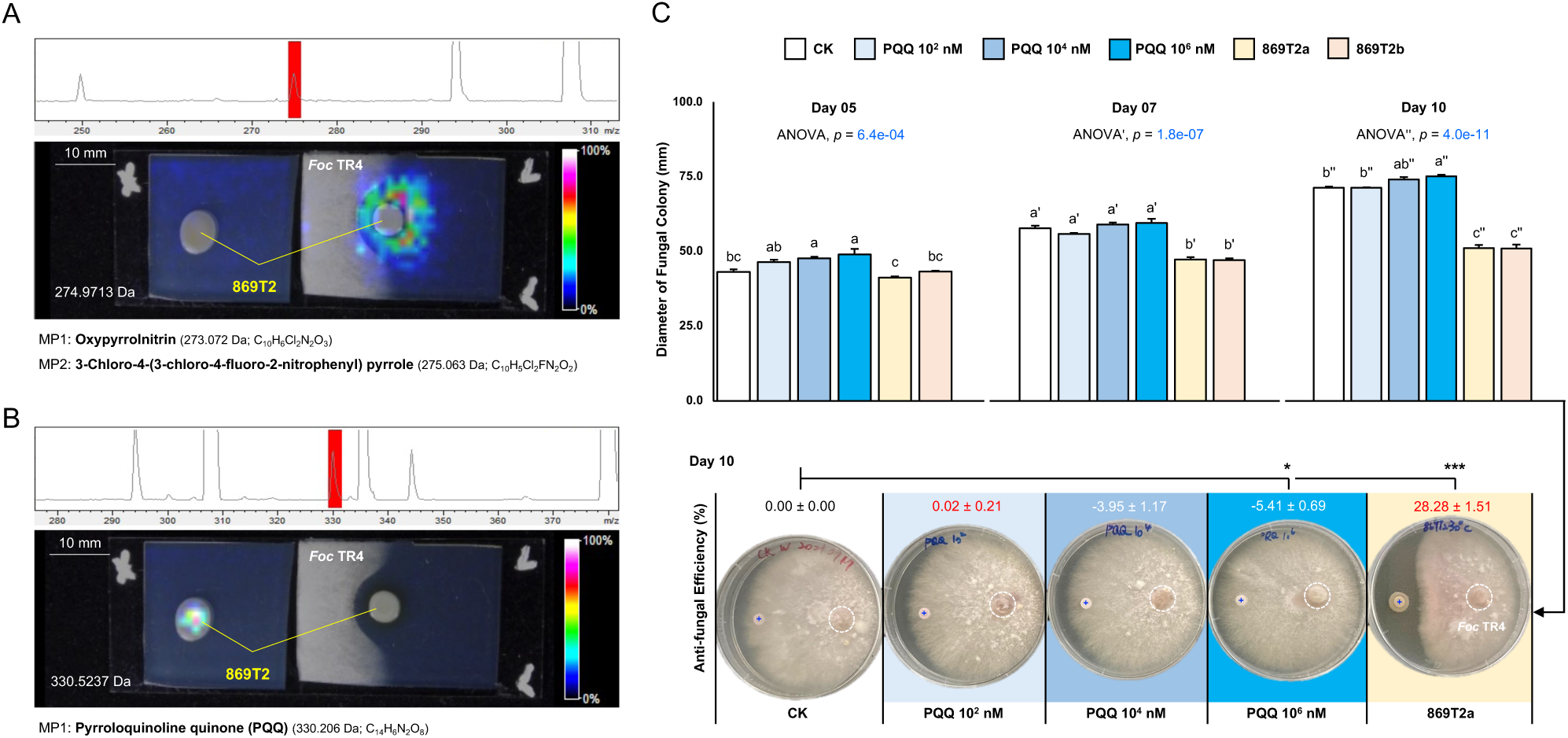
Identification and quantification of potential endophytic biostimulants. *In situ* imaging mass results of endophyte-pathogen co-incubation under the metabolic signalling detection under **A** 274.9713 m/z and **B** 330.5237 m/z, respectively. 869T2, *Burkholderia seminalis* 869T2. *Foc* TR4, *Fusarium oxysporum* f. sp. cubense tropical race 4. MP, metabolite of prediction. Colour keys refer to the metabolic abundance of detection. **C** Antagonism assay for testing different concentrations of pyrroloquinoline quinone (PQQ) against *Foc* TR4 *in vitro*. PQQ was prepared as 10^2^, 10^4^, and 10^6^ nM for the assay. *Burkholderia seminalis* 869T2 precultured under precultured under 30°C (869T2a) or 37°C (869T2b) was set as the positive control, and the ddH_2_O was set as the negative control (CK). The sample disk of each treatment and Foc TR4 were highlighted by a blue cross and a white dashed circle, respectively, within the plate images. Data were presented as mean ± SEM, N = 3. Statistical significances between samples were indicated by different letters (p<0.05) or by asterisks (*p<0.05; ***p<0.005); otherwise, not significant.

### Exogenous supplementation of PQQ induces the growth and development of banana plants

Firstly, 5 nM, 50 nM, 100 nM and 1000 nM PQQ, hereinafter referred to as P05, P50, P100 and P1000, respectively, were examined on tissue culture banana plantlets; about twenty biological replications were set for each treatment. At 48 days after treatment (DPT), the results showed that exogenous PQQ supplementation contributes to the differentiation of both shoot and root (Fig. 3A-C). Among them, by estimating using the developmental indicator N_50_, defined as the shoot/root length of the shortest organ at 50% of the total shoot/root length, P100 seems an ideal dosage for triggering plant leaf and root development. For root development: (a) P100 had three plantlets without visible root development (CK had seven); (b) the root N_50_ of P100 was at level 7-8 (CK was at level 3-4); (c) the maximum root order of P100 was at level >9 (CK was at level 5-6) (Fig. 3B). For leaf development: (a) P100 had four plantlets without visible leaf development (CK had the same); (b) the leaf N_50_ of P100 was at level 3 (CK was at level 2); (c) the maximum leaf order of P100 was at level 5 (CK was at level 3) (Fig. 3C).

**Fig. 3.**
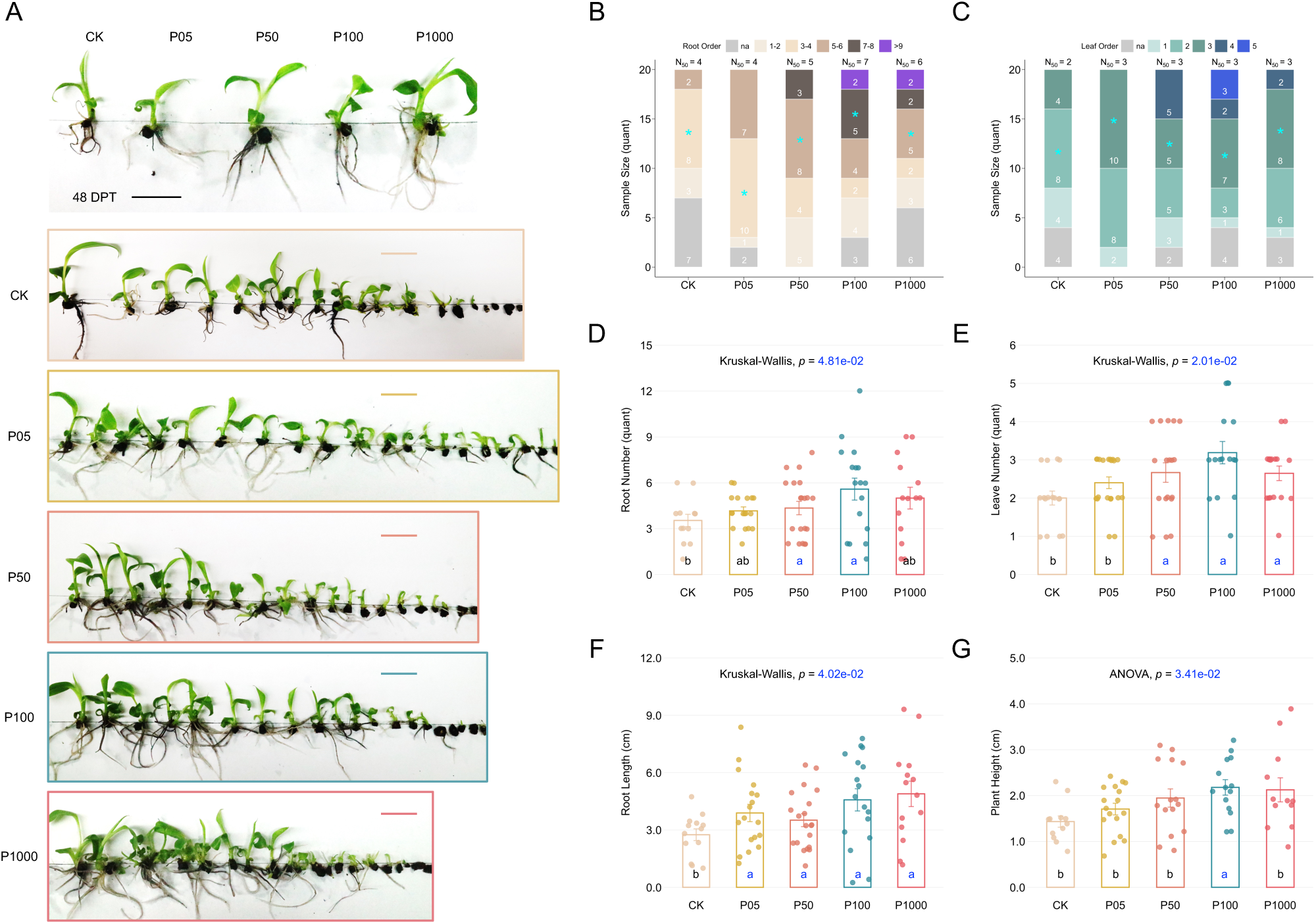
Exogenous supplementation of PQQ benefits the differentiation and development of banana plants in tissue culture conditions. **A** Phenotypic photographs of banana plants in response to different concentrations of supplementary pyrroloquinoline quinone (PQQ) at 48 days after treatment (DPT). CK, P05, P50, P100 and P1000 indicate the 0 nM, 5 nM, 50 nM, 100 nM and 1000 nM PQQ treatment, respectively. Scale bars, 3 cm. The distribution of **B** root number and **C** leaf number of banana plants at 48 DPT. Numbers in stacked bar charts indicate the sample size. Asterisks highlight the distribution groups of defined N_50_ for plant growth-promotion evaluation, as mentioned in Materials and Methods. na, non-differentiated samples. **D** Root number, **E** leaf number, **F** root length and **G** plant height of banana plants at 48 DPT, excluding non-differentiated samples in **B** or **C**. Each dot represents a sample. Data are mean ± SEM; different letters indicate significant differences based on multiple comparisons, Tukey method after ANOVA or Dunn method after the Kruskal-Wallis test.

Besides, significantly more root and leaf numbers were both observed in P50 and P100 (Fig. 3D,E); all treatments significantly elongated the root length (Fig. 3F) while only P100 promoted the plant height compared to the CK with statistical significance (Fig. 3G). These positive effects on plant growth and development were found to remain after the bananas were transplanted to soil pots at least until 137 DPT (Fig. S2). Given that P1000 did not constantly improve plant growth and even contributed less than P100, we hypothesise that a high dosage of PQQ supplementation may inhibit plant growth. Therefore, additional experimental batches, including P100, P1000, and P2000 (2000 nM PQQ), were performed to delineate the boundary of its working concentration. The phenotypic results together suggested that the supplementation concentration over 100 nM had no significant positive effects and, in some cases, harmed the plants instead (Fig. S3). For instance, the fresh weight and plant height of the 1000 nM treatments decreased from 0.66 ± 0.06 g to 0.59 ± 0.08 g (Fig. S3B) and from 2.46 ± 0.13 cm to 2.25 ± 0.12 cm (Fig. S3E), respectively. Accordingly, PQQ concentrations over 100 nM were considered to affect plants negatively and were excluded from the follow-up experiments.

### Transcriptomic landscape of banana plants in response to PQQ

To unveil the molecular roles of PQQ in plants, total RNA was extracted from the CK and 100 nM PQQ treatments, respectively, and used for the following transcriptomic analysis. Totally ∼6.5 Gb raw reads were obtained from each sample through the Illumina Hiseq4000 2 x 150 bp sequencing. After mapping the resultants to the banana reference genome (NCBI TaxID: 214687), a total of 25,070 variables were identified (Fig. 4A). Among them, 1,181 and 592 differentially expressed genes (DEGs) were identified through gene ontology (GO) enrichment analysis (*p* < 0.05 and FC >2) and Kyoto Encyclopedia of Genes and Genomes (KEGG) analysis (*p* < 0.05 and FC >1), respectively (Fig. 4B,C).

**Fig. 4.**
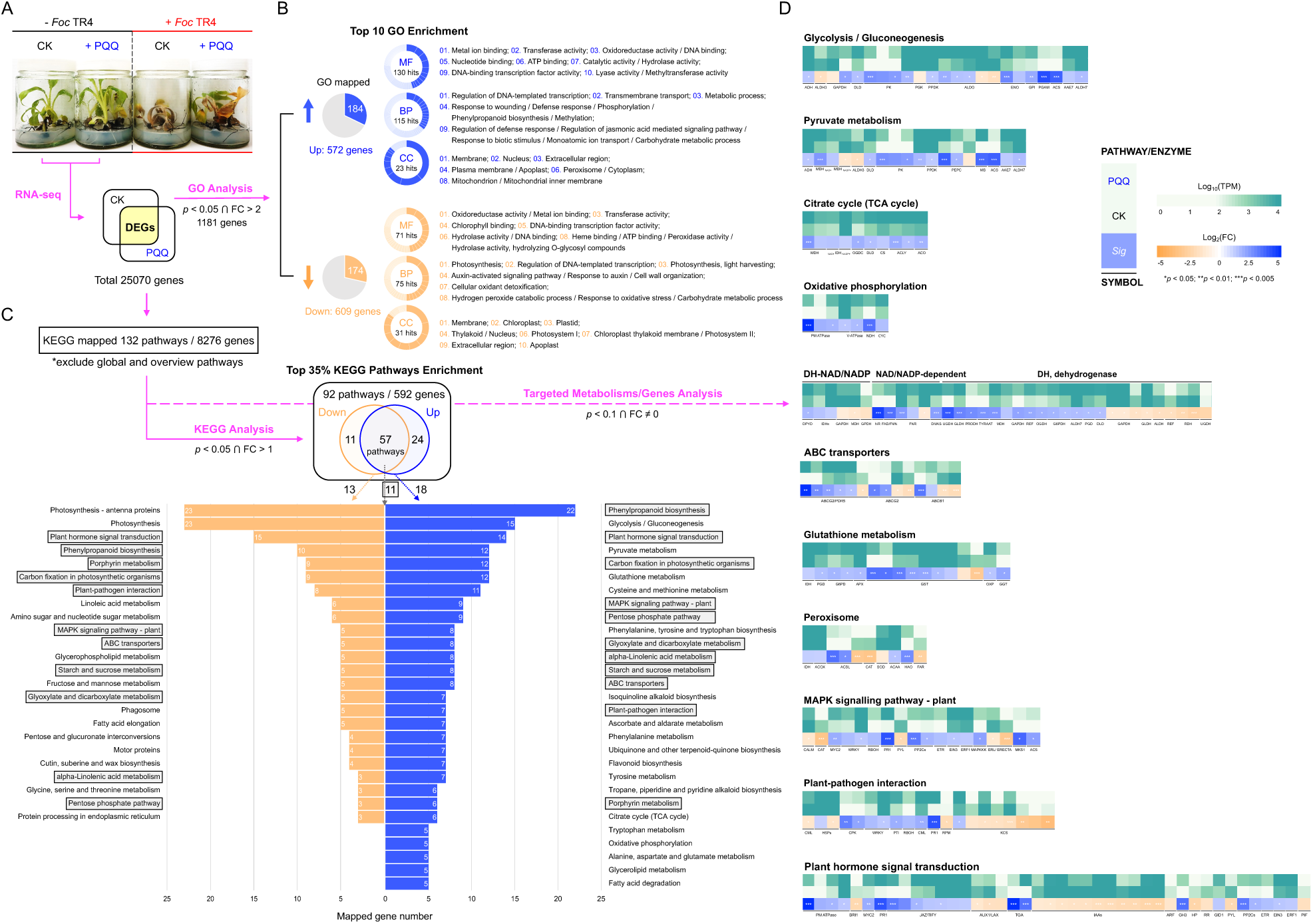
Transcriptomic landscape of banana plants in response to PQQ. **A** Phenotypes of banana plant treated with 100 nM PQQ and infected by *Fusarium oxysporum* f. sp. cubense tropical race 4 (Foc TR4) at tissue culture growing stage. Total RNA was collected from these plants for the follow-up transcriptomic analysis. Differentially expressed genes (DEGs) defined by certain criteria were then subjected to **B** Gene Ontology (GO) terms enrichment analysis, **C** Kyoto Encyclopedia of Genes and Genomes (KEGG) pathways analysis and **D** targeted metabolisms and genes expression analysis. **B** The number inside the pie chart indicates the number of mapped GO terms, and the number inside the donut chart indicates the total hits of DEGs by the top 10 mapped GO terms presented orderly at the right of the donut chart. **C** The number inside the pie chart indicates the number of total DEGs-mapped KEGG pathways; the top 35% were labelled beside arrows linking to the column chart below; the number inside the column chart indicates the number of mapped DEGs; those mapped KEGG pathways shared by up– and down-regulated DEGs were marked by grey boxes. **D** Heatmaps showed the specific gene expression of targeted KEGG pathways. Details of gene symbols can be accessed in Table S7. The colour keys refer to the transcripts per kilobase million (TPM) and the corresponding fold changes (FC). Statistical significances between samples were indicated by asterisks (*p<0.05; **p<0.01; ***p<0.005); otherwise, not significant.

The GO analysis showed that molecular function terms like regulations in heme binding, peroxidase activity, ATPase activity, etc. were found to be enriched regulated. Several DEGs were identified as involved within the chloroplast and mitochondria-related metabolisms within biological processes and cell components terms (Fig. 4B). For KEGG, the DEGs were assigned to 92 pathways, and the 11 pathways, including phenylpropanoid biosynthesis, plant hormone signal transduction, ABC transporters, MAPK signalling pathway – plant, plant-pathogen interaction, etc., were further targeted as they were identified as enriched regulated from both down– and up-regulated (Fig. 4C). The detailed profiles of GO and KEGG analyses were provided in supporting information (Tables S6,S7). Besides, the DEGs were accessed to 11 metabolic pathways or enzymatic genes for further analysis based on our enrichment analysis and previous studies to explore the molecular roles participating within plant physiology (Figs. 4D,S4-S19). Considering the transmission of PQQ across the host and microbe, the ABC transporters ABCB1 and ABCG2/PDR5, which are capable of transporting small molecules, were identified as being regulated with statistical significance (Figs. 4D, S4). Echoing previous works that showed that PQQ has reactive oxygen species (ROS) scavenging and stress-resilience availabilities, 16 and 10 genes of glutathione and peroxisome metabolisms were found to be differentially regulated, respectively (Figs. 4D,S5,S6).

Interestingly, abundant genes responsible for plant respiration and energic metabolisms were found mainly mediated: genes encode essential proteins for the TCA cycle (*e.g.*, citrate synthase, aconitase, isocitrate dehydrogenase, α-ketoglutarate dehydrogenase and malate dehydrogenase) (Figs. 4D,S7) and oxidative phosphorylation (*e.g.*, plasma membrane ATPase, V-type proton ATPase, mitochondrial external alternative NAD(P)H-ubiquinone oxidoreductase B2 and cytochrome c) (Figs. 4D,S8). Genes involved in the glycolysis/gluconeogenesis pathway and pyruvate metabolism (*e.g.*, acetyl-CoA synthetase, aldo-keto reductase, dihydrolipoyl dehydrogenase, phosphoenolpyruvate carboxylase, pyruvate kinase, pyruvate phosphate dikinase, etc.) were also been up-regulated (Fig. 4D,S9,S10) (Table S7). These results strengthen the perspectives on the participation of PQQ in intercellular electron transmission and redox reactions in plant cells. We also analysed those enriched regulated NAD/NADP-dependent enzymes and dehydrogenase to dig into their coordination (Table S8). Although PQQ was mostly suggested to mediate lactate dehydrogenases in animal and microbial cells, these regulations were not found significantly within plant cells. Instead, its regulations on the other dehydrogenases corresponding to aldehyde, malate, glutamate, isocitrates, etc., while ferredoxin reductases were identified as enrichment with significance (Fig. 4D) (Table S7). This result distinguishes the divergence of molecular roles and functions of PQQ as (NAD/NADP-dependent) dehydrogenase among different life kingdoms.

In view of the PGP and anti-*Fusarium* properties of PQQ, the DEGs involved in phenylpropanoid metabolisms, phytohormone biosynthesis and signal transduction, the MAPK signalling pathway, and plant-pathogen interaction were focused. Among them, the phenylpropanoid metabolism for the biosynthesis of structural lignin components, which were essential to the plant cell wall construction and strength and rigidity, were been induced (Fig. S11). The tryptophan metabolism for indole acetate (IAA) and biosynthesis pathways of several chemicals with biotic or abiotic stresses alleviating and with antioxidant activities like lutein and zeaxanthin, carotenoids, dopamine, melatonin and flavonoids were also found induced by PQQ (Fig. S12-S16). For plant hormone signal transduction regulations, auxin and stress-associated pathways were particularly regulated, and pathways for those stress hormones, e.g., ethylene, jasmonic acid, and salicylic acid, were significantly enhanced (Fig. S17). Furthermore, in a total of 23 DEGs of the plant’s MAPK signalling and plant-pathogen interaction, metabolisms were also induced, which included various pathogen-associated transcriptional factors, ethylene regulations and pathogenesis-related proteins, suggesting putative immune responses to protecting plants against pathogens could be pre-triggered by the exogenous supplementation of PQQ (Figs. S18,S19).

### Exogenous supplementation of PQQ boosts the immunity of banana plants

The transcriptomics results confirmed previous findings that mutagenesis of PQQ biosynthesis genes negatively impacts microbial biocontrol ability in plants (20, 21, 39). These results also supported our hypothesis that PQQ coordinates plant growth and immunity to protect the host plant against pathogens. To determine if the antifungal activity observed in tissue culture (Figs. 4A, 5A) persists under soil growing conditions, we transplanted banana plants treated with different concentrations of PQQ into soil pots infected with Foc TR4. We then evaluated four indicators of Panama Disease—leaf yellowing (LY), pseudostem splitting (SS), pseudostem bending (SB), and rhizome dark brown discolouration (RB)—to assess the role of PQQ in protecting plants against *Fusarium* wilt of bananas (Fig. 5B, C). Three months after the *Foc* TR4 infection, the Panama Disease indicators of plants growing in soil conditions were investigated. The unweighted pair group method with arithmetic mean (UPGMA) dendrogram clustering and the principal component analysis (PCA) were performed for an integrative evaluation of the diseasing patterns. The UPGMA results indicated that the LY, RB, and SB diseasing patterns of all PQQ treatments were much closer to the negative control (health plants) than the positive control (diseased plants) (Fig. 5D-F). Alike clustering results were found in P50, P100, and P1000 for the SS diseasing patterns, except for P05 (Fig. 5G).

**Fig. 5.**
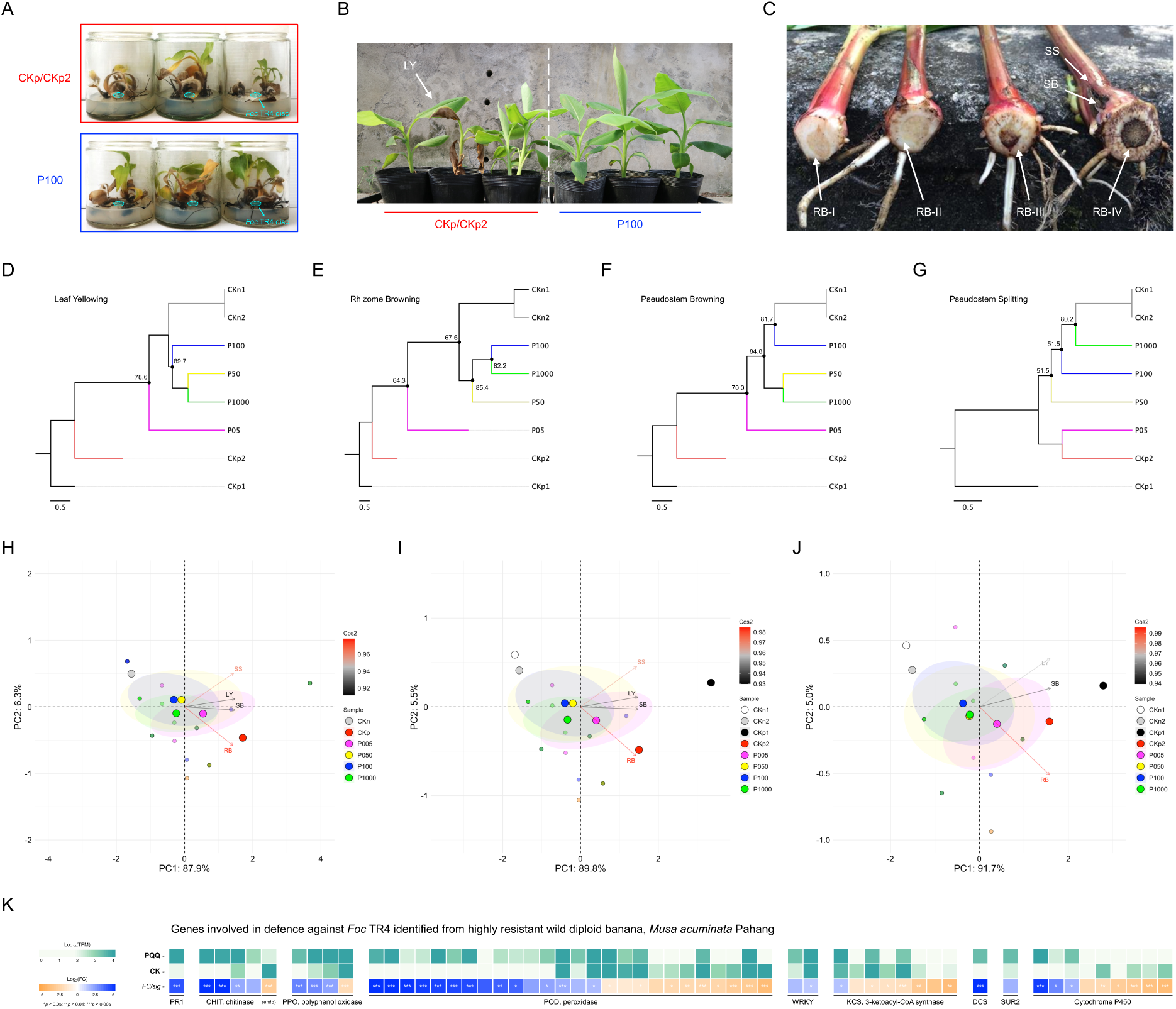
PQQ contributes to the immunity of banana plants combating *Fusarium* wilt. Exogenous supplementation of PQQ contributes to the tolerance of banana plants against *Fusarium* oxysporum f. sp. cubense tropical race 4 (*Foc* TR4) in **A** tissue culture and in **B** soil conditions. *Foc* TR4 infection was performed through **A** discs (1.0 x 10^6^ CFU/mL) placed at the centre of three replicates, highlighted by cyan dashed circles, or through **B** transplantation using infected soil (2.8 x 10^3^ spore/gram). CKp/CKp2 and P100 are samples treated by ddH_2_O and 100 nM PQQ, respectively. White arrows highlight four phenotypic indicators: **B** leaf yellowing (LY), **C** pseudostem splitting (SS), pseudostem (SB) and rhizome dark brown discolouration (RB) for the Panama disease resistance estimation; roman numerals suffixes of RB indicate the levels of disease severity from low to high as described in Materials and Methods. **D-F** Principal component analysis (PCA) on Panama disease indicators (LY, RB, SB and SS) of non-infected (CKn = CKn2; non-PQQ-pretreated) and Foc TR4-infected banana plants pretreated with different concentrations of PQQ; 0 mM (CKp = CKp2), 5 nM (P05), 50 nM (P50), 100 nM (P100) and 1000 nM (P1000); CKn1 and CKp1 are virtual negative and positive controls, respectively, included to assist the analyses; see Material and Methods for further details. Biplot vectors are the disease indicator loadings, whereas the position of each treatment is shown. Cos2 represents the quality of representation for variables on the factor map. **G-J** Unweighted pair group method with arithmetic mean (UPGMA) clustering dendrograms on Panama disease indicators of banana plants treated with different concentrations of PQQ; the dataset was accessed from **E** The trees were constructed based on the Euclidean distancing method with 1,000 multiscale bootstrap re-samplings; the approximately unbiased p-value of those internal nodes with < 90% were labelled. **K** Heatmaps showed the specific gene expression of genes involved in the immunity of wild bananas combating *Fusarium*. Details of gene symbols can be accessed in Table S8. The colour keys refer to the transcripts per kilobase million (TPM) and the corresponding fold changes (FC). Statistical significances between samples were indicated by asterisks (*p<0.05; **p<0.01; ***p<0.005); otherwise, not significant.

A follow-up PCA suggested a similar result to the UPGMA that the PQQ treatments, especially P50 and P100, help to protect bananas against *Foc* TR4 to stand healthier like those non-infected plants ((Fig. 5H-J). Notably, in addition to those DEGs well-known in phytoalexin accumulation and the plant’s systemic immunity induction (Fig. 4D), genes encoding pathogenesis-related protein 1, chitinase, polyphenol oxidase, putative glucuronosyltransferase PGSIP8, 3-ketoacyl-CoA synthase, WRKY transcription factor and disease resistance protein RGA2, were found to be highly regulated (Fig. 5K) (Table S8). These genes have been reported to be highly expressed and critical in the immunity of wild bananas against *Fusarium* wilt (40). Collectively, exogenous supplementation of 100 nM PQQ in the early growing stage not only promotes the organ development of banana plantlets but also triggers the plant’s systemic immunity, protecting bananas against *Fusarium* wilt either in tissue culture or in soil. These results suggested that the exogenous supplementation of PQQ to bananas in their early growth stage could contribute to triggering plant immune systems similar to wild bananas to combat *Fusarium* wilt, providing an untapped opportunity for in-field disease maintenance of the banana industry.

## Discussion

### Molecular roles of endophytic PQQ in plant-microbe mutualisms

On account of being a redox cofactor to types of dehydrogenases, PQQ participates in multiple essential metabolisms across kingdoms. Several studies have reported its effects on either prokaryotic or eukaryotic systems: acting as a strong antioxidant to scavenge reactive oxygen species (41), regulating the balance of nicotinamide adenine dinucleotide (9), inducing mitochondrial biogenesis (10, 11), driving the oxidative phosphorylation and ATP synthesis (12, 13), etc. However, unlike microbes and animals, its molecular roles *in planta* remains limited.

PQQ has been found to be produced by microbes, prevalent by Gram-negative bacteria (42), and benefits the growth of some plants in different stages (16, 17, 43). In this study, the PQQ exogenous supplementation on bananas and the follow-up transcriptomics (Figs. 3,4) provided new materials and insights into its molecular effects in plants. Our findings suggested that PQQ could affect mitochondrial and energetic metabolisms to improve the overall growth of bananas (Figs. S7-S10), which echoed previous works on animal cells. In addition, multiple phytohormone signalling pathways and photosynthesis of banana plants were also found to be coordinated (Figs. 4,S17). These results provided clues about the molecular roles of PQQ in plants and revealed the molecular roles of endophytic PQQ in mutualistic relationships between plants and microbes.

On the other hand, research showed that the disruption of *pqq* genes causes the loss of microbial antifungal activities (20, 21). Han et al. also determined the expression change of a few genes participating in plant immunity, proposing the putative involvement of PQQ in molecular plant-microbe-pathogen tripartite interactions (20). Herein, we confirmed that PQQ has no anti-fungal activity *in vitro* (Fig. 2C) but induces plant immunity *in planta* to protect banana plants against *Fusarium* wilt (Figs. 4-6,S17-S19). Interestingly, the seven immune-related genes with continuously high expression levels of a commercial *Fusarium*-resilient banana cultivar (40) were also found to be highly expressed within banana plants after being treated with 100 nM PQQ (Fig. 5K) (Table S8). This result highlights the potential of endophytic biostimulants, as PQQ here, in breeding biofortification or resistance that achieves similar aims but via fewer costs, deserving more attention for future agriculture.

**Fig. 6.**
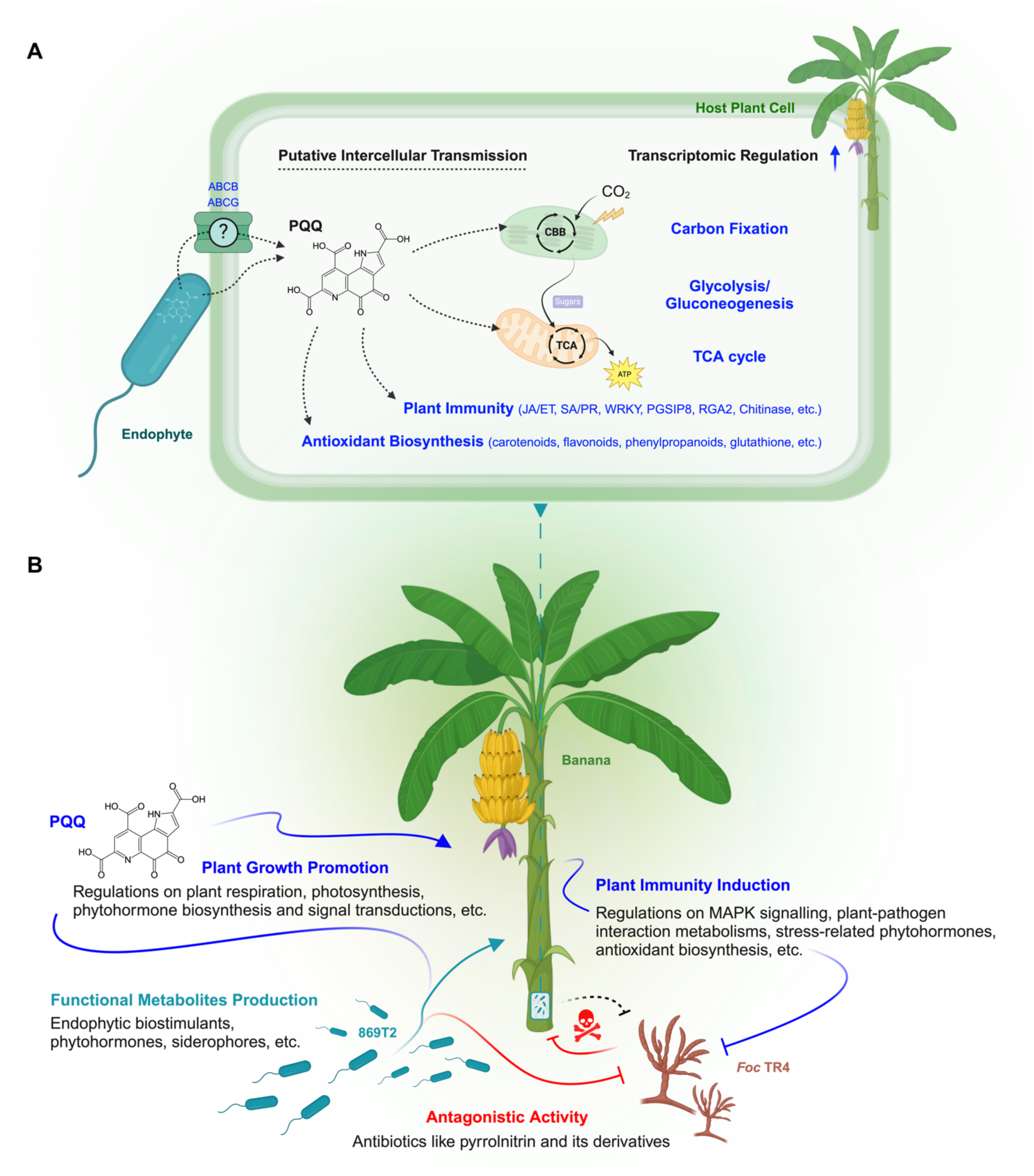
Schematic illustration of plant-endophyte interactions that PQQ confers endophytic niches by coordinating host development. **A** Proposed PQQ molecular mechanisms in plant cells. Dashed lines indicate putative transmission from prokaryotic cells to or within eukaryotic cells. The mainly regulated genes and metabolisms identified with transcriptomics were highlighted in blue. **B** An overall viewpoint on how endophytes benefit the health of host plants and combat the *Fusarium* wilt of bananas. Icons were created with BioRender.com.

### Evolutionary and functional divergence of PQQ

Although the chemical mechanisms of PQQ biosynthesis have been well studied (44), the biosynthesis origin and evolution of this highly conserved order of open reading frame within the *pqq* operon remain unclear. Shen et al. determined the distribution of the main components*, i.e., pqqABCDE*, of the *pqq* operon over 126 prokaryotes (42). Similarly, the *pqq* operon was identified from nearly all genomes of *Burkholderia* species, including environmental isolates, pathogens and symbionts (Fig. 1). In this work, further comparative genomics targeting the *pqq* operon was performed with the purpose of promoting our understanding of such unclear evolutionary linkages between the species.

Interestingly, the result showed an environmental isolate *B. cenocepacia* BCMSMB has two copies of the *pqq* operon (*pqqABCDE*, WL90_31170-31190 and *pqqBCDE*, WL90_32390-32405) (Figs. 1B,S20). The incomplete operon was phylogenetically close to *B. cenocepacia*, while the complete one was distinct to both species. Moreover, the complete operon was further found to be located at a predicted gene island (Fig. S20B), suggesting an occurrence of potential evolution events like inter-species horizontal gene transfer. Besides, the *pqq* operon of a *B. seminalis* sub-group (BS0018+BS1628 clade) seemed to get lost during their evolution. Further checking on the amino acid sequences of the operon, a total of nine single amino acid polymorphisms (SAPs) between the species, five in PqqB, two in PqqC and two in PqqE, were identified by multiple sequence alignment (Fig. S21). The phylogenic trees constructed by amino acid sequences also distinguished two species into different clades (Fig. S22). The catalyst within a PqqA/D/E complex is considered the radical beginning of the PQQ biosynthesis, and the products of spontaneous cyclisation after the PqqB-catalysed reaction is the substrate of PqqC in the final step of PQQ biogenesis (44). Our comparative genomics revealed the potential evolutionary events and phylogenic divergences of the operon, especially PqqB, PqqC and PqqE, of virulence and non-virulence *Burkholderia* strains from different hosts and habitats. The results also shed new light on studying the obscure roles of PQQ among molecular host-microbe interactions and their evolution.

Moreover, distinct from participating as a cofactor for lactate dehydrogenase in animal and microbial cells, our transcriptomic result suggested that PQQ mainly involve regulations related to aldehydes and carboxylic acids in plant cells (Fig. 4D) (Table S7). These differences may be due to the metabolic preference of lactate and ethanol metabolisms in plant cells across the evolution and the results of host-symbiotic co-evolution over time. A better understanding of such evolutionary and functional divergence among lives is believed to provide great potential for academia and industry towards sustainable agriculture. Beyond, some intriguing topics remain for future research, e.g., why and how do non-PQQ-producing organisms, especially in eukaryotic cells like plants, utilise PQQ? To better understand these questions, comprehensive investigations using integrated approaches among PQQ participating host-microbe interactions are necessary for follow-up research.

## Conclusion

Previous studies showed the versatile benefits of endophytes to plants in promoting host health and stress resistance (45–48). This work used multi-omics approaches and imaging mass technologies to reveal a PQQ-involving arm racing among molecular plant-endophyte-pathogen interactions. In summary, we demonstrated that *B. seminalis* 869T2 produces multiple biostimulants (*e.g.*, PQQ and cyclopeptides) and phytohormones (e.g., auxins, cytokinins, gibberellins, etc.) to improve the growth of host plants for symbiotic relationships in general conditions; however, when host plants suffer from pathogens, it also secrets antibiotics (*e.g.*, pyrrolnitrin and the derivates) to fight against the attackers (Fig. 2) (Table S4,S5). Meanwhile, the PQQ induces plant immunity to promote the health and resistance of the host plants, thus establishing an endophytic niche in defending those pathogenic invaders (Fig. 6).

In conclusion, this is the first work through the omics landscape to explain how PQQ involved plant-endophyte interactions and contributed to plant immunity. These results facilitate our understanding of the realm and highlight PQQ as a potent biostimulant candidate for future agriculture. Beyond the findings presented here, many intriguing questions remain about PQQ in molecular plant-microbe interactions. Why and how do non-PQQ-producing organisms utilize PQQ, particularly in plant cells and the organelles? Further studies on their evolutionary and functional biology are expected to enhance our understanding of this unique microbial redox cofactor in the future.

## Materials and methods

### Microbial and plant materials

The strain 869T2, *Fusarium oxysporum* f. sp. *cubense* tropical race 4 (*Foc* TR4) and *Arabidopsis thaliana* ecotype Columbia (Col-0) seeds were retrieved from Chieh-Chen Huang Lab’s stocks. The plantlets of banana (*Musa acuminata* Cavendish cv. Pei-Chiao) were purchased from Taiwan Banana Research Institute (TBRI), Pingtung, Taiwan.

### Bacterial genome sequencing, assembly and annotation

The procedures for genome sequencing and analysis were based on those described in our previous work on bacterial genomes (49–51). The same kits were used following the manufacturer’s protocols, and all bioinformatics tools were used with the default settings unless stated otherwise.

Briefly, the total DNA of strain 869T2 was extracted and sequenced by Illumina MiSeq 2 × 300 bp paired-end (MS-103-1003) and Oxford Nanopore Technologies (ONT) MinION with R9.4 flow cell (FLO-MIN106) followed by Guppy v3.4.5 for basecalling. A hybrid *de novo* assembly was produced using Unicycler v0.4.9-beta (52). For validation, the Illumina and ONT raw reads were mapped to the assembly using BWA v0.7.12 (53) and Minimap2 v2.15 (54). The results were programmatically checked using SAMtools v1.2 (55) and manually inspected using IGV v2.3.57 (56). For assembly completeness evaluation, the benchmarking universal single-copy orthologous (BUSCO) analysis with the Burkholderiales dataset as a reference was executed on gVolante (57). The finalised assembly was submitted to the National Center for Biotechnology Information (NCBI) and annotated using their Prokaryotic Genome Annotation Pipeline (PGAP) (58). The KofamKOALA (59) was further used to examine putative metabolic pathways.

### Bacterial comparative genomics

The genome sequences of *Burkholderia* strains examined by Wallner *et al.* (36), including closely related strains of *B. seminalis* and *B. cenocepacia*, plus an endophyte strain *Paraburkholderia phytofirmans* PsJN (60), were retrieved from NCBI GenBank (61) (accessed on 23 Jul 2021) for comparative analyses with 869T2 (Table S2). FastANI v1.124 (62) was used for whole-genome comparison to calculate the proportion of genomic segments mapped and the average nucleotide identity (ANI). The homologous gene clusters were identified using OrthoMCL v1.3 (63) for maximum likelihood phylogeny construction and gene content analysis. These genes were used to produce a concatenated alignment using MUSCLE v3.8.31 (64) for maximum-likelihood inference by PhyML v.3.3.20180214 (65) and visualised using FigTree v1.4.4. PHYLIP v3.69723 (66) was used for bootstrap analysis. For gene content analysis, the previously reported human virulence-facilitating (VIR) and environmental adaptation and MPMI (ENV) related genes were mainly targeted (36). *B. cenocepacia* J2315^T^ and *B. seminalis* LMG24067^T^ were set as the default reference for the VIR and ENV checks, respectively. Given several incomplete genomes were included and the genetic complexity of the *Burkholderia cepacia* complex (BCC), besides homologous gene clusters inferred by OrthoMCL, an additional manual check on the gene length and annotation was also carried out with BLASTP v2.6.0 (67) with e-value cut-off set to 1e-15 and the amino acid sequence identity cut-off set to 80%. Lists of targeted genes for gene content analysis were provided as supporting information (Table S3).

### Imaging mass spectrometry analysis

Strain 869T2 was first incubated in LB media at 28°C for 24 hours and then transplanted 2 μL of bacterial culture to a PDA plate as a spot. The spot was incubated alone, or a 4 mm mycelial plug of *Foc* TR4 from a 7-day fungal plate was placed at the opposite edge of the bacterial spot for the dual incubation for five days to monitor the metabolic changes by using matrix-assisted laser desorption/ionisation (MALDI) time-of-flight (TOF) imaging mass spectrometry (IMS). All procedures and settings followed the previously described unless otherwise stated (68, 69). Briefly, the sample areas of agar containing the bacteria alone or with fungal mycelia were excised and transferred onto the MALDI stainless steel target plate. A 1:1 mixture of α-cyano-4-hydroxycinnamic acid and 2,5-dihydroxybenzoic acid (universal matrix, Sigma-Aldrich) was spread on top of the samples, then exposed in a 37°C incubator overnight until it was deemed dried. The IMS data were collected using a Bruker Autoflex Speed MALDI-TOF/TOF mass spectrometer (Bruker Daltonik GmbH, Bremen, Germany) with the following setting: 1000 shots/raster, 19.0 and 16.6 kV for the ion sources 1 and 2, 8.7 kV for the lens, 21 and 9.4 kV for reflectors 1 and 2, respectively. The mass ranges from m/z 100-2000 with 25% laser power, positive and reflectron mode was selected, and the raster was set to 800 μm. The universal matrix and peptide standard I (Bruker Daltonik GmbH, Bremen, Germany) were used for MS calibration. All IMS data were processed with TIC normalisation mode and analysed using Fleximaging 3.3 (Bruker Daltonik GmbH, Bremen, Germany).

### *In vitro* antagonism assay

The *in vitro* anti-*Fusarium* activity was assessed by comparing fungal hyphal growth rates under the presence and absence of the examined targets. All procedures followed previous work (28) with minor modifications. Strain 869T2 was set as the positive control, and the ddH_2_O was used as the mocking negative control. Three concentrations of PQQ (10^2^, 10^4^ and 10^6^ nM) were examined. The antagonistic plate images were photographed, the fungal hyphal growth was measured through ImageJ (70), and the antifungal efficiency (AE) was calculated by AE = (DC-DT)/DC x 100%. DC and DT are *Foc* TR4 colony diameters of the negative control and the comparing target, respectively.

### Exogenous supplementation of pyrroloquinoline quinone to plants

The banana plantlets were grown with MS media in the NCHU plant growth chamber as previously described (28) with synthetic PQQ supplementation in different concentrations (Experiment I: 0, 5, 50, 100 and 1000 nM; Experiment II: 0, 100, 1000 and 2000 nM). Phenotypes were photographed, and the growth indicators of visible root formation rate, average leaves number and distribution, plant height, fresh weight, etc., were recorded to determine the plant-growth-promoting (PGP) effects and efficient working dosage of PQQ in plants. Besides, the PQQ-treated banana plantlets were normalised and then transplanted into soil pots, moved to the NCHU greenhouse as described above, and then used for the following *Fusarium* infection assay.

### *Fusarium* infection assay and Panama disease resistance evaluation

All procedures of the infection assay carried out in tissue culture containers or soil pots were followed in previous works (28) unless otherwise stated. *Foc* TR4 infectious inocula were prepared in a final concentration of 10^6^ conidia per mL. For the tissue culture system, a 5 mm diameter filter paper disc soaked within the infectious inocula was placed at the centre of the banana plantlets for the infection. For the soil pot system, the bananas were domesticated in the NCHU greenhouse and then infected by transplantation into pots containing *Foc* TR4, with a final concentration of 2.8 x 10^3^ spores per gram of soil. The planting matrixes prepared with and without *Foc* TR4 were set as the positive (CKp = CKp2) and negative control (CKn = CKn2), respectively. Four indicators of Panama disease, *i.e.*, leaf yellowing (LY), pseudostem splitting (SS), rhizome (RB) and pseudostem dark brown discolouration (SB) (71), and plant morbidity were determined three months after the infection. Plants without and with LY/SS/SB were recorded as 0 and 1, respectively; plants with RB were recorded as 0, 1, 2 and 3 according to the severity of none, low, medium and high (72); dead plants were weighted by adding 1 to the total score. The fully-scored virtual positive control (CKp1) and zero-scored virtual negative control (CKn1) were also included to assist the follow-up analyses.

The principal component analysis (PCA) and the unweighted pair group method with arithmetic mean (UPGMA) were used to analyse the above score data to evaluate the contribution of exogenous supplementation of PQQ to Panama disease resistance. The statistical analyses were performed in R version 4.3.2 (73). For PCA, the function prcomp() was used for primary analysis, and the result was plotted by using the function fviz_pca_biplot(). For UPGMA, the function pvclust() with the distancing method of “Euclidean”, hierarchical clustering method of “average”, and 1,000 times multiscale bootstrap re-samplings were performed. FigTree v1.4.4 was used for the tree visualisation. To test the effect of the clustering method, we also used the function dist() for calculating the Euclidean distancing-based similarity matrix calculation, then used the function upgma() to construct the dendrogram. As both alternatives produced the same clustering suggestion, only the result based on the pvclust() method with multiscale bootstrap analysis was reported.

### Plant total RNA extraction and transcriptomic analysis

The RNA sequencing and analysis procedures were based on those presented in our previous works on plant transcriptomics (50, 72, 74). All kits were used according to the manufacturer’s protocols, and all bioinformatics tools were used with the default settings and followed Welgene Biotech’s in-house pipeline unless stated otherwise. Banana plantlets were harvested after 11 weeks and grown in ½ MS media with the 100 nM PQQ (treatment) and without PQQ (control), respectively. Total RNA was extracted from the mixtures of three bananas for each treatment using Trizol reagent. The library constructions were generated using TruSeq Stranded mRNA Library Prep Kit (RS-122-2101; Illumina, San Diego, CA USA) and assessed to Agilent Bioanalyzer 2100 system using DNA High Sensitivity Chips for qualification. Illumina Hiseq 4000 platform was used for 150-bp paired-end sequencing. Illumina’s program bcl2fastq v2.20 was used for basecalling, and Trimmomatic v0.3642 (75) was implemented to trim off low-quality reads based on Q20 accuracy. The resultant was mapped to *Musa acuminata* subsp. *malaccensis* (NCBI: txid214687) genome (76) using HISAT2 (77). The transcript per million (TPM) method was used for normalisation; genes with low expression levels (< 0.3 TPM value) were excluded. Genes with at least 2.0-fold differences and a probability of at least 0.95 were defined as differentially expressed genes (DEGs). StringTie v2.1.4 (78) and DEseq2 v1.28.1 (79) were used for genome bias detection/correction. Gene set enrichment analysis (GSEA) (80), Gene Ontology (GO) (81) and Kyoto Encyclopedia of Genes and Genomes (KEGG) databases (82) were accessed for the metabolic analyses.

## Statistics

The Shapiro-Wilk test and Levene’s test were used to check the normal distribution of variables and homoscedasticity, respectively. One-way ANOVA analysis with Tukey’s post hoc HSD test for parametric analysis or Kruskal-Wallis test with Dunn’s post hoc test for non-parametric analysis was performed to estimate the statistical significance between samples. The data analysis was performed using the Real Statistics Resource Pack v7.7.1 (83). At least three independent biological replicates were tested for all experiments, and data are mainly shown as mean ± SE. \**P* ≤ 0.05, \*\**P* ≤ 0.01, \*\*\**P* ≤ 0.005 and \*\*\*\**P* ≤ 0.001 indicated significant differences between samples; otherwise, no significance.

## Acknowledgements

The Illumina MiSeq and ONT MinION sequencing services for bacterial genome was provided by the Genomic Technology Core (Institute of Plant and Microbial Biology, Academia Sinica). The RNA-seq services were provided by Welgene Biotech Co., Ltd. (Taipei, Taiwan) and Genomics BioSci & Tech Co., Ltd. (Taipei, Taiwan). We thank the technical assistance and the support from Taiwan Banana Research Institute (TBRI), Pingtung, Taiwan and Taichung District Agricultural Research and Extension Station, Ministry of Agriculture of Taiwan.

## Author Contributions

SHWH, MYY and CCH designed research; CCH and CHK acquired funding; CCH supervised the project; SHWH, MYY, CHL, TCH, JHP and YNH performed research; SHWH, CHL, TCH, JHP and YNH analysed data; SHWH wrote the paper – original draft; SHWH, JHP, NYJ, CHK, YNH, EPIC, HHH and CCH wrote the paper – review & editing.

## Data availability

The complete genome sequence of *Burkholderia seminalis* 869T2 has been deposited in GenBank under the accession numbers CP072520-22. The bacterial genome and the plant transcriptome sequencing project, including the associated raw reads, have been deposited in the NCBI under BioProject PRJNA243842 and BioProject PRJNA786465, respectively.

## References

1. J. G. Hauge, Kinetics and specificity of glucose dehydrogenase from Bacterium anitratum. Biochimica et Biophysica Acta 45, 263–269 (1960).

2. S. A. Salisbury, H. S. Forrest, W. B. T. Cruse, O. Kennard, A novel coenzyme from bacterial primary alcohol dehydrogenases. Nature 280, 843–844 (1979).

3. L. Chistoserdova, S.-W. Chen, A. Lapidus, M. E. Lidstrom, Methylotrophy in Methylobacterium extorquens AM1 from a genomic point of view. J Bacteriol 185, 2980–2987 (2003).

4. J. A. Duine, The PQQ story. J Biosci Bioeng 88, 231–236 (1999).

5. M. Matsutani, T. Yakushi, Pyrroloquinoline quinone-dependent dehydrogenases of acetic acid bacteria. Appl Microbiol Biotechnol 102, 9531–9540 (2018).

6. M. Naveed, K. Tariq, H. Sadia, H. Ahmad, A. S. Mumtaz, The life history of pyrroloquinoline quinone (pqq): a versatile molecule with novel impacts on living systems. IJMBOA 1, 29–46 (2016).

7. C. Anthony, Pyrroloquinoline quinone (PQQ) and quinoprotein enzymes. Antioxid Redox Signal 3, 757–774 (2001).

8. P. M. Goodwin, C. Anthony, The biochemistry, physiology and genetics of PQQ and PQQ-containing enzymes. Adv Microb Physiol 40, 1–80 (1998).

9. K. R. Jonscher, W. Chowanadisai, R. B. Rucker, Pyrroloquinoline-quinone is more than an antioxidant: a vitamin-like accessory factor important in health and disease prevention. Biomolecules 11, 1441 (2021).

10. W. Chowanadisai, et al., Pyrroloquinoline Quinone Stimulates Mitochondrial Biogenesis through cAMP Response Element-binding Protein Phosphorylation and Increased PGC-1α Expression*. Journal of Biological Chemistry 285, 142–152 (2010).

11. K. Saihara, R. Kamikubo, K. Ikemoto, K. Uchida, M. Akagawa, Pyrroloquinoline Quinone, a Redox-Active o-Quinone, Stimulates Mitochondrial Biogenesis by Activating the SIRT1/PGC-1α Signaling Pathway. Biochemistry 56, 6615–6625 (2017).

12. A. Canovai, et al., Pyrroloquinoline quinone drives ATP synthesis in vitro and in vivo and provides retinal ganglion cell neuroprotection. Acta Neuropathologica Communications 11, 146 (2023).

13. M. C. Ebeling, et al., Improving retinal mitochondrial function as a treatment for age-related macular degeneration. Redox Biology 34, 101552 (2020).

14. R. Carreño-López, J. M. Alatorre-Cruz, V. Marín-Cevada, “Pyrroloquinoline quinone (PQQ): role in plant-microbe interactions” in Secondary Metabolites of Plant Growth Promoting Rhizomicroorganisms: Discovery and Applications, H. B. Singh, C. Keswani, M. S. Reddy, E. Sansinenea, C. García-Estrada, Eds. (Springer, 2019), pp. 169–184.

15. M. Ben Farhat, A. Fourati, H. Chouayekh, Coexpression of the pyrroloquinoline quinone and glucose dehydrogenase genes from Serratia marcescens CTM 50650 conferred high mineral phosphate-solubilizing ability to Escherichia coli. Appl Biochem Biotechnol 170, 1738–1750 (2013).

16. Y.-S. Cho, et al., PQQ-dependent organic acid production and effect on common bean growth by Rhizobium tropici CIAT 899. Journal of Microbiology and Biotechnology 13, 955–959 (2003).

17. O. Choi, et al., Pyrroloquinoline quinone is a plant growth promotion factor produced by Pseudomonas fluorescens B16. Plant Physiol 146, 657–668 (2008).

18. L. Li, Z. Jiao, L. Hale, W. Wu, Y. Guo, Disruption of gene pqqA or pqqB reduces plant growth promotion activity and biocontrol of crown gall disease by Rahnella aquatilis HX2. PLoS One 9, e115010 (2014).

19. A. H. Patel, V. Chovatia, S. Shah, Expression of pyrroloquinoline quinone in Rhizobium leguminosarum for phosphate solubilization. Environment and Ecology 33, 621–624 (2015).

20. S. H. Han, et al., Inactivation of pqq genes of Enterobacter intermedium 60-2G reduces antifungal activity and induction of systemic resistance. FEMS Microbiol Lett 282, 140– 146 (2008).

21. J. Xu, et al., The pqqC gene is essential for antifungal activity of Pseudomonas kilonensis JX22 against Fusarium oxysporum f. sp. lycopersici. FEMS Microbiology Letters 353, 98–105 (2014).

22. F. Garcia-Bastidas, Fusarium oxysporum f.sp. cubense tropical race 4 (Foc TR4). CABI Compendium **CABI Compendium**, 59074053 (2022).

23. R. C. Ploetz, Fusarium Wilt of Banana. Phytopathology® 105, 1512–1521 (2015).

24. TR4 Global Network, TR4 Basics. Available at https://www.fao.org/tr4gn/tr4-basics/en/. Deposited 15 January 2024.

25. A. A. Ismaila, et al., Fusarium wilt of banana: Current update and sustainable disease control using classical and essential oils approaches. Horticultural Plant Journal 9, 1– 28 (2023).

26. G. Bubici, M. Kaushal, M. I. Prigigallo, C. Gómez-Lama Cabanás, J. Mercado-Blanco, Biological Control Agents Against Fusarium Wilt of Banana. Frontiers in Microbiology 10, 616 (2019).

27. C. Du, et al., Construction of a compound microbial agent for biocontrol against Fusarium wilt of banana. Frontiers in Microbiology 13, 1066807 (2022).

28. Y.-N. Ho, et al., In planta biocontrol of soilborne Fusarium wilt of banana through a plant endophytic bacterium, Burkholderia cenocepacia 869T2. Plant Soil 387, 295–306 (2015).

29. Z. Zhu, et al., Spatiotemporal biocontrol and rhizosphere microbiome analysis of Fusarium wilt of banana. Commun Biol 6, 1–10 (2023).

30. T. Damodaran, et al., Secondary metabolite induced tolerance to Fusarium oxysporum f.sp. cubense TR4 in banana cv. Grand Naine through in vitro bio-immunization: a prospective research translation from induction to field tolerance. Frontiers in Microbiology 14, 1233469 (2023).

31. L. Zhang, et al., Biocontrol Potential of Endophytic Streptomyces malaysiensis 8ZJF-21 From Medicinal Plant Against Banana Fusarium Wilt Caused by Fusarium oxysporum f. sp. cubense Tropical Race 4. Frontiers in Plant Science 13, 874819 (2022).

32. Y.-N. Ho, C.-C. Huang, Draft genome sequence of Burkholderia cenocepacia strain 869T2, a plant-beneficial endophytic bacterium. Genome Announcements 3, e01327–15 (2015).

33. H.-H. Hwang, et al., A plant endophytic bacterium, Burkholderia seminalis strain 869T2, promotes plant growth in Arabidopsis, Pak Choi, Chinese amaranth, lettuces, and other vegetables. Microorganisms 9, 1703 (2021).

34. S.-H. W. Hung, et al., Endophytic biostimulants for smart agriculture: Burkholderia seminalis 869T2 benefits heading leafy vegetables in-field management in Taiwan. Agronomy 13, 967 (2023).

35. B.-A. T. Nguyen, et al., Biodegradation of dioxins by Burkholderia cenocepacia strain 869T2: role of 2-haloacid dehalogenase. Journal of Hazardous Materials 401, 123347 (2021).

36. A. Wallner, E. King, E. L. M. Ngonkeu, L. Moulin, G. Béna, Genomic analyses of Burkholderia cenocepacia reveal multiple species with differential host-adaptation to plants and humans. BMC Genomics 20, 803 (2019).

37. A. Baldwin, P. A. Sokol, J. Parkhill, E. Mahenthiralingam, The Burkholderia cepacia epidemic strain marker is part of a novel genomic island encoding both virulence and metabolism-associated genes in Burkholderia cenocepacia. Infect Immun 72, 1537– 1547 (2004).

38. S. Pawar, A. Chaudhari, R. Prabha, R. Shukla, D. P. Singh, Microbial Pyrrolnitrin: Natural Metabolite with Immense Practical Utility. Biomolecules 9, 443 (2019).

39. Y. B. Guo, et al., Mutations That Disrupt Either the pqq or the gdh Gene of Rahnella aquatilis Abolish the Production of an Antibacterial Substance and Result in Reduced Biological Control of Grapevine Crown Gall. Applied and Environmental Microbiology 75, 6792–6803 (2009).

40. L. Zhang, et al., Transcriptomic analysis of resistant and susceptible banana corms in response to infection by Fusarium oxysporum f. sp. cubense tropical race 4. Sci Rep 9, 8199 (2019).

41. H. S. Misra, et al., Pyrroloquinoline-quinone: a reactive oxygen species scavenger in bacteria. FEBS Letters 578, 26–30 (2004).

42. Y.-Q. Shen, et al., Distribution and properties of the genes encoding the biosynthesis of the bacterial cofactor, pyrroloquinoline quinone. Biochemistry 51, 2265–2275 (2012).

43. L. B. Xiong, J. Sekiya, N. Shimose, Occurrence of Pyrroloquinoline Quinone (PQQ) in Pistils and Pollen Grains of Higher Plants. Agricultural and Biological Chemistry 54, 249–250 (1990).

44. Zhu, J. P. Klinman, Biogenesis of the peptide-derived redox cofactor pyrroloquinoline quinone. Current Opinion in Chemical Biology 59, 93–103 (2020).

45. S. Card, L. Johnson, S. Teasdale, J. Caradus, Deciphering endophyte behaviour: the link between endophyte biology and efficacious biological control agents. FEMS Microbiol Ecol 92, fiw114 (2016).

46. P. Chaudhary, U. Agri, A. Chaudhary, A. Kumar, G. Kumar, Endophytes and their potential in biotic stress management and crop production. Frontiers in Microbiology 13, 933017 (2022).

47. J. Hallmann, A. Quadt-Hallmann, W. F. Mahaffee, J. W. Kloepper, Bacterial endophytes in agricultural crops. Can. J. Microbiol. 43, 895–914 (1997).

48. P. R. Hardoim, et al., The hidden world within plants: ecological and evolutionary considerations for defining functioning of microbial endophytes. Microbiol Mol Biol Rev 79, 293–320 (2015).

49. S.-H. W. Hung, M.-C. Chiu, C.-C. Huang, C.-H. Kuo, Complete genome sequence of Curtobacterium sp. C1, a beneficial endophyte with the potential for in-plant salinity stress alleviation. MPMI 35, 731–735 (2022).

50. S.-H. W. Hung, et al., A cyclic dipeptide for salinity stress alleviation and the trophic flexibility of endophyte provide insights into saltmarsh plant–microbe interactions. ISME Communications 4, ycae041 (2024).

51. S.-H. W. Hung, I.-C. Wu, C.-C. Huang, C.-H. Kuo, Complete genome sequence of Erythrobacteraceae bacterium WH01K, a strain isolated from corals (Acropora sp.) in Taiwan. Microbiology Resource Announcements 12, e00830–23 (2023).

52. R. R. Wick, L. M. Judd, C. L. Gorrie, K. E. Holt, Unicycler: resolving bacterial genome assemblies from short and long sequencing reads. PLoS Comput Biol 13, e1005595 (2017).

53. H. Li, R. Durbin, Fast and accurate short read alignment with Burrows-Wheeler transform. Bioinformatics 25, 1754–1760 (2009).

54. H. Li, Minimap2: pairwise alignment for nucleotide sequences. Bioinformatics 34, 3094–3100 (2018).

55. H. Li, et al., The sequence alignment/map format and SAMtools. Bioinformatics 25, 2078–2079 (2009).

56. J. T. Robinson, et al., Integrative genomics viewer. Nat Biotechnol 29, 24–26 (2011).

57. O. Nishimura, Y. Hara, S. Kuraku, Evaluating genome assemblies and gene models using gVolante. Methods Mol Biol 1962, 247–256 (2019).

58. T. Tatusova, et al., NCBI prokaryotic genome annotation pipeline. Nucleic Acids Res 44, 6614–6624 (2016).

59. T. Aramaki, et al., KofamKOALA: KEGG Ortholog assignment based on profile HMM and adaptive score threshold. Bioinformatics 36, 2251–2252 (2020).

60. A. Weilharter, et al., Complete genome sequence of the plant growth-promoting endophyte Burkholderia phytofirmans strain PsJN. Journal of Bacteriology 193, 3383– 3384 (2011).

61. D. A. Benson, et al., GenBank. Nucleic Acids Res 46, D41–D47 (2018).

62. C. Jain, L. M. Rodriguez-R, A. M. Phillippy, K. T. Konstantinidis, S. Aluru, High throughput ANI analysis of 90K prokaryotic genomes reveals clear species boundaries. Nat Commun 9, 5114 (2018).

63. L. Li, C. J. Stoeckert, D. S. Roos, OrthoMCL: identification of ortholog groups for eukaryotic genomes. Genome Res. 13, 2178–2189 (2003).

64. R. C. Edgar, MUSCLE: multiple sequence alignment with high accuracy and high throughput. Nucleic Acids Research 32, 1792–1797 (2004).

65. S. Guindon, O. Gascuel, A simple, fast, and accurate algorithm to estimate large phylogenies by maximum likelihood. Systematic Biology 52, 696–704 (2003).

66. J. Felsenstein, PHYLIP – phylogeny inference package (version 3.2). Cladistics 5, 164–166 (1989).

67. C. Camacho, et al., BLAST+: architecture and applications. BMC Bioinformatics 10, 421 (2009).

68. Y.-N. Ho, et al., Specific inactivation of an antifungal bacterial siderophore by a fungal plant pathogen. ISME J 15, 1858–1861 (2021).

69. Y.-L. Yang, Y. Xu, P. Straight, P. C. Dorrestein, Translating metabolic exchange with imaging mass spectrometry. Nat Chem Biol 5, 885–887 (2009).

70. C. A. Schneider, W. S. Rasband, K. W. Eliceiri, NIH image to ImageJ: 25 years of image analysis. Nat Methods 9, 671–675 (2012).

71. Y. Wu, et al., Systemic acquired resistance in Cavendish banana induced by infection with an incompatible strain of Fusarium oxysporum f. sp. cubense. Journal of Plant Physiology 170, 1039–1046 (2013).

72. M.-Y. Yu, “A study of pyrroloquinoline quinone on Pei-Chiao (Musa spp.) growth and biocontrol improvement,” National Chung Hsing University, Taichung, Taiwan. (2019).

73. R Core Team, R: A Language and Environment for Statistical Computing. (2024). Deposited 18 March 2024.

74. Y.-H. Tsai, “Transcriptomic analysis on adaptive immunity of Pei-Chiao (Musa spp., Cavendish AAA) against Fusarium wilt caused by Fusarium oxysporum f. sp. cubense tropical race4.,” National Chung Hsing University, Taichung, Taiwan. (2019).

75. A. M. Bolger, M. Lohse, B. Usadel, Trimmomatic: a flexible trimmer for Illumina sequence data. Bioinformatics 30, 2114–2120 (2014).

76. A. D’Hont, et al., The banana (Musa acuminata) genome and the evolution of monocotyledonous plants. Nature 488, 213–217 (2012).

77. D. Kim, B. Langmead, S. L. Salzberg, HISAT: a fast spliced aligner with low memory requirements. Nat Methods 12, 357–360 (2015).

78. M. Pertea, et al., StringTie enables improved reconstruction of a transcriptome from RNA-seq reads. Nat Biotechnol 33, 290–295 (2015).

79. M. I. Love, W. Huber, S. Anders, Moderated estimation of fold change and dispersion for RNA-seq data with DESeq2. Genome Biology 15, 550 (2014).

80. A. Subramanian, et al., Gene set enrichment analysis: A knowledge-based approach for interpreting genome-wide expression profiles. Proc. Natl. Acad. Sci. U.S.A. 102, 15545–15550 (2005).

81. H. Mi, A. Muruganujan, D. Ebert, X. Huang, P. D. Thomas, PANTHER version 14: more genomes, a new PANTHER GO-slim and improvements in enrichment analysis tools. Nucleic Acids Research 47, D419–D426 (2019).

82. M. Kanehisa, S. Goto, KEGG: Kyoto encyclopedia of genes and genomes. Nucleic Acids Res 28, 27–30 (2000).

83. C. Zaiontz, Real Statistics Resource Pack software. Available at www.real-statistics.com. Deposited 21 February 2024.

