## Supporting Tables for "Endophytic pyrroloquinoline quinone enhances banana growth and immunity against *Fusarium* wilt for plant-microbe mutualisms"

**Running Title:** PQQ coordinates plant growth and immunity

Shih-Hsun Walter Hung[1,2,3,4], Man-Yun Yu[1], Chia-Ho Liu[1], Tsai-Ching Huang[1], Jian-Hau Peng[5,6], Nai-Yun Jang[1], Chih-Horng Kuo[2,5,6,7], Ying-Ning Ho[8,9,10], En-Pei Isabel Chiang[3,4,5,6,11], Hau-Hsuan Hwang[1,3,4,5,6], and Chieh-Chen Huang[1,3,4,5,6]*

[1] Department of Life Sciences, National Chung Hsing University, Taichung 402202, Taiwan; [2] Institute of Plant and Microbial Biology, Academia Sinica, Taipei 115201, Taiwan; [3] Advanced Plant and Food Crop Biotechnology Center, National Chung Hsing University, Taichung 402202, Taiwan; [4] Innovation and Development Centre of Sustainable Agriculture, National Chung Hsing University, Taichung 402202, Taiwan; [5] PhD Program in Microbial Genomics, National Chung Hsing University, Taichung 402202, Taiwan; [6] PhD Program in Microbial Genomics, Academia Sinica, Taipei 115201, Taiwan; [7] Biotechnology Center, National Chung Hsing University, Taichung 402202, Taiwan; [8] Institute of Marine Biology, College of Life Science, National Taiwan Ocean University, Keelung 202301, Taiwan; [9] Center of Excellence for the Oceans, National Taiwan Ocean University, Keelung 202301, Taiwan; [10] Taiwan Ocean Genome Center, National Taiwan Ocean University, Keelung 202301, Taiwan; [11] Department of Food Science and Biotechnology, National Chung Hsing University, Taichung 402202, Taiwan

**ORCID** SHW Hung, 0000-0002-8749-5653; CH Liu, 0000-0001-9166-3966; TC Huang, 0000-0002-3410-2051; JH Peng, 0000-0001-6052-8606; CH Kuo, 0000-0002-2857-0529; YN Ho, 0000-0003-0943-1416; EPI Chiang, 0000-0002-0158-0962; HH Hwang, 0000-0001-9132-0242; CC Huang, 0000-0002-3739-6315.

**Keywords:** endophyte, symbiosis, pyrroloquinoline quinone (PQQ), Fusarium wilt of banana, molecular plant-microbe interactions (MPMI)

**Supplementary Tables**

**Table S1 Genomic features of *Burkholderia seminalis* 869T2**

**Table S2 List of analysed strains**

**Table S3 Lists of targeted genes for gene content analysis**

**Table S4 List of endophytic biostimulants identified from the metabolome of *B. seminalis* 869T2**

**Table S5 List of cyclic dipeptides identified from the metabolome of *B. seminalis* 869T2**

**Table S6 Profiling result of GO analysis**

**Table S7 List of gene symbols used in transcriptomic analysis**

**Table S8 List of genes associated with the immunity of wild bananas against *Fusarium* wilt**

### Supplementary Tables

#### Table S1 Genomic features of *Burkholderia seminalis* 869T2

| Features | Chromosome1 | Chromosome2 | Chromosome3 |
|---|---|---|---|
| **Accession** | NZ_CP072520.1 | NZ_CP072521.1 | NZ_CP072522.1 |
| **Size (bp)** | 3,561,262 | 3,012,498 | 1,448,576 |
| **GC content (%)** | 67 | 67.5 | 66 |
| **Protein-coding genes** | 3214 | 2654 | 1222 |
| **Pseudogenes** | 25 | 39 | 48 |
| **Ribosomal RNA genes** | 12 | 3 | 3 |
| **Transfer RNA genes** | 60 | 5 | 2 |
| **Other noncoding RNA genes** | 4 | 0 | 0 |

#### Table S2 List of analysed strains

| Strain[a] | Accession | Assembly | Genome size (Mb) | GC (%) | CDS | Source | Note |
|---|---|---|---|---|---|---|---|
| *Burkholderia seminalis* 869T2 | GCA_000705535.2 | Complete: 3 | 8.02 | 67.05 | 7048 | plant | endophyte; this study |
| *B. seminalis* FL-5-5-10-S1-D0 | GCA_001524085.1 | Contig: 55 | 7.55 | 67.40 | 6693 | soil | na |
| *B. seminalis* FL-5-4-10-S1-D7 | GCA_001718535.1 | Complete: 3 | 7.65 | 67.27 | 6747 | soil | na |
| *B. seminalis* TC3.4.2R3 | GCA_001742345.1 | Contig: 84 | 7.67 | 67.20 | 6827 | plant | endophyte |
| *B. seminalis* Bp9022 | GCA_003854315.1 | Contig: 72 | 7.64 | 67.30 | 6787 | water | na |
| *B. seminalis* Bp8988 | GCA_003854335.1 | Contig: 82 | 7.85 | 67.20 | 7032 | soil | na |
| *B. seminalis* BC00027 | GCA_016505695.1 | Contig: 83 | 7.66 | 67.30 | 6857 | human | no disease |
| *B. seminalis* LMG24067[T] | GCA_902832885.1 | Scaffold: 55 | 7.97 | 67.10 | 7019 | human | cystic fibrosis |
| *B. seminalis* BC00018 | GCA_016508205.1 | Contig: 59 | 7.38 | 67.10 | 6628 | human | no disease |
| *B. seminalis* BCC1628 | GCA_902833035.1 | Scaffold: 115 | 8.24 | 66.70 | 7280 | plant | na |
| *B. cenocepacia* VC7848[b] | GCA_001999785.1 | Complete: 1 | 7.50 | 66.90 | 6647 | human | cystic fibrosis |
| *B. cenocepacia* J2315[T] | GCA_000009485.1 | Complete: 4 | 8.06 | 66.92 | 7201 | human | cystic fibrosis |
| *B. cenocepacia* MSMB384WGS | GCA_001718895.1 | Complete: 3 | 7.78 | 67.24 | 6918 | water | na |
| *B. cenocepacia* K56-2 | GCA_014357995.1 | Complete: 4 | 7.74 | 67.02 | 6851 | human | cystic fibrosis |
| *B. cenocepacia* K56-2Valvano | GCA_000333155.2 | Contig: 19 | 7.75 | 67.00 | 6813 | human | cystic fibrosis |
| *B. cenocepacia* F01 | GCA_900240025.1 | Contig: 91 | 8.03 | 67.20 | 7193 | soil | na |
| *Paraburkholderia phytofirmans* PsJN[T] | GCA_000020125.1 | Complete: 3 | 8.21 | 62.32 | 7177 | plant | endophyte |

[a] Superscripts following the strain names: T = type strain. [b] The strain was suggested to be re-assigned as *Burkholderia* sp. nov. (Wallner et al., 2019).

### Table S3 Lists of targeted genes for gene content analysis

| Category[a] | Gene | Description |
|---|---|---|
| ENV | *pqqABCDE* | pyrroloquinoline quinone operon |
| ENV | *prnABCD* | pyrrolnitrin biosynthesis cluster |
| ENV | *uxa* | altronate dehydratase |
| ENV | *oxd* | phenylacetaldoxime dehydratase |
| ENV | *nthAB* | nitrile hydratase cluster |
| ENV | *iaaM* | tryptophan-2-monooxygenase |
| ENV | *iaaH* | indole-3-acetamide hydrolase |
| ENV | *faeB* | feruloyl esterase |
| ENV | *acdS* | 1-aminocyclopropane-1-carboxylic acid deaminase |
| ENV | *llpA* | lectin like bacteriocin 88 |
| VIR | *kdgR* | transcriptional regulator |
| VIR | *telA* | toxic anion resistance protein |
| VIR | *xsc* | sulfoacetaldehyde acetyltransferase |
| VIR | *tauX* | taurine dehydrogenase |
| VIR | *narIJHG* | respiratory nitrate reductase cluster |
| VIR | *cblBACDSTR* | cable pilus assembly operon |
| VIR | *adhA* | 22 kDa adhesion |
| VIR | *baiE* | bile acid 7-alpha dehydratase |
| VIR | afcE | antifungal activity cluster |
| VIR/BCESM | *opcI* | porin |
| VIR/BCESM | *cciI* | N-acyl homoserine lactone synthase gene |
| VIR/BCESM | *cciR* | transcriptional regulator |
| VIR/BCESM | *esmR* | transcriptional regulator |
| VIR/BCESM | *amiI* | amidase |

[a] Category abbreviation: ENV = environmental adaptation and molecular plant-microbe interactions related genes, VIR = human virulence-facilitating related genes and BCESM = Burkholderia cepacia epidemic strain marker.

**Table S4 List of endophytic biostimulants identified from the metabolome of *B. seminalis* 869T2**

| Category[a] | Name | Description |
|---|---|---|
| EVN | Isopyrrolnitrin | Antifungal; Pyrrolnitrin derivate |
| EVN | oxypyrrolnitrin | Antifungal; Pyrrolnitrin derivate |
| EVN | Fluoropyrrolnitrin | Antifungal; Pyrrolnitrin derivate |
| EVN | 3-chloro-4-(2-nitrophenyl)-1H-pyrrole | Antifungal; Pyrrolnitrin derivate |
| EVN | Flavacid | Antifungal |
| EVN | Indole-3-ethanol | Antifungal |
| EVN | Karnamicin A3, D1 | Antifungal |
| EVN | Aerugine | Antifungal; Antioomycete |
| EVN | N-Salicyloyl-2-aminopropan-1-ol | Antibacterial |
| EVN | N-Salicyloyl-2-aminopropan-1,3-diol | Antibacterial |
| EVN | Indoleacryloisonitrile, Antibiotics B 371 | Antibacterial |
| EVN | 7-Chloro-4a-hydroxy-8-methoxy-N-methyl-tetracycline | Antibacterial |
| EVN | Indole-3-carbaldehyde | Phytoalexin biosynthesis related |
| EVN | Indol-3-carboxylic acid | Phytoalexin biosynthesis related |
| EVN | Naphthoquinone and other antibiotics | Antibiotics |
| MPMI | Pyrroloquinoline quinone | Endophytic biostimulants, redox cofactor |
| MPMI | Indole-3-acetic acid | Auxin, IAA |
| MPMI | Indole-3-acetic acid methyl ester | Auxin, IAA |
| MPMI | Chloroindole | Cytochrome P450 substrate |
| MPMI | Phenylacetic acid | Auxin, PAA |
| MPMI | Benzylcyanide | Nitrilase-mediated PAA conversion |
| MPMI | trans-Zeatin | Cytokinin, Zeatin |
| MPMI | Zeatin-riboside | Cytokinin, Zeatin |
| MPMI | Zeatin-2'-deoxyriboside | Cytokinin, Zeatin |
| MPMI | 1'-Methyl-zeatin | Cytokinin, Zeatin |
| MPMI | Gibbestatin-B | GA-induced α-amylase regulation |
| MPMI | Rhizobitoxin | ACC synthase inhibitor |
| MPMI | Salicylic acid | Salicylic acid, SA |
| MPMI | Pyocyanine, PCN | ISR (PCH/PCN) |
| MPMI | Pyochelin, PCH | Siderophores |
| MPMI | Neopyochelin | Siderophores |
| MPMI | Proferrioxamines | Siderophores |

[a] Category abbreviation: ENV = environmental adaptation and MPMI = molecular plant-microbe interactions related genes.

**Table S5 List of cyclic dipeptides identified from the metabolome of *B. seminalis* 869T2**

| Name | Formula | MW | Description |
|---|---|---|---|
| cyclo(L-Pro, L-Val) | $C_{10}H_{16}N_2O_2$ | 196.246 | Auxin-like activity; Anti-fungal; Quorum sensing |
| cyclo(L-Tyr, D-4OH-Pro) | $C_{14}H_{16}N_2O_4$ | 276.288 | Putative sponge-bacteria interaction |
| cyclo(L-Tyr, L-4OH-Pro) | $C_{14}H_{16}N_2O_4$ | 276.288 | Putative sponge-bacteria interaction |
| cyclo(L-4OH-Pro, D-Leu) | $C_{11}H_{18}N_2O_3$ | 226.272 | Plant growth-promoting factor |
| cyclo(L-Pro, L-Ile) | $C_{11}H_{18}N_2O_2$ | 210.273 | Anti-fungal |
| cyclo(L-Pro, L-Met) | $C_{10}H_{16}N_2O_2S$ | 228.311 | Anti-fungal |
| cyclo(D-Pro, D-Val) | $C_{10}H_{16}N_2O_2$ | 196.246 | Anti-bacterial |
| cyclo(D-Pro, D-Ile) | $C_{11}H_{18}N_2O_2$ | 210.273 | Anti-bacterial |
| cyclo(Asp, Leu) | $C_9H_{14}N_2O_4$ | 214.218 | Anti-bacterial |
| cyclo(D-Phe, D-Pro) | $C_{14}H_{16}N_2O_2$ | 244.289 | Anti-bacterial |
| cyclo(D-Phe, L-Pro) | $C_{14}H_{16}N_2O_2$ | 244.289 | Anti-larval/fouling |
| cyclo(L-Trp, L-Pro) | $C_{16}H_{17}N_3O_2$ | 283.325 | Anti-larval/fouling |
| cyclo(L-Arg, D-Pro) | $C_{11}H_{19}N_5O_2$ | 253.301 | Anti-yeast, chitinase Inhibitors |
| cyclo(L-Phe, L-Pro) | $C_{14}H_{16}N_2O_2$ | 244.289 | Anti-bacterial/larval; Quorum sensing |
| cyclo(L-Tyr, L-Pro) | $C_{14}H_{16}N_2O_3$ | 260.288 | Anti-bacterial/fungal; Quorum sensing |
| cyclo(D-Pro, L-Tyr) | $C_{14}H_{16}N_2O_3$ | 260.288 | Anti-bacterial/fungal; Quorum sensing |
| cyclo(L-Leu, L-Pro) | $C_{11}H_{18}N_2O_2$ | 210.273 | Anti-bacterial/fungal/larval/fouling; Quorum sensing |
| cyclo(D-Leu, D-Pro) | $C_{11}H_{18}N_2O_2$ | 210.273 | Anti-bacterial/fungal/larval/fouling; Quorum sensing |
| cyclo(Leu, Pro) | $C_{11}H_{18}N_2O_2$ | 210.273 | Anti-bacterial/fungal/larval/fouling; Quorum sensing |
| cyclo(Gly, D-Pro) | $C_7H_{10}N_2O_2$ | 154.167 | Anti-neoplastic |
| cyclo(Phe, dhAbu) | $C_{13}H_{14}N_2O_2$ | 230.262 | Quorum sensing |
| cyclo(hexenyl, Gly) | $C_8H_{13}NO_2$ | 155.194 | Amino acid antagonist |
| cyclo(D-Pro, L-Val) | $C_{10}H_{16}N_2O_2$ | 196.246 | Specific b-glucosidase inhibitor |
| cyclo(L-Leu, Gly) | $C_8H_{14}N_2O_2$ | 170.209 | TBD |
| cyclo(Ala, OH-Pro)(S,S,R) | $C_8H_{12}N_2O_3$ | 184.192 | TBD |
| cyclo(Ala, OH-Pro)(S,R,R) | $C_8H_{12}N_2O_3$ | 184.192 | TBD |
| cyclo(D-Pipecolinyl, L-Ile) | $C_{12}H_{20}N_2O_2$ | 224.299 | TBD |
| cyclo(Ala, Trp)(3R,6R) | $C_{14}H_{15}N_3O_2$ | 257.288 | TBD |
| cyclo(D-Tyr, D-Pro) | $C_{14}H_{16}N_2O_3$ | 260.288 | TBD |
| cyclo(N-Me-Tyr, dhAbu) | $C_{14}H_{16}N_2O_3$ | 260.288 | TBD |
| cyclo(L-Ile, L-Pro, L-Leu, L-Pro) | $C_{22}H_{36}N_4O_4$ | 420.546 | TBD |

#### Table S6 Profiling result of GO analysis

| Up-regulation | | | | |
|---|---|---|---|---|
| Overall | GO | Hits | Ratio (%) | |
| | mapped | 184 | 32.17 | |
| | unmapped | 388 | 67.83 | |
| Top 10 MF Terms | Molecular Function (MF), total 130 hits | | | |
| | Enrichment | Count | Ratio (%) | Elements |
| | 1 | 27 | 8.13 | metal ion binding |
| | 2 | 20 | 6.02 | transferase activity |
| | 3 | 19 | 5.72 | oxidoreductase activity |
| | 3 | 19 | 5.72 | DNA binding |
| | 5 | 15 | 4.52 | nucleotide binding |
| | 6 | 11 | 3.31 | ATP binding |
| | 7 | 10 | 3.01 | catalytic activity |
| | 7 | 10 | 3.01 | hydrolase activity |
| | 9 | 8 | 2.41 | DNA-binding transcription factor activity |
| | 10 | 6 | 1.81 | lyase activity |
| | 10 | 6 | 1.81 | methyltransferase activity |
| Top 10 BP Terms | Biological Process (BP), total 115 hits | | | |
| | Enrichment | Count | Ratio (%) | Elements |
| | 1 | 14 | 6.9 | regulation of DNA-templated transcription |
| | 2 | 8 | 3.9 | transmembrane transport |
| | 3 | 6 | 3.0 | metabolic process |
| | 4 | 5 | 2.5 | response to wounding |
| | 4 | 5 | 2.5 | defense response |
| | 4 | 5 | 2.5 | phosphorylation |
| | 4 | 5 | 2.5 | phenylpropanoid biosynthetic process |
| | 4 | 5 | 2.5 | methylation |
| | 9 | 4 | 2.0 | regulation of defense response |
| | 9 | 4 | 2.0 | regulation of jasmonic acid mediated signaling pathway |
| | 9 | 4 | 2.0 | response to biotic stimulus |
| | 9 | 4 | 2.0 | monoatomic ion transport |
| | 9 | 4 | 2.0 | carbohydrate metabolic process |
| Top 10 BP Terms | Cellular Component (CC), total 23 hits | | | |
| | Enrichment | Count | Ratio (%) | Elements |
| | 1 | 42 | 38.2 | membrane |
| | 2 | 18 | 16.4 | nucleus |

| | | | | |
|---|---|---|---|---|
| | 3 | 12 | 10.9 | extracellular region |
| | 4 | 6 | 5.5 | plasma membrane |
| | 4 | 6 | 5.5 | apoplast |
| | 6 | 4 | 3.6 | peroxisome |
| | 6 | 4 | 3.6 | cytoplasm |
| | 8 | 2 | 1.8 | mitochondrion |
| | 8 | 2 | 1.8 | mitochondrial inner membrane |
| | 9 | 1 | 0.9 | proteasome complex |
| | 9 | 1 | 0.9 | ribosome |
| | 9 | 1 | 0.9 | phosphopyruvate hydratase complex |
| | 9 | 1 | 0.9 | Golgi membrane |
| | 9 | 1 | 0.9 | monoatomic ion channel complex |
| | 9 | 1 | 0.9 | peroxisomal membrane |
| | 9 | 1 | 0.9 | endoplasmic reticulum membrane |
| | 9 | 1 | 0.9 | intracellular membrane-bounded organelle |
| | 9 | 1 | 0.9 | ribonucleoprotein complex |
| | 9 | 1 | 0.9 | endoplasmic reticulum |
| | 9 | 1 | 0.9 | vacuolar membrane |
| | 9 | 1 | 0.9 | vacuole |
| | 9 | 1 | 0.9 | organelle membrane |
| | 9 | 1 | 0.9 | chloroplast membrane |
| **Down-regulation** | | | | |
| Overall | GO | Hits | Ratio (%) | |
| | mapped | 174 | 28.57 | |
| | unmapped | 435 | 71.43 | |
| Top 10 MF Terms | Molecular Function (MF), total 71 hits | | | |
| | Enrichment | Count | Ratio (%) | Elements |
| | 1 | 14 | 7.61 | oxidoreductase activity |
| | 1 | 14 | 7.61 | metal ion binding |
| | 3 | 10 | 5.43 | transferase activity |
| | 4 | 9 | 4.89 | chlorophyll binding |
| | 5 | 8 | 4.35 | DNA-binding transcription factor activity |
| | 6 | 6 | 3.26 | hydrolase activity |
| | 6 | 6 | 3.26 | DNA binding |
| | 8 | 5 | 2.72 | heme binding |
| | 8 | 5 | 2.72 | ATP binding |
| | 8 | 5 | 2.72 | peroxidase activity |
| | 8 | 5 | 2.72 | hydrolase activity, hydrolyzing O-glycosyl compounds |

<table>
<tr><td rowspan="12">Top 10 BP Terms</td><td colspan="4">Biological Process (BP), total 75 hits</td></tr>
<tr><td>Enrichment</td><td>Count</td><td>Ratio (%)</td><td>Elements</td></tr>
<tr><td>1</td><td>22</td><td>11.7</td><td>photosynthesis</td></tr>
<tr><td>2</td><td>21</td><td>11.2</td><td>regulation of DNA-templated transcription</td></tr>
<tr><td>3</td><td>9</td><td>4.8</td><td>photosynthesis, light harvesting</td></tr>
<tr><td>4</td><td>7</td><td>3.7</td><td>auxin-activated signaling pathway</td></tr>
<tr><td>4</td><td>7</td><td>3.7</td><td>response to auxin</td></tr>
<tr><td>4</td><td>7</td><td>3.7</td><td>cell wall organization</td></tr>
<tr><td>7</td><td>6</td><td>3.2</td><td>cellular oxidant detoxification</td></tr>
<tr><td>8</td><td>5</td><td>2.7</td><td>hydrogen peroxide catabolic process</td></tr>
<tr><td>8</td><td>5</td><td>2.7</td><td>response to oxidative stress</td></tr>
<tr><td>8</td><td>5</td><td>2.7</td><td>carbohydrate metabolic process</td></tr>
<tr><td rowspan="12">Top 10 BP Terms</td><td colspan="4">Cellular Component (CC), total 31 hits</td></tr>
<tr><td>Enrichment</td><td>Count</td><td>Ratio (%)</td><td>Elements</td></tr>
<tr><td>1</td><td>44</td><td>21.2</td><td>membrane</td></tr>
<tr><td>2</td><td>20</td><td>9.6</td><td>chloroplast</td></tr>
<tr><td>3</td><td>18</td><td>8.7</td><td>plastid</td></tr>
<tr><td>4</td><td>17</td><td>8.2</td><td>thylakoid</td></tr>
<tr><td>5</td><td>17</td><td>8.2</td><td>nucleus</td></tr>
<tr><td>6</td><td>16</td><td>7.7</td><td>photosystem I</td></tr>
<tr><td>7</td><td>14</td><td>6.7</td><td>chloroplast thylakoid membrane</td></tr>
<tr><td>8</td><td>14</td><td>6.7</td><td>photosystem II</td></tr>
<tr><td>9</td><td>13</td><td>6.3</td><td>extracellular region</td></tr>
<tr><td>10</td><td>5</td><td>2.4</td><td>apoplast</td></tr>
</table>

### Table S7 List of gene symbols used in KEGG analysis

| Metabolic Pathway (PathID) | geneID | KOID | Description | Symbol |
|---|---|---|---|---|
| Glycolysis / Gluconeogenesis (mus00010) | 103977033 | K00002 | aldo-keto reductase family 4 member C9-like | ADH |
| | 103988073 | K00128 | aldehyde dehydrogenase family 3 member H1-like | ALDH3 |
| | 103992275 | K00131 | NADP-dependent glyceraldehyde-3-phosphate dehydrogenase-like | GAPDH |
| | 103995602 | K00134 | glyceraldehyde-3-phosphate dehydrogenase GAPCP1, chloroplastic | GAPDH |
| | 103969606 | K00382 | dihydrolipoyl dehydrogenase, mitochondrial-like | DLD |
| | 103971433 | K00873 | pyruvate kinase, cytosolic isozyme-like | PK |
| | 103986225 | K00873 | pyruvate kinase, cytosolic isozyme | PK |
| | 103997454 | K00873 | pyruvate kinase 1, cytosolic | PK |
| | 103987726 | K00873 | pyruvate kinase 1, cytosolic-like | PK |
| | 103982121 | K00927 | phosphoglycerate kinase, chloroplastic | PGK |
| | 103969489 | K01006 | pyruvate, phosphate dikinase 2 | PPDK |
| | 103993470 | K01623 | fructose-bisphosphate aldolase 1, cytoplasmic-like | ALDO |
| | 103974221 | K01623 | fructose-bisphosphate aldolase 1, chloroplastic-like | ALDO |
| | 103987382 | K01623 | fructose-bisphosphate aldolase 1, chloroplastic-like | ALDO |
| | 103988094 | K01623 | fructose-bisphosphate aldolase, chloroplastic | ALDO |
| | 103977957 | K01623 | fructose-bisphosphate aldolase 5, cytosolic | ALDO |
| | 103997940 | K01689 | enolase 1, chloroplastic-like | ENO |
| | 103978811 | K01689 | enolase-like | ENO |
| | 103969164 | K01810 | glucose-6-phosphate isomerase 1, chloroplastic-like | GPI |
| | 103976941 | K01834 | uncharacterized protein LOC103976941 | PGAM |
| | 104000631 | K01895 | acetyl-coenzyme A synthetase, chloroplastic/glyoxysomal | ACS |
| | 103977657 | K01913 | acetate/butyrate--CoA ligase AAE7, peroxisomal-like | AAE7 |
| | 103995197 | K14085 | aldehyde dehydrogenase family 7 member B4 | ALDH7 |
| Pyruvate (mus00620) | 103977033 | K00002 | aldo-keto reductase family 4 member C9-like | ADH |
| | 103986793 | K00026 | malate dehydrogenase, chloroplastic-like | MDH NAD+ |
| | 103991412 | K00026 | malate dehydrogenase, chloroplastic-like | MDH NAD+ |
| | 103973387 | K00051 | malate dehydrogenase [NADP], chloroplastic | MDH NADP+ |
| | 103988073 | K00128 | aldehyde dehydrogenase family 3 member H1-like | ALDH3 |
| | 103969606 | K00382 | dihydrolipoyl dehydrogenase, mitochondrial-like | DLD |
| | 103971433 | K00873 | pyruvate kinase, cytosolic isozyme-like | PK |
| | 103986225 | K00873 | pyruvate kinase, cytosolic isozyme | PK |
| | 103997454 | K00873 | pyruvate kinase 1, cytosolic | PK |

| | | | | |
|---|---|---|---|---|
| | 103987726 | K00873 | pyruvate kinase 1, cytosolic-like | PK |
| | 103969489 | K01006 | pyruvate, phosphate dikinase 2 | PPDK |
| | 103989174 | K01595 | phosphoenolpyruvate carboxylase 2-like | PEPC |
| | 103997092 | K01595 | phosphoenolpyruvate carboxylase 2-like | PEPC |
| | 103992398 | K01595 | phosphoenolpyruvate carboxylase 4 | PEPC |
| | 103995996 | K01638 | malate synthase, glyoxysomal | MS |
| | 104000631 | K01895 | acetyl-coenzyme A synthetase, chloroplastic/glyoxysomal | ACS |
| | 103977657 | K01913 | acetate/butyrate--CoA ligase AAE7, peroxisomal-like | AAE7 |
| | 103995197 | K14085 | aldehyde dehydrogenase family 7 member B4 | ALDH7 |
| Citrate cycle (TCA cycle) (mus00020) | 103986793 | K00026 | malate dehydrogenase, chloroplastic-like | MDH |
| | 103991412 | K00026 | malate dehydrogenase, chloroplastic-like | MDH |
| | 103989292 | K00030 | isocitrate dehydrogenase [NAD] regulatory subunit 1, mitochondrial-like | IDH NAD+ |
| | 103972343 | K00031 | cytosolic isocitrate dehydrogenase [NADP]-like | IDH NADP+ |
| | 103986147 | K00164 | 2-oxoglutarate dehydrogenase, mitochondrial-like | OGDC |
| | 103969606 | K00382 | dihydrolipoyl dehydrogenase, mitochondrial-like | DLD |
| | 103983279 | K01647 | LOW QUALITY PROTEIN: citrate synthase, mitochondrial-like | CS |
| | 103970434 | K01648 | ATP-citrate synthase beta chain protein 1 | ACLY |
| | 103972390 | K01648 | ATP-citrate synthase alpha chain protein 2 | ACLY |
| | 103970120 | K01681 | aconitate hydratase, cytoplasmic-like | ACO |
| Oxidative phosphorylation (mus00190) | 103985482 | K01535 | plasma membrane ATPase | PM ATPase |
| | 103972227 | K01535 | plasma membrane ATPase-like | PM ATPase |
| | 103969684 | K01535 | plasma membrane ATPase | PM ATPase |
| | 103981973 | K01535 | plasma membrane ATPase-like | PM ATPase |
| | 103993768 | K02147 | V-type proton ATPase subunit B 2 | V-ATPase |
| | 103982075 | K03885 | external alternative NAD(P)H-ubiquinone oxidoreductase B2, mitochondrial | NDH |
| | 103988136 | K08738 | cytochrome c | CYC |
| ABC transporters (mus02010) | 103979001 | K08711 | ABC transporter G family member 45 | ABCG2/PDR5 |
| | 103990131 | K08711 | ABC transporter G family member 36 | ABCG2/PDR5 |
| | 103985721 | K08711 | ABC transporter G family member 36-like isoform X1 | ABCG2/PDR5 |
| | 103996808 | K08711 | ABC transporter G family member 36-like | ABCG2/PDR5 |
| | 103998064 | K08711 | ABC transporter G family member 42-like | ABCG2/PDR5 |
| | 103993578 | K08711 | pleiotropic drug resistance protein 2-like | ABCG2/PDR5 |
| | 103971262 | K05681 | ABC transporter G family member 25-like | ABCG2 |
| | 103974315 | K05681 | ABC transporter G family member 11-like | ABCG2 |

| | | | | |
|---|---|---|---|---|
| | 103985729 | K05681 | ABC transporter G family member 11-like | ABCG2 |
| | 103984607 | K05681 | ABC transporter G family member 11-like | ABCG2 |
| | 103983286 | K05658 | putative multidrug resistance protein isoform X1 | ABCB1 |
| | 103972762 | K05658 | ABC transporter B family member 4 isoform X1 | ABCB1 |
| | 103981822 | K05658 | ABC transporter B family member 2-like | ABCB1 |
| | 103981250 | K05658 | ABC transporter B family member 1 | ABCB1 |
| Glutathione metabolism (mus00480) | 103972343 | K00031 | cytosolic isocitrate dehydrogenase [NADP]-like | IDH |
| | 103983842 | K00033 | 6-phosphogluconate dehydrogenase, decarboxylating 1 | PGD |
| | 103987991 | K00036 | glucose-6-phosphate 1-dehydrogenase, chloroplastic-like | G6PD |
| | 103989311 | K00036 | glucose-6-phosphate 1-dehydrogenase, cytoplasmic isoform-like | G6PD |
| | 103968393 | K00434 | probable L-ascorbate peroxidase 6, chloroplastic isoform X1 | APX |
| | 103999071 | K00799 | glutathione S-transferase U20-like | GST |
| | 103997659 | K00799 | glutathione S-transferase U17-like | GST |
| | 103976154 | K00799 | glutathione S-transferase U18-like | GST |
| | 103984645 | K00799 | glutathione S-transferase U18-like | GST |
| | 103984648 | K00799 | glutathione S-transferase U18-like | GST |
| | 103989732 | K00799 | protein IN2-1 homolog B | GST |
| | 103984516 | K00799 | glutathione S-transferase U10-like | GST |
| | 103985287 | K00799 | glutathione S-transferase PARB-like | GST |
| | 103999735 | K00799 | glutathione transferase GST 23-like | GST |
| | 103996908 | K01469 | 5-oxoprolinase | OXP |
| | 103987881 | K18592 | gamma-glutamyltranspeptidase 3 | GGT |
| Peroxisome (mus04146) | 103972343 | K00031 | cytosolic isocitrate dehydrogenase [NADP]-like | IDH |
| | 103976712 | K00232 | acyl-coenzyme A oxidase 2, peroxisomal | ACOX |
| | 103976051 | K01897 | long chain acyl-CoA synthetase 9, chloroplastic | ACSL |
| | 103987074 | K01897 | long chain acyl-CoA synthetase 9, chloroplastic-like | ACSL |
| | 103993123 | K01897 | long chain acyl-CoA synthetase 2 | ACSL |
| | 103978257 | K03781 | catalase isozyme A-like isoform X1 | CAT |
| | 104000193 | K04565 | superoxide dismutase [Cu-Zn], chloroplastic | SOD |
| | 103973111 | K07513 | 3-ketoacyl-CoA thiolase 2, peroxisomal-like isoform X1 | ACAA |
| | 103988537 | K11517 | peroxisomal (S)-2-hydroxy-acid oxidase GLO1 | HAO |
| | 103999561 | K13356 | fatty acyl-CoA reductase 3-like | FAR |
| MAPK signalling pathway - plant (mus04016) | 103982436 | K20772 | 1-aminocyclopropane-1-carboxylate synthase | ACS |
| | 103990761 | K20725 | protein MKS1-like | MKS1 |

| | | | | |
|---|---|---|---|---|
| | 103990855 | K20718 | LRR receptor-like serine/threonine-protein kinase ERECTA isoform X4 | ERL/ERECTA |
| | 103983253 | K20718 | LRR receptor-like serine/threonine-protein kinase ERL1 | ERL/ERECTA |
| | 103987224 | K20716 | mitogen-activated protein kinase kinase kinase NPK1-like | MAPKKK |
| | 103972093 | K14516 | ethylene-responsive transcription factor 1B-like | ERF1 |
| | 103995384 | K14514 | ETHYLENE INSENSITIVE 3-like 1 protein | EIN3 |
| | 103970202 | K14509 | ethylene receptor 2-like | ETR |
| | 103986109 | K14497 | probable protein phosphatase 2C 30 | PP2Cs |
| | 103970203 | K14497 | probable protein phosphatase 2C 75 | PP2Cs |
| | 103989695 | K14496 | abscisic acid receptor PYL4-like | PYL |
| | 103975648 | K13449 | pathogenesis-related protein 1-like | PR1 |
| | 103987581 | K13447 | respiratory burst oxidase homolog protein B | RBOH |
| | 103994080 | K13424 | probable WRKY transcription factor 26 isoform X3 | WRKY |
| | 103989687 | K13424 | WRKY transcription factor WRKY24-like | WRKY |
| | 103994582 | K13422 | transcription factor MYC2-like | MYC2 |
| | 103978257 | K03781 | catalase isozyme A-like isoform X1 | CAT |
| | 103968609 | K02183 | probable calcium-binding protein CML45 | CALM/CML |
| Plant-pathogen interaction (mus04626) | 103968609 | K02183 | probable calcium-binding protein CML45 | CALM/CML |
| | 103999891 | K04079 | heat shock protein 81-1-like | HSPs |
| | 103979428 | K09487 | endoplasmin homolog | HSPs |
| | 103980807 | K13412 | calcium-dependent protein kinase 20-like | CPK |
| | 103973106 | K13412 | calcium-dependent protein kinase 15 | CPK |
| | 103994080 | K13424 | probable WRKY transcription factor 26 isoform X3 | WRKY |
| | 103989687 | K13424 | WRKY transcription factor WRKY24-like | WRKY |
| | 103971903 | K13434 | ethylene-responsive transcription factor ERF069-like | PT! |
| | 103987581 | K13447 | respiratory burst oxidase homolog protein B | RBOH |
| | 103979085 | K13448 | probable calcium-binding protein CML48 | CML |
| | 103975648 | K13449 | pathogenesis-related protein 1-like | PR1 |
| | 103999439 | K13457 | disease resistance protein RPM1-like | RPM |
| | 103990292 | K15397 | 3-ketoacyl-CoA synthase 11-like | KCS |
| | 103976761 | K15397 | 3-ketoacyl-CoA synthase 6-like | KCS |
| | 103987574 | K15397 | 3-ketoacyl-CoA synthase 10-like | KCS |
| | 103984568 | K15397 | 3-ketoacyl-CoA synthase 1-like | KCS |
| | 103977157 | K15397 | 3-ketoacyl-CoA synthase 10-like | KCS |
| | 103973066 | K15397 | 3-ketoacyl-CoA synthase 11-like | KCS |

| | | | | |
|---|---|---|---|---|
| | 103988964 | K15397 | 3-ketoacyl-CoA synthase 12-like | KCS |
| | 103977577 | K15397 | 3-ketoacyl-CoA synthase 6 | KCS |
| Plant hormone signal transduction (mus04075) | 103985482 | K01535 | plasma membrane ATPase | PM ATPase |
| | 103972227 | K01535 | plasma membrane ATPase-like | PM ATPase |
| | 103969684 | K01535 | plasma membrane ATPase | PM ATPase |
| | 103981973 | K01535 | plasma membrane ATPase-like | PM ATPase |
| | 103973700 | K13415 | brassinosteroid LRR receptor kinase-like | BRI1 |
| | 103994582 | K13422 | transcription factor MYC2-like | MYC2 |
| | 103975648 | K13449 | pathogenesis-related protein 1-like | PR1 |
| | 103977146 | K13464 | protein TIFY 10a-like | JAZ/TIFY |
| | 103996636 | K13464 | protein TIFY 9 | JAZ/TIFY |
| | 103983987 | K13464 | protein TIFY 10a-like | JAZ/TIFY |
| | 103977817 | K13464 | protein TIFY 10a-like | JAZ/TIFY |
| | 103997923 | K13464 | protein TIFY 10b-like | JAZ/TIFY |
| | 103970778 | K13464 | protein TIFY 10a | JAZ/TIFY |
| | 103984053 | K13464 | protein TIFY 6b-like isoform X2 | JAZ/TIFY |
| | 103994040 | K13946 | auxin transporter-like protein 4 | AUX1/LAX |
| | 103986124 | K13946 | auxin transporter-like protein 1 | AUX1/LAX |
| | 103990205 | K13946 | auxin transporter-like protein 2 | AUX1/LAX |
| | 104000242 | K14431 | transcription factor TGAL5-like isoform X1 | TGA |
| | 103998485 | K14431 | transcription factor TGA2.1-like isoform X1 | TGA |
| | 103997082 | K14484 | auxin-responsive protein IAA21 | IAAs |
| | 103969278 | K14484 | auxin-responsive protein IAA17-like | IAAs |
| | 103995121 | K14484 | auxin-responsive protein IAA30 isoform X1 | IAAs |
| | 103989163 | K14484 | auxin-responsive protein IAA21-like | IAAs |
| | 103997574 | K14484 | auxin-responsive protein IAA30 isoform X1 | IAAs |
| | 103991365 | K14484 | auxin-responsive protein IAA17-like | IAAs |
| | 103995122 | K14484 | auxin-induced protein 22D-like | IAAs |
| | 103997573 | K14484 | auxin-responsive protein IAA31-like | IAAs |
| | 103999151 | K14484 | auxin-responsive protein IAA9 isoform X1 | IAAs |
| | 103983123 | K14484 | auxin-induced protein 22D | IAAs |
| | 103983124 | K14484 | auxin-responsive protein IAA30 | IAAs |
| | 103976465 | K14486 | auxin response factor 11-like | ARF |
| | 103980181 | K14487 | probable indole-3-acetic acid-amido synthetase GH3.8 | GH3 |

| | | | | |
|---|---|---|---|---|
| | 104000870 | K14490 | pseudo histidine-containing phosphotransfer protein 2 | HP |
| | 103988140 | K14492 | two-component response regulator ORR1 | RP |
| | 103987951 | K14493 | gibberellin receptor GID1C-like | GID1 |
| | 103989695 | K14496 | abscisic acid receptor PYL4-like | PYL |
| | 103986109 | K14497 | probable protein phosphatase 2C 30 | PP2Cs |
| | 103970203 | K14497 | probable protein phosphatase 2C 75 | PP2Cs |
| | 103970202 | K14509 | ethylene receptor 2-like | ETR |
| | 103995384 | K14514 | ETHYLENE INSENSITIVE 3-like 1 protein | EIN3 |
| | 103972093 | K14516 | ethylene-responsive transcription factor 1B-like | ERF1 |
| | 103989881 | K16189 | transcription factor PIF1-like isoform X1 | PIF |
| NAD/NADP-dependent dehydrogenase | 103995258 | K00207 | dihydropyrimidine dehydrogenase (NADP(+)), chloroplastic | DPYD |
| | 103989292 | K00030 | isocitrate dehydrogenase [NAD] regulatory subunit 1, mitochondrial-like | IDHs |
| | 103972343 | K00031 | cytosolic isocitrate dehydrogenase [NADP]-like | IDHs |
| | 103992275 | K00131 | NADP-dependent glyceraldehyde-3-phosphate dehydrogenase-like | GAPDH |
| | 103973387 | K00051 | malate dehydrogenase [NADP], chloroplastic | MDH |
| | 103970505 | K00006 | probable glycerol-3-phosphate dehydrogenase [NAD(+)] 1, cytosolic isoform X1 | GPDH |
| NAD/NADP-dependent enzyme | 103977556 | K10534 | nitrate reductase [NADH]-like | NR |
| | 103982075 | K03885 | external alternative NAD(P)H-ubiquinone oxidoreductase B2, mitochondrial | FAD/FMN |
| | 103982242 | K02641 | ferredoxin--NADP reductase, embryo isozyme, chloroplastic-like | FNR |
| | 103977597 | K22374 | non-functional NADPH-dependent codeinone reductase 2-like | DMAS |
| | 103982502 | K02641 | ferredoxin--NADP reductase, root isozyme, chloroplastic | FNR |
| | 103994455 | K02641 | ferredoxin--NADP reductase, leaf isozyme, chloroplastic-like | FNR |
| Dehydrogenase | 103987519 | K00012 | UDP-glucose 6-dehydrogenase 5-like | UGDH |
| | 103975873 | K00261 | glutamate dehydrogenase 2 | GLDH |
| | 103985656 | K00318 | proline dehydrogenase 1, mitochondrial-like | PRODH |
| | 103990067 | K15227 | arogenate dehydrogenase 1, chloroplastic | TYRAAT |
| | 103986793 | K00026 | malate dehydrogenase, chloroplastic-like | MDH |
| | 103995602 | K00134 | glyceraldehyde-3-phosphate dehydrogenase GAPCP1, chloroplastic | GAPDH |
| | 103978737 | K12355 | aldehyde dehydrogenase family 2 member C4-like | REF |
| | 103986147 | K00164 | 2-oxoglutarate dehydrogenase, mitochondrial-like | OGDH |
| | 103987991 | K00036 | glucose-6-phosphate 1-dehydrogenase, chloroplastic-like | G6PDH |
| | 103995197 | K14085 | aldehyde dehydrogenase family 7 member B4 | ALDH7 |
| | 103983842 | K00033 | 6-phosphogluconate dehydrogenase, decarboxylating 1 | PGD |
| | 103989311 | K00036 | glucose-6-phosphate 1-dehydrogenase, cytoplasmic isoform-like | G6PDH |

| | | | | |
|---|---|---|---|---|
| | 103969606 | K00382 | dihydrolipoyl dehydrogenase, mitochondrial-like | DLD |
| | 103991412 | K00026 | malate dehydrogenase, chloroplastic-like | MDH |
| | 103972115 | K05298 | glyceraldehyde-3-phosphate dehydrogenase B, chloroplastic isoform X1 | GAPDH |
| | 104000636 | K05298 | glyceraldehyde-3-phosphate dehydrogenase A, chloroplastic | GAPDH |
| | 103978746 | K00281 | glycine dehydrogenase (decarboxylating), mitochondrial | GLDH |
| | 103987962 | K05298 | glyceraldehyde-3-phosphate dehydrogenase A, chloroplastic-like | GAPDH |
| | 103988073 | K00128 | aldehyde dehydrogenase family 3 member H1-like | ALDH |
| | 103986978 | K12355 | aldehyde dehydrogenase family 2 member C4 | REF |
| | 103991720 | K11153 | short-chain dehydrogenase TIC 32, chloroplastic-like | RDH |
| | 103981139 | K00012 | UDP-glucose 6-dehydrogenase 4-like | UGDH |
| | 103995443 | K11153 | short-chain dehydrogenase TIC 32, chloroplastic | RDH |

### Table S8 List of genes associated with the immunity of wild bananas against *Fusarium* wilt

| Symbols | KEGG PathID | geneID | KOID | Description |
|---|---|---|---|---|
| PR1 | mus04075 | 103975648 | K13449 | pathogenesis-related protein 1-like |
| CHIT | mus00520 | 103973814 | K01183 | acidic endochitinase-like |
| | | 103975026 | K01183 | chitinase 6-like |
| | | 103985238 | K01183 | chitinase 6-like |
| | | 103988682 | K01183 | chitinase 6 |
| | | 103989476 | K01183 | acidic mammalian chitinase-like |
| POD | mus00480 | 103968393 | K00434 | probable L-ascorbate peroxidase 6, chloroplastic isoform X1 |
| | | 103972200 | K00430 | peroxidase 60-like isoform X1 |
| | | 103972232 | K00430 | peroxidase 4 |
| | | 103972416 | K00430 | peroxidase 3 |
| | | 103974374 | K00430 | peroxidase 47-like |
| | | 103975462 | K00430 | peroxidase P7-like isoform X2 |
| | | 103976746 | K00430 | peroxidase 59-like |
| | | 103976913 | K00430 | peroxidase 18-like |
| | | 103977010 | K00430 | peroxidase P7 |
| | | 103977073 | K00430 | cationic peroxidase SPC4-like |
| | | 103978205 | K00430 | peroxidase 21 |
| | | 103980512 | K00430 | peroxidase 18 isoform X1 |
| | | 103980763 | K00430 | peroxidase 72-like |
| | | 103985683 | K00430 | peroxidase 5-like |
| | | 103986030 | K00430 | peroxidase 51 |
| | | 103987043 | K00430 | peroxidase P7 |
| | | 103988131 | K00430 | peroxidase 47 |
| | | 103988707 | K00430 | peroxidase 2-like |
| | | 103988743 | K00430 | peroxidase 27-like |
| | | 103990806 | K00430 | peroxidase 55-like |
| | | 103990991 | K00430 | peroxidase N-like |
| | | 103991820 | K00430 | peroxidase A2-like |
| | | 103993172 | K00430 | peroxidase 5-like |
| | | 103993173 | K00430 | peroxidase 5 |
| | | 103993311 | K00430 | peroxidase 5-like |
| | | 103996217 | K00430 | peroxidase 25 |

| | | | | |
|---|---|---|---|---|
| PPO | mus00950 | 103968782 | K00422 | polyphenol oxidase I, chloroplastic-like |
| | | 103974846 | K00422 | polyphenol oxidase, chloroplastic-like |
| | | 103990742 | K00422 | polyphenol oxidase, chloroplastic-like |
| | | 103995522 | K00422 | polyphenol oxidase, chloroplastic-like |
| CYC | mus00905 | 103969391 | K12638 | cytochrome P450 90D2-like isoform X1 |
| | | 103985373 | K20623 | cytochrome P450 71A1-like |
| | | 103991557 | K15639 | cytochrome P450 734A6-like |
| | | 103996195 | K09587 | cytochrome P450 90B1 |
| | mus00073 | 103985950 | K15398 | cytochrome P450 86A8-like |
| | | 103993609 | K21995 | cytochrome P450 77A3-like |
| | | 103995930 | K21995 | cytochrome P450 77A2-like |
| | mus00380 | 103978547 | K24541 | cytochrome P450 71A1-like |
| | mus00100 | 103995072 | K09832 | cytochrome P450 710A1-like |
| WRKY | mus04626 | 103989687 | K13424 | WRKY transcription factor WRKY24-like |
| | | 103994080 | K13424 | probable WRKY transcription factor 26 isoform X3 |
| DCS | mus00941 | 103977235 | K00660 | phenylpropanoylacetyl-CoA synthase-like |
| KCS | mus04626 | 103973066 | K15397 | 3-ketoacyl-CoA synthase 11-like |
| | | 103976761 | K15397 | 3-ketoacyl-CoA synthase 6-like |
| | | 103977157 | K15397 | 3-ketoacyl-CoA synthase 10-like |
| | | 103977577 | K15397 | 3-ketoacyl-CoA synthase 6 |
| | | 103984568 | K15397 | 3-ketoacyl-CoA synthase 1-like |
| | | 103987574 | K15397 | 3-ketoacyl-CoA synthase 10-like |
| | | 103988964 | K15397 | 3-ketoacyl-CoA synthase 12-like |
| | | 103990292 | K15397 | 3-ketoacyl-CoA synthase 11-like |
| SUR2 | mus00600 | 103977198 | K04713 | sphinganine C4-monooxygenase 1-like |
