## Supporting Figures for "Endophytic pyrroloquinoline quinone enhances banana growth and immunity against *Fusarium* wilt for plant-microbe mutualisms"

**Running Title:** PQQ coordinates plant growth and immunity

Shih-Hsun Walter Hung[1,2,3,4], Man-Yun Yu[1], Chia-Ho Liu[1], Tsai-Ching Huang[1], Jian-Hau Peng[5,6], Nai-Yun Jang[1], Chih-Horng Kuo[2,5,6,7], Ying-Ning Ho[8,9,10], En-Pei Isabel Chiang[3,4,5,6,11], Hau-Hsuan Hwang[1,3,4,5,6], and Chieh-Chen Huang[1,3,4,5,6]*

[1] Department of Life Sciences, National Chung Hsing University, Taichung 402202, Taiwan; [2] Institute of Plant and Microbial Biology, Academia Sinica, Taipei 115201, Taiwan; [3] Advanced Plant and Food Crop Biotechnology Center, National Chung Hsing University, Taichung 402202, Taiwan; [4] Innovation and Development Centre of Sustainable Agriculture, National Chung Hsing University, Taichung 402202, Taiwan; [5] PhD Program in Microbial Genomics, National Chung Hsing University, Taichung 402202, Taiwan; [6] PhD Program in Microbial Genomics, Academia Sinica, Taipei 115201, Taiwan; [7] Biotechnology Center, National Chung Hsing University, Taichung 402202, Taiwan; [8] Institute of Marine Biology, College of Life Science, National Taiwan Ocean University, Keelung 202301, Taiwan; [9] Center of Excellence for the Oceans, National Taiwan Ocean University, Keelung 202301, Taiwan; [10] Taiwan Ocean Genome Center, National Taiwan Ocean University, Keelung 202301, Taiwan; [11] Department of Food Science and Biotechnology, National Chung Hsing University, Taichung 402202, Taiwan

**ORCID** SHW Hung, 0000-0002-8749-5653; CH Liu, 0000-0001-9166-3966; TC Huang, 0000-0002-3410-2051; JH Peng, 0000-0001-6052-8606; CH Kuo, 0000-0002-2857-0529; YN Ho, 0000-0003-0943-1416; EPI Chiang, 0000-0002-0158-0962; HH Hwang, 0000-0001-9132-0242; CC Huang, 0000-0002-3739-6315.

**Keywords:** endophyte, symbiosis, pyrroloquinoline quinone (PQQ), Fusarium wilt of banana, molecular plant-microbe interactions (MPMI)

Supplementary Figures

**Fig. S1 Endophytic PQQ production of *B. seminalis* 869T2.**

**Fig. S2 Early-staged exogenous supplementation of PQQ contributes to the later growth of banana plants transplanted to soil conditions.**

**Fig. S3 Higher dosage of PQQ exogenous supplementation effects on banana plants.**

**Fig. S4 KEGG predicted PQQ effects on ABC transporters regulations.**

**Fig. S5 KEGG predicted PQQ effects on glutathione metabolism regulations.**

**Fig. S6 KEGG predicted PQQ effects on peroxisome regulations.**

**Fig. S7 KEGG predicted PQQ effects on citrate cycle (TCA cycle) regulations.**

**Fig. S8 KEGG predicted PQQ effects on oxidative phosphorylation regulations.**

**Fig. S9 KEGG predicted PQQ effects on glycolysis/gluconeogenesis pathways regulations.**

**Fig. S10 KEGG predicted PQQ effects on pyruvate metabolism regulations.**

**Fig. S11 KEGG predicted PQQ effects on phenylpropanoid metabolism regulations.**

**Fig. S12 KEGG predicted PQQ effects on tryptophan metabolism regulations.**

**Fig. S13 KEGG predicted PQQ effects on tyrosine metabolism regulations.**

**Fig. S14 KEGG predicted PQQ effects on carotenoid biosynthesis regulations.**

**Fig. S15 KEGG predicted PQQ effects on the biosynthesis of various plant secondary metabolites.**

**Fig. S16 KEGG predicted PQQ effects on flavonoid biosynthesis regulations.**

**Fig. S17 KEGG predicted PQQ effects on plant hormone signal transduction regulations.**

**Fig. S18 KEGG predicted PQQ effects on MAPK signalling pathway – plant regulations.**

**Fig. S19 KEGG predicted PQQ effects on plant-pathogen interaction regulations.**

**Fig. S20 Organisation of the PQQ operon among *Burkholderia* species.**

**Fig. S21 Multiple protein sequence alignment of PQQ biosynthesis genes.**

**Fig. S22 Evolutionary divergence of PQQ operon among *Burkholderia* species.**

### Supplementary Figures and Legends

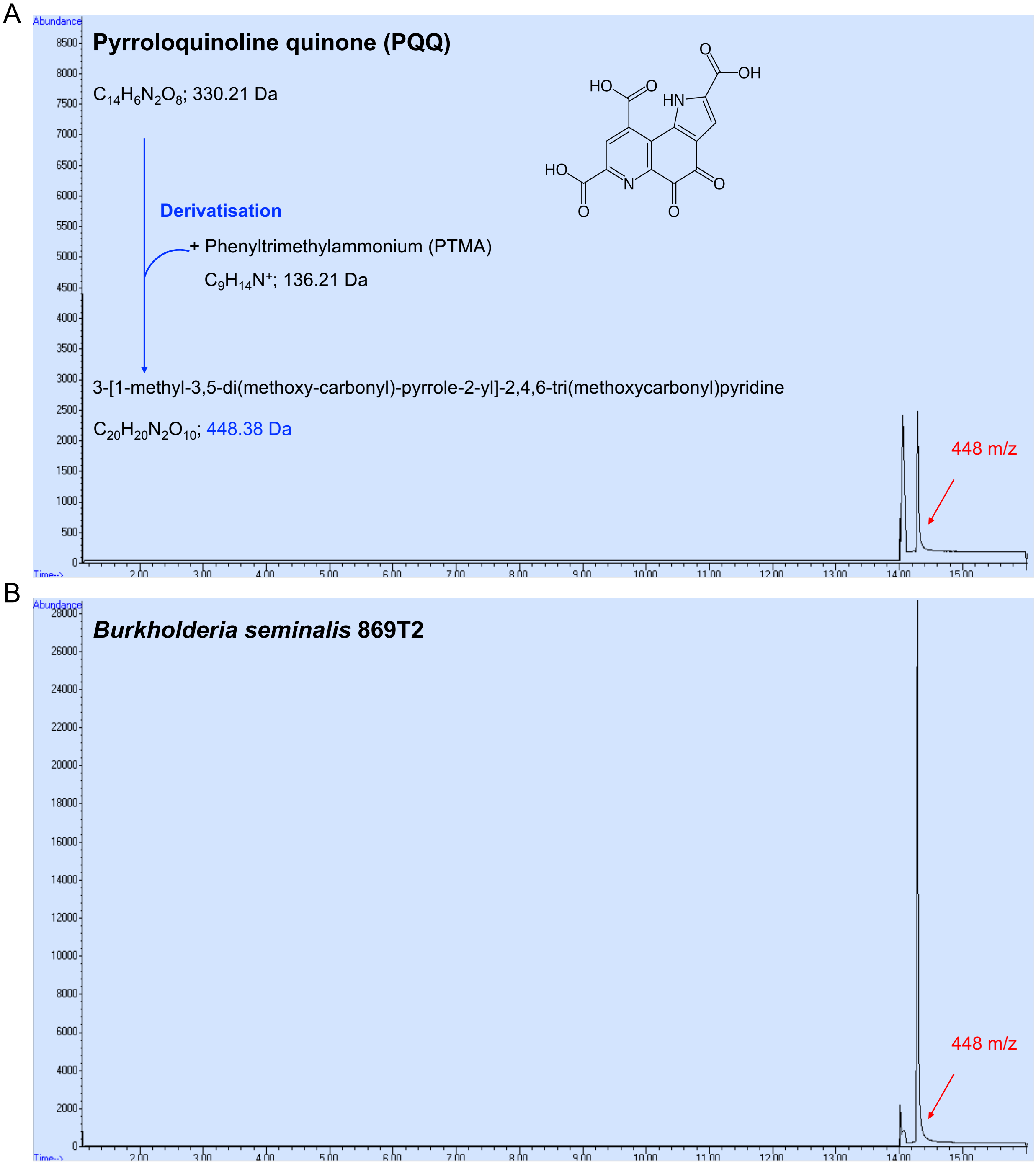


**Fig. S1 Endophytic PQQ production of *B. seminalis* 869T2.** The gas chromatography-mass spectrometry (GC-MS) result of **A** synthetic PQQ standard (>98%) and **B** extracted *B. seminalis* 869T2 cultures after PTMA derivatisation.

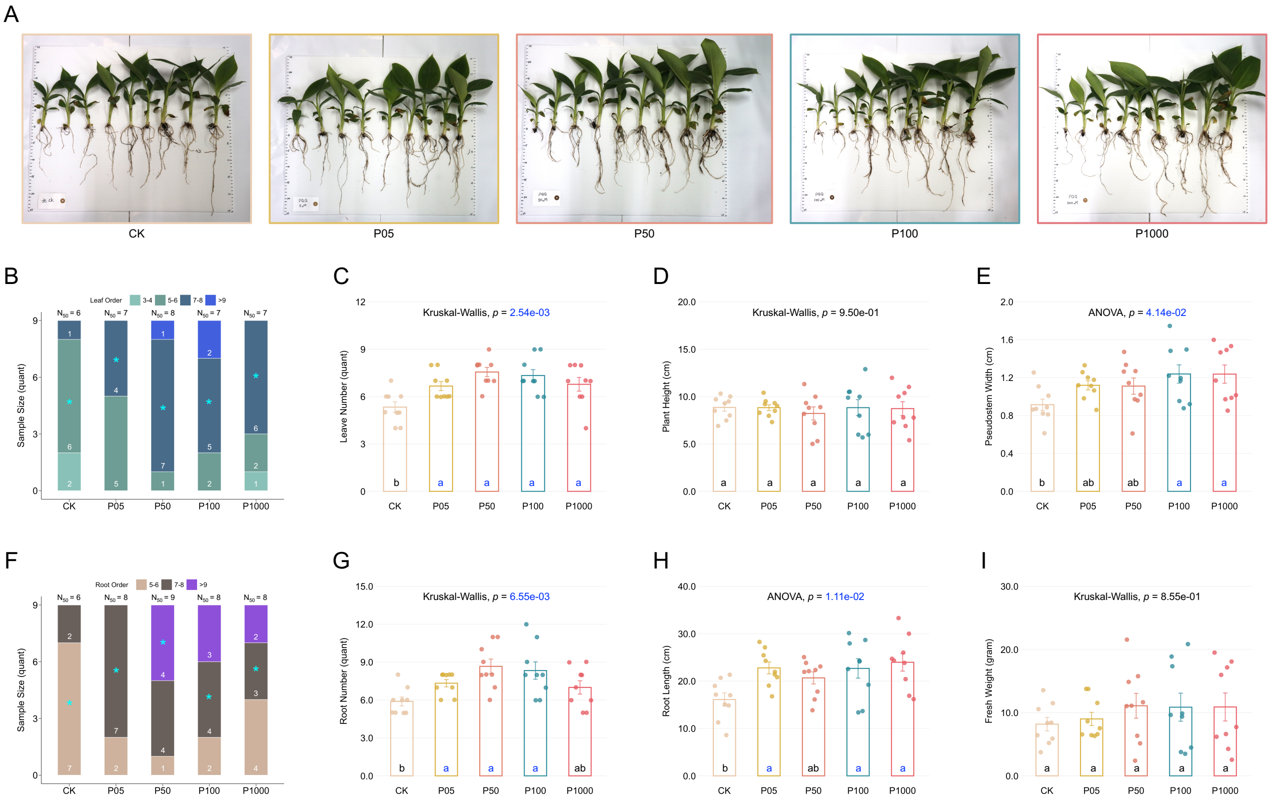


**Fig. S2 Early-staged exogenous supplementation of PQQ contributes to the later growth of banana plants transplanted to soil conditions. A** Phenotypic photograph of banana plants in response to different concentrations of supplementary PQQ at 137 days after treatment (DPT). CK, P05, P50, P100 and P1000 indicate the 0 nM, 5 nM, 50 nM, 100 nM and 1000 nM PQQ treatment, respectively. The distribution of **B** leaf number and **F** root number of banana plants at 137 DPT. Numbers in stacked bar charts indicate the sample size. Asterisks highlight the distribution groups of defined $N_{50}$ for plant growth-promotion evaluation, as mentioned in Materials and Methods. **C** Leaf number, **D** plant height, **E** pseudostem width, **G** root number, **H** root length and **I** fresh weight of banana plants at 137 DPT, excluding non-differentiated samples in **B** or **F**. Each dot represents a sample. Data are mean ± SEM; different letters indicate significant differences based on multiple comparisons, Tukey method after ANOVA or Dunn method after the Kruskal-Wallis test.

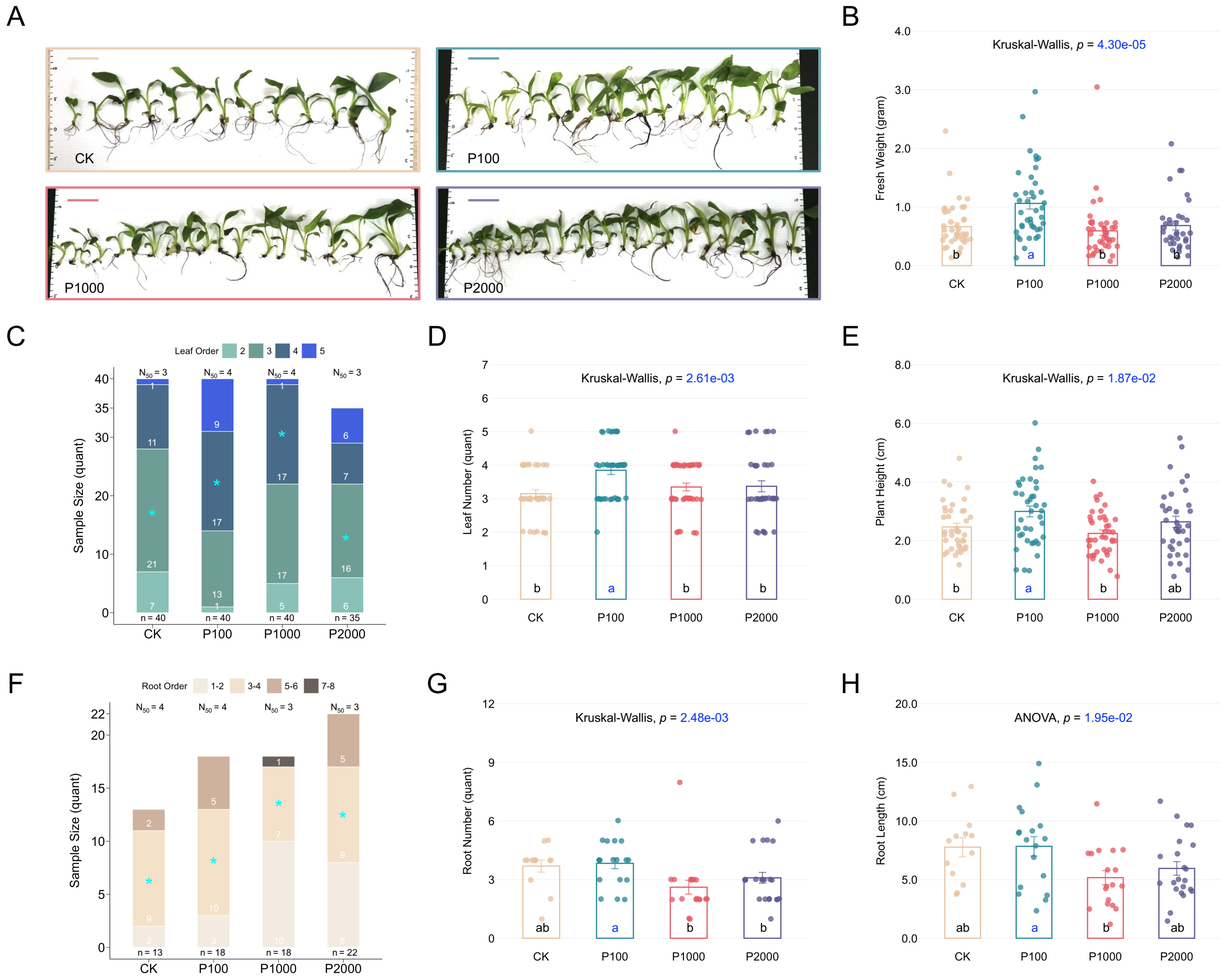


**Fig. S3 Higher dosage of PQQ exogenous supplementation effects on banana plants. A** Phenotypic photograph of banana plants in response to different concentrations of supplementary PQQ at 81 days after treatment (DPT). CK, P100, P1000 and P2000 indicate the 0 nM, 100 nM, 1000 nM and 2000 nM PQQ treatment, respectively. The distribution of **C** leaf number and **F** root number of banana plants at 81 DPT. Numbers in stacked bar charts indicate the sample size. Asterisks highlight the distribution groups of defined $N_{50}$ for plant growth-promotion evaluation, as mentioned in Materials and Methods. **B** Fresh weight, **D** leaf number, **E** plant height, **G** root number and **H** root length of banana plants at 81 DPT, excluding non-differentiated samples in **C** or **F**. Each dot represents a sample. Data are mean ± SEM; different letters indicate significant differences based on multiple comparisons, Tukey method after ANOVA or Dunn method after the Kruskal-Wallis test.

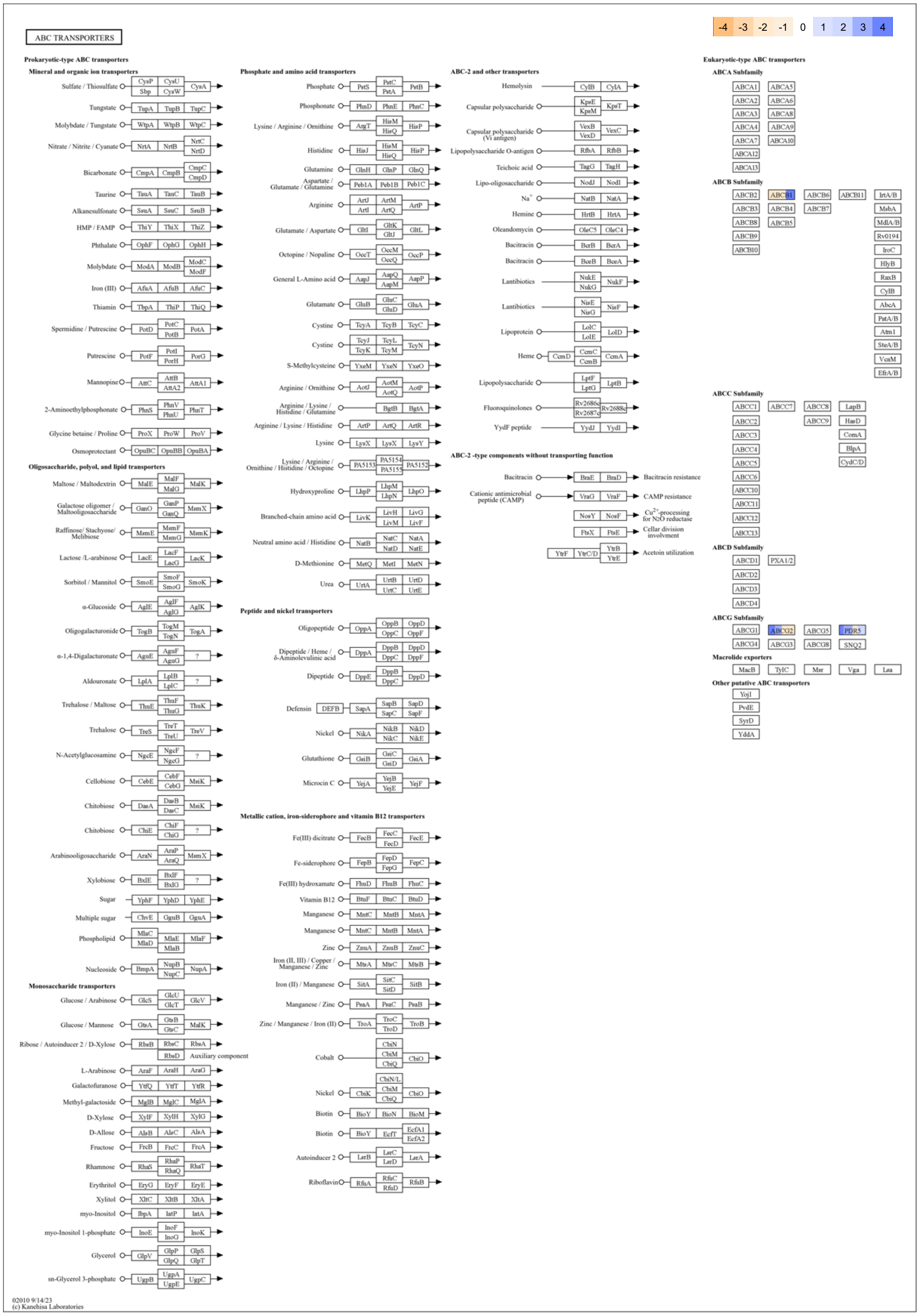


**Fig. S4 KEGG predicted PQQ effects on ABC transporters regulations.**

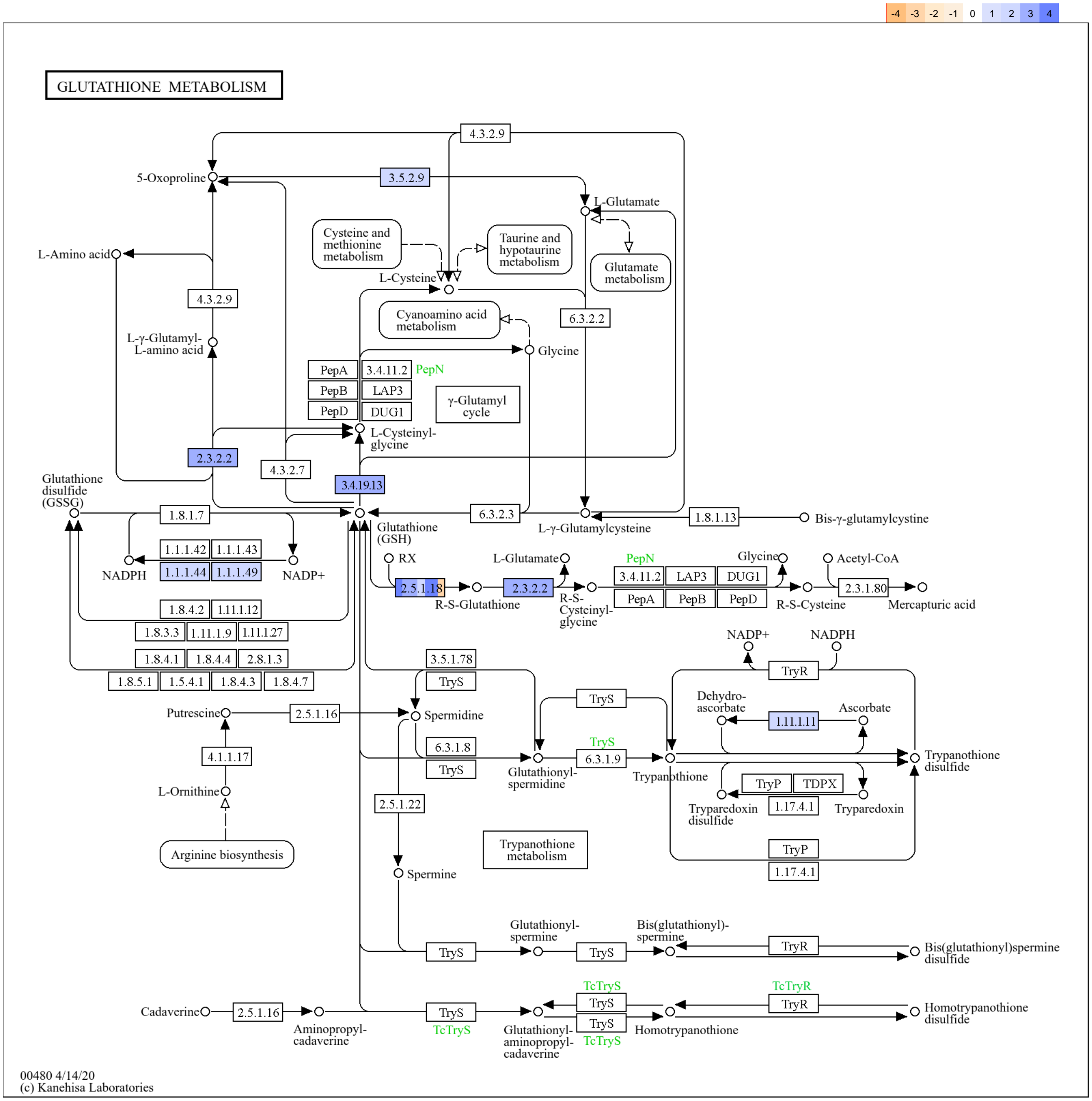


**Fig. S5 KEGG predicted PQQ effects on glutathione metabolism regulations.**

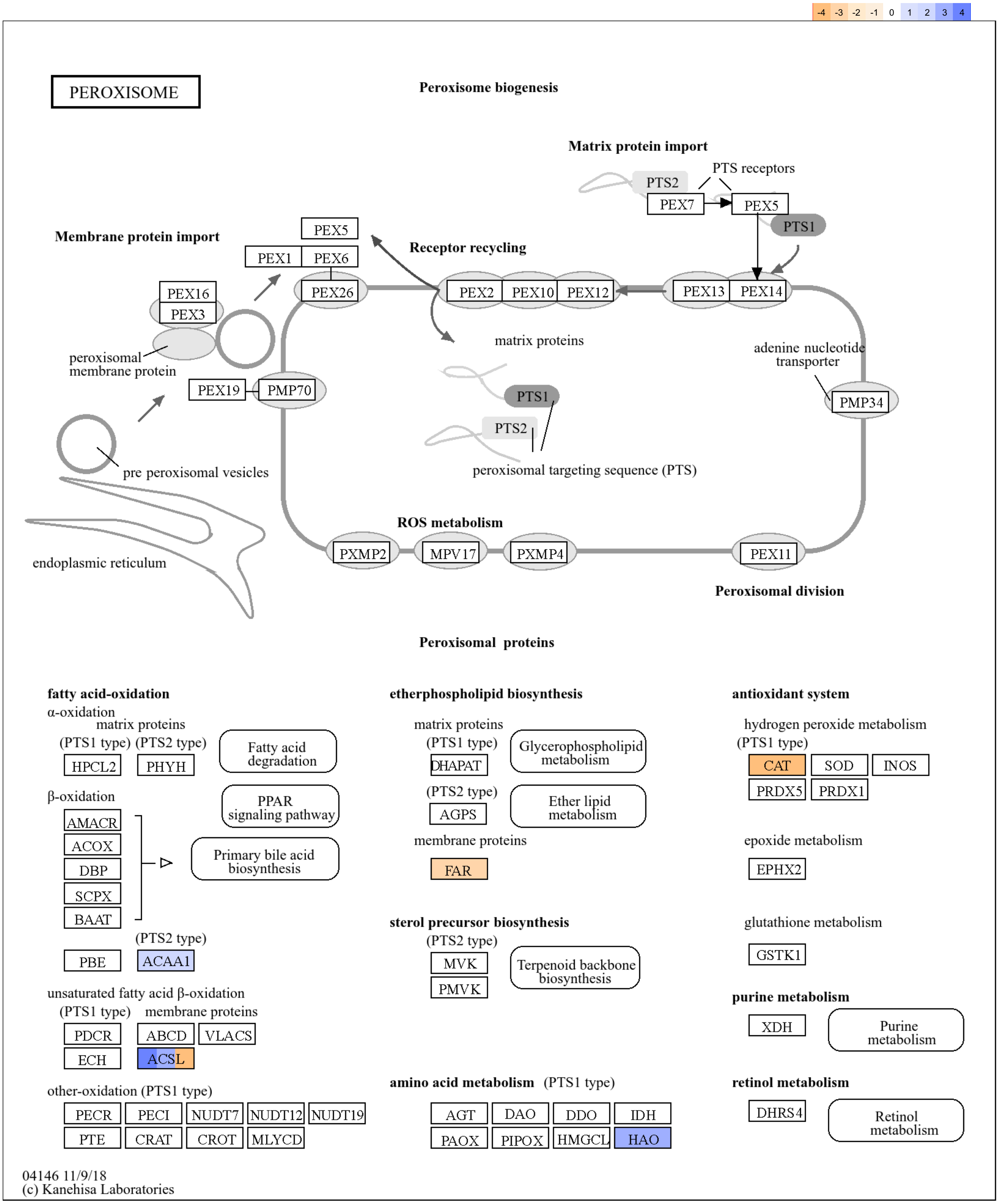


**Fig. S6 KEGG predicted PQQ effects on peroxisome regulations.**

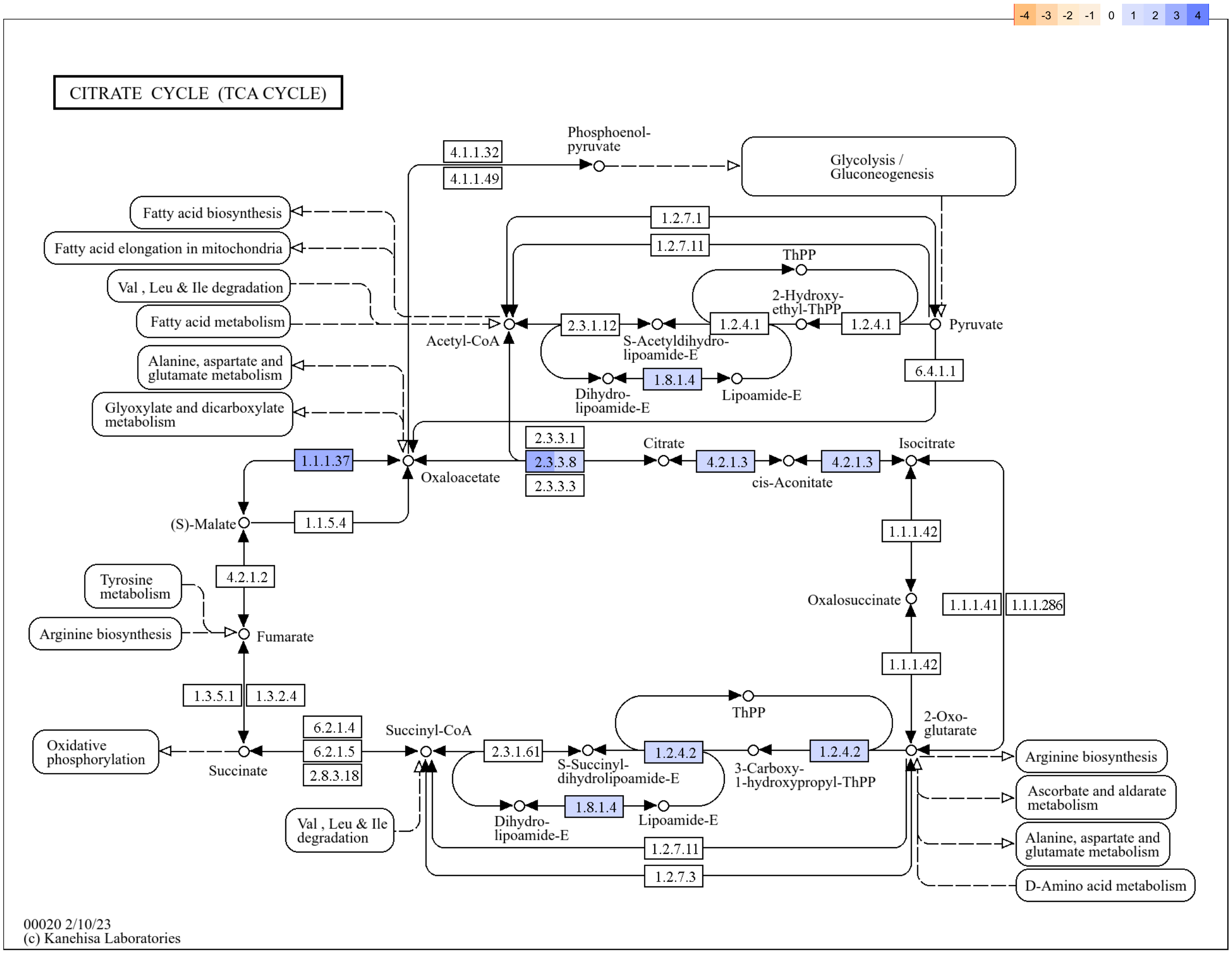


**Fig. S7 KEGG predicted PQQ effects on citrate cycle (TCA cycle) regulations.**

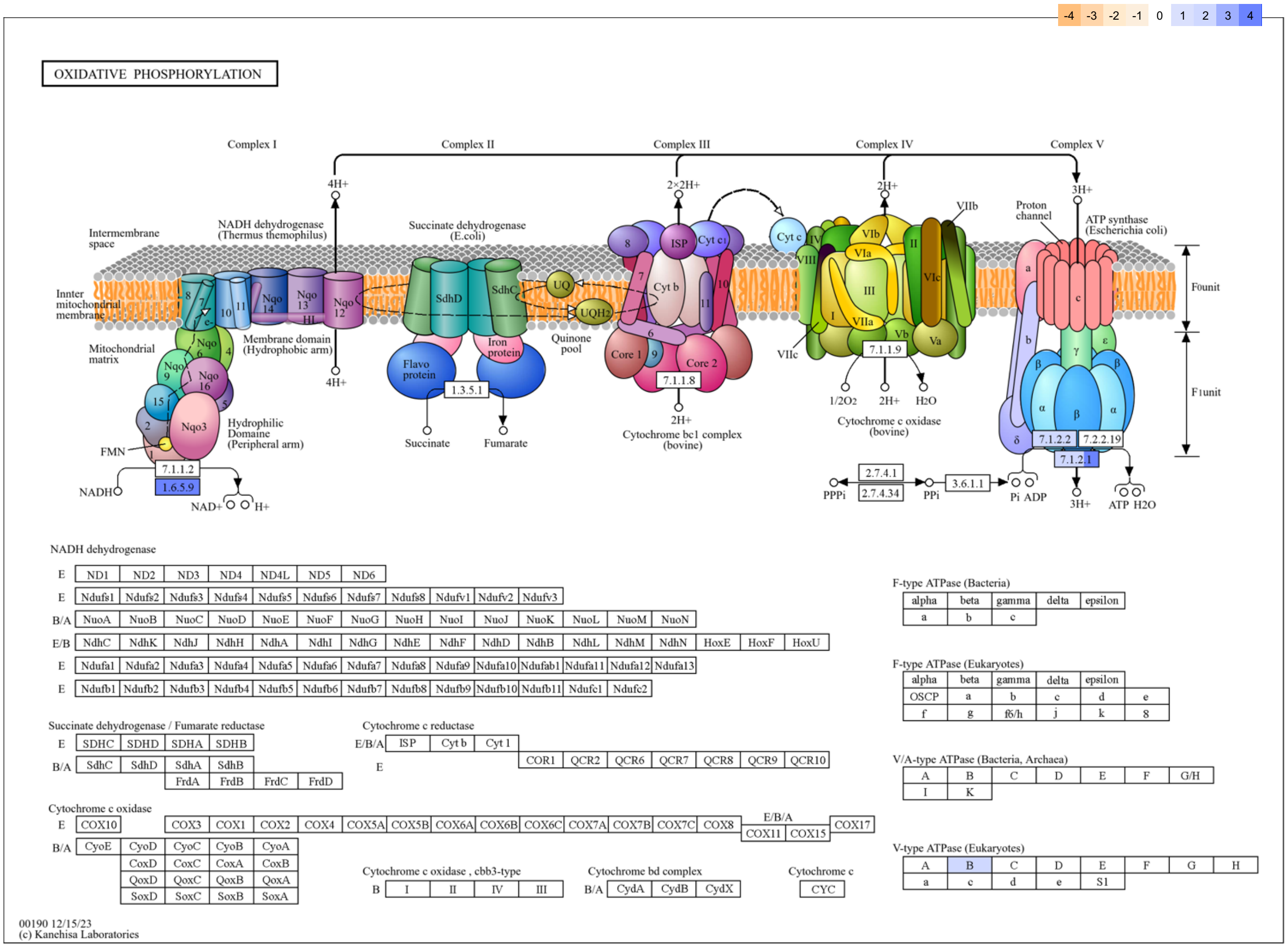


**Fig. S8 KEGG predicted PQQ effects on oxidative phosphorylation regulations.**

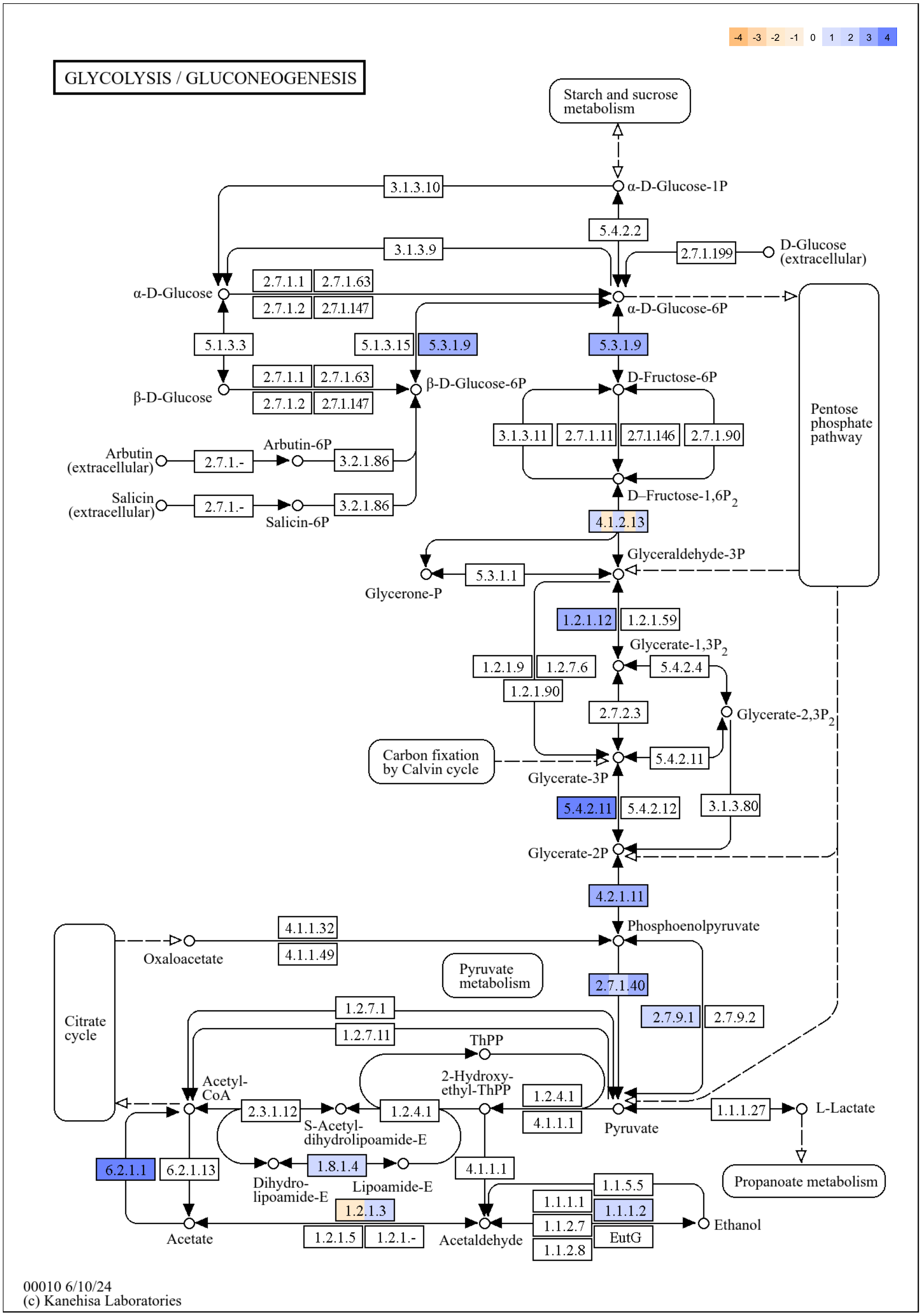


**Fig. S9 KEGG predicted PQQ effects on glycolysis/gluconeogenesis pathways regulations.**

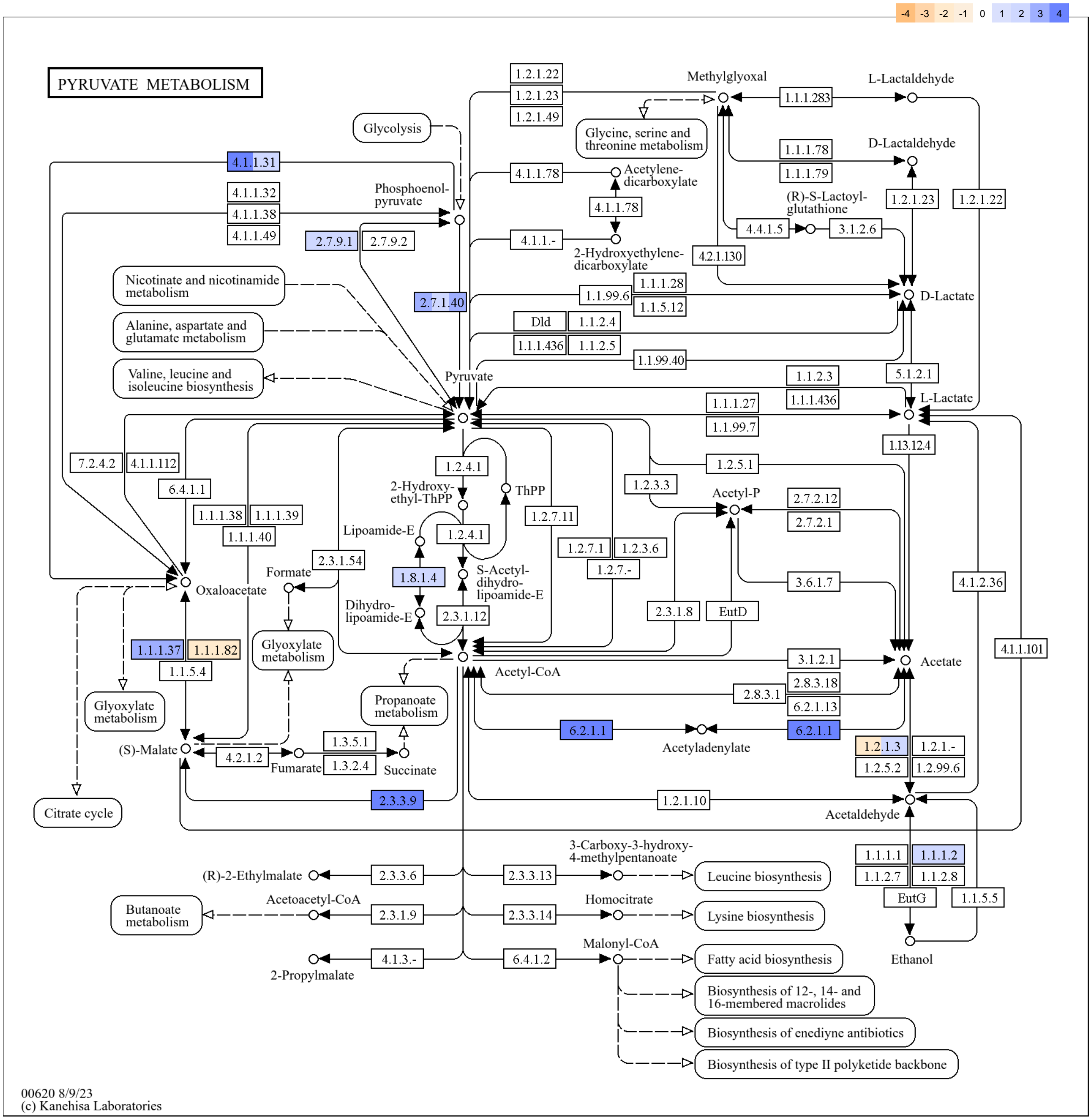


**Fig. S10 KEGG predicted PQQ effects on pyruvate metabolism regulations.**

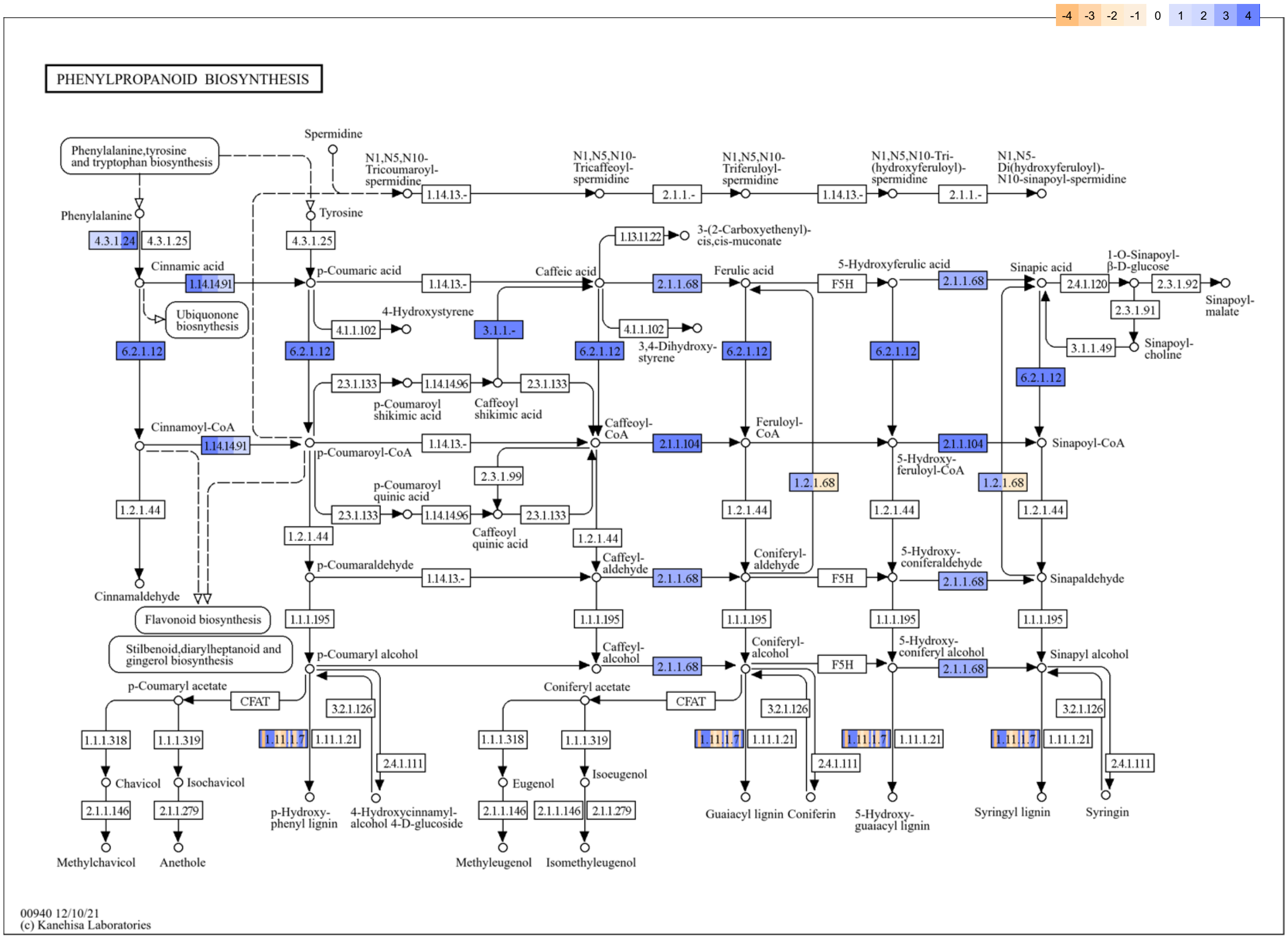


**Fig. S11 KEGG predicted PQQ effects on phenylpropanoid metabolism regulations.**

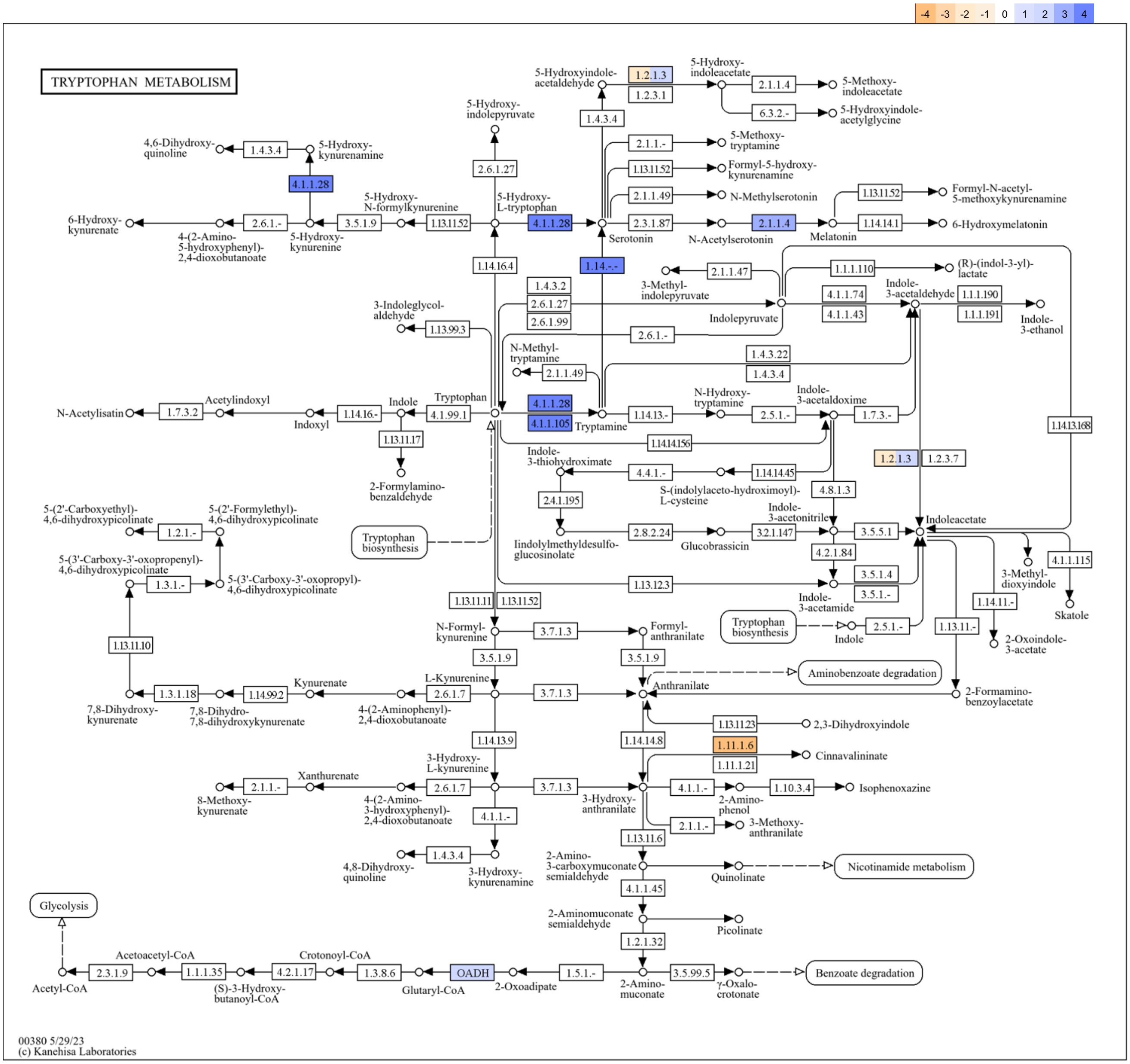


**Fig. S12 KEGG predicted PQQ effects on tryptophan metabolism regulations.**

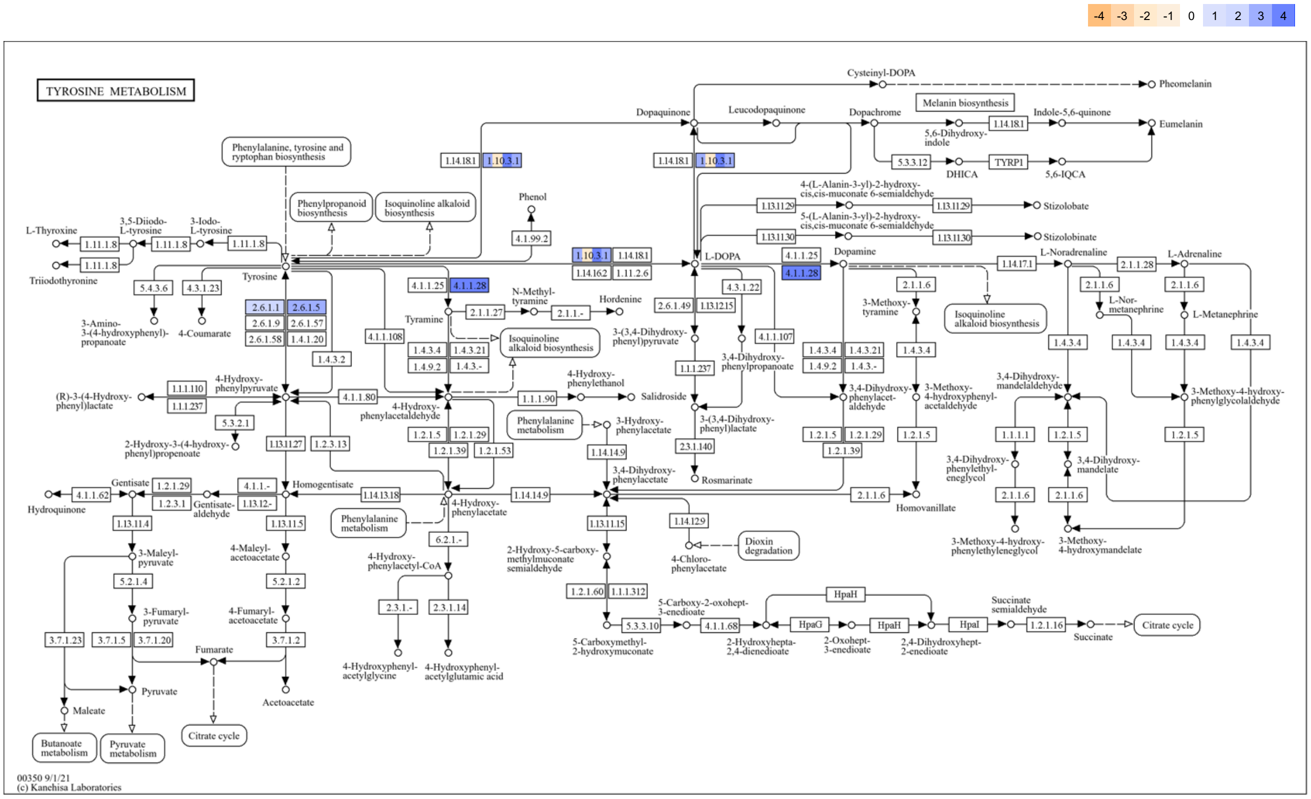


**Fig. S13 KEGG predicted PQQ effects on tyrosine metabolism regulations.**

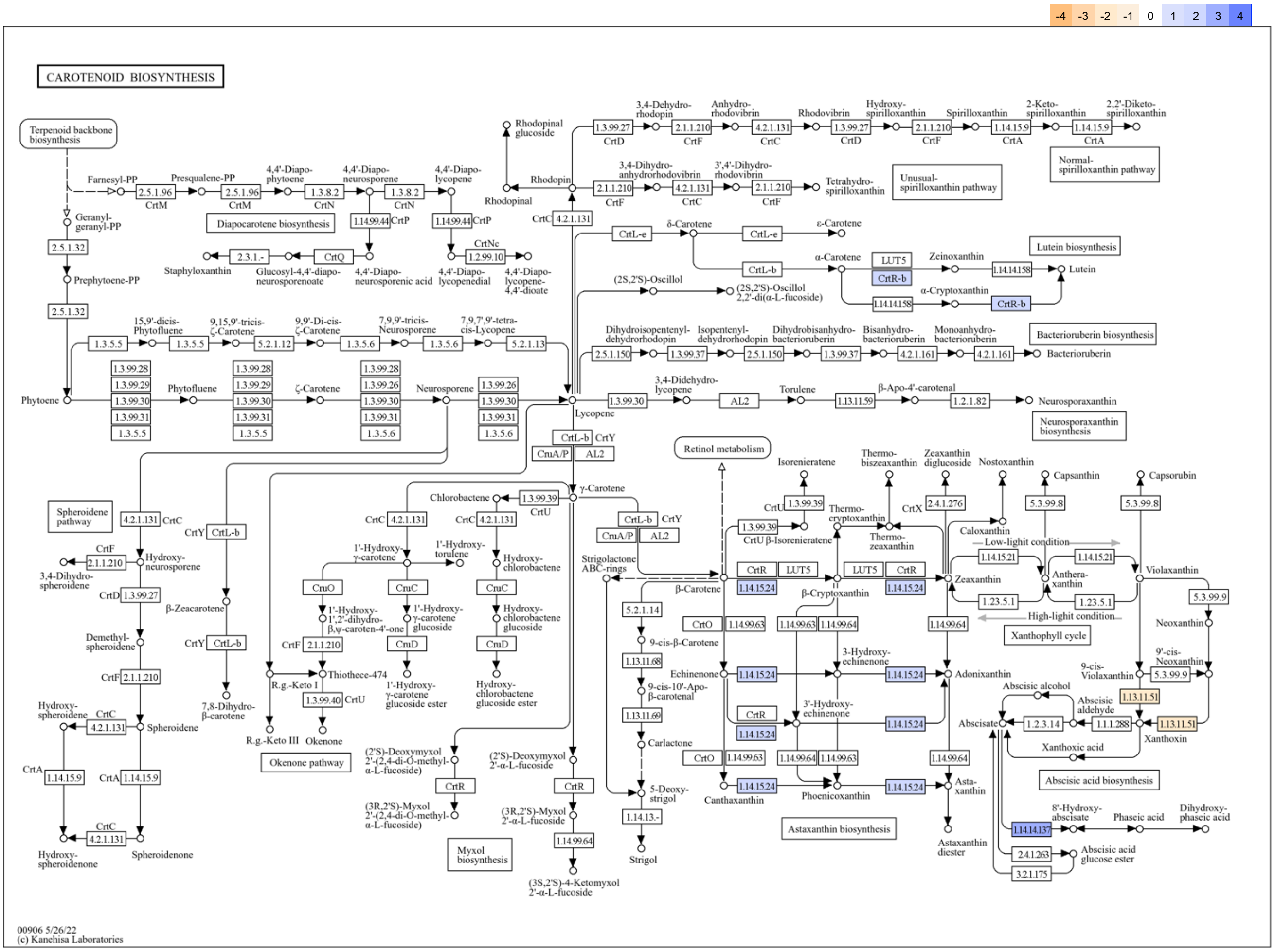


**Fig. S14 KEGG predicted PQQ effects on carotenoid biosynthesis regulations.**

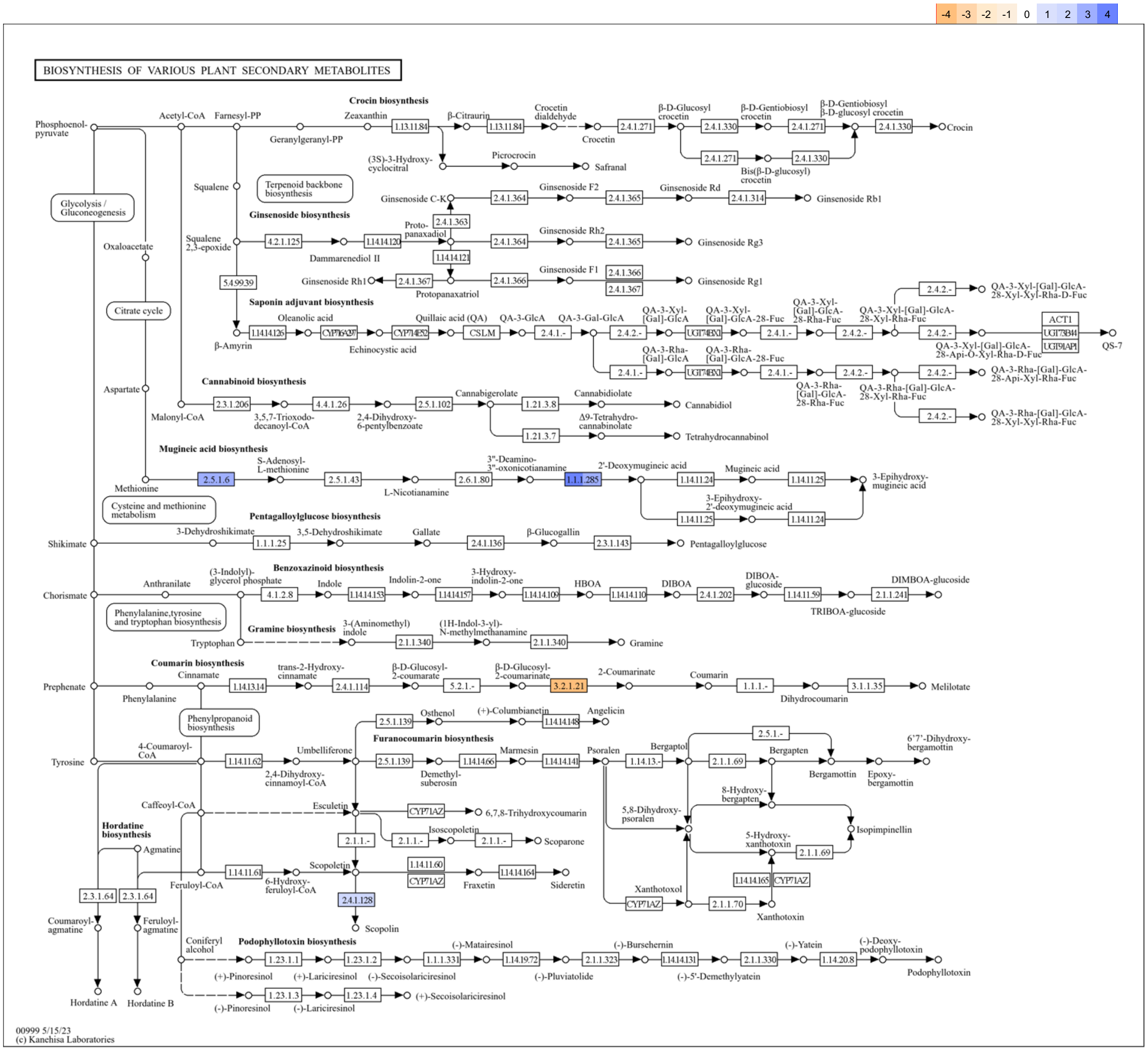


**Fig. S15 KEGG predicted PQQ effects on the biosynthesis of various plant secondary metabolites.**

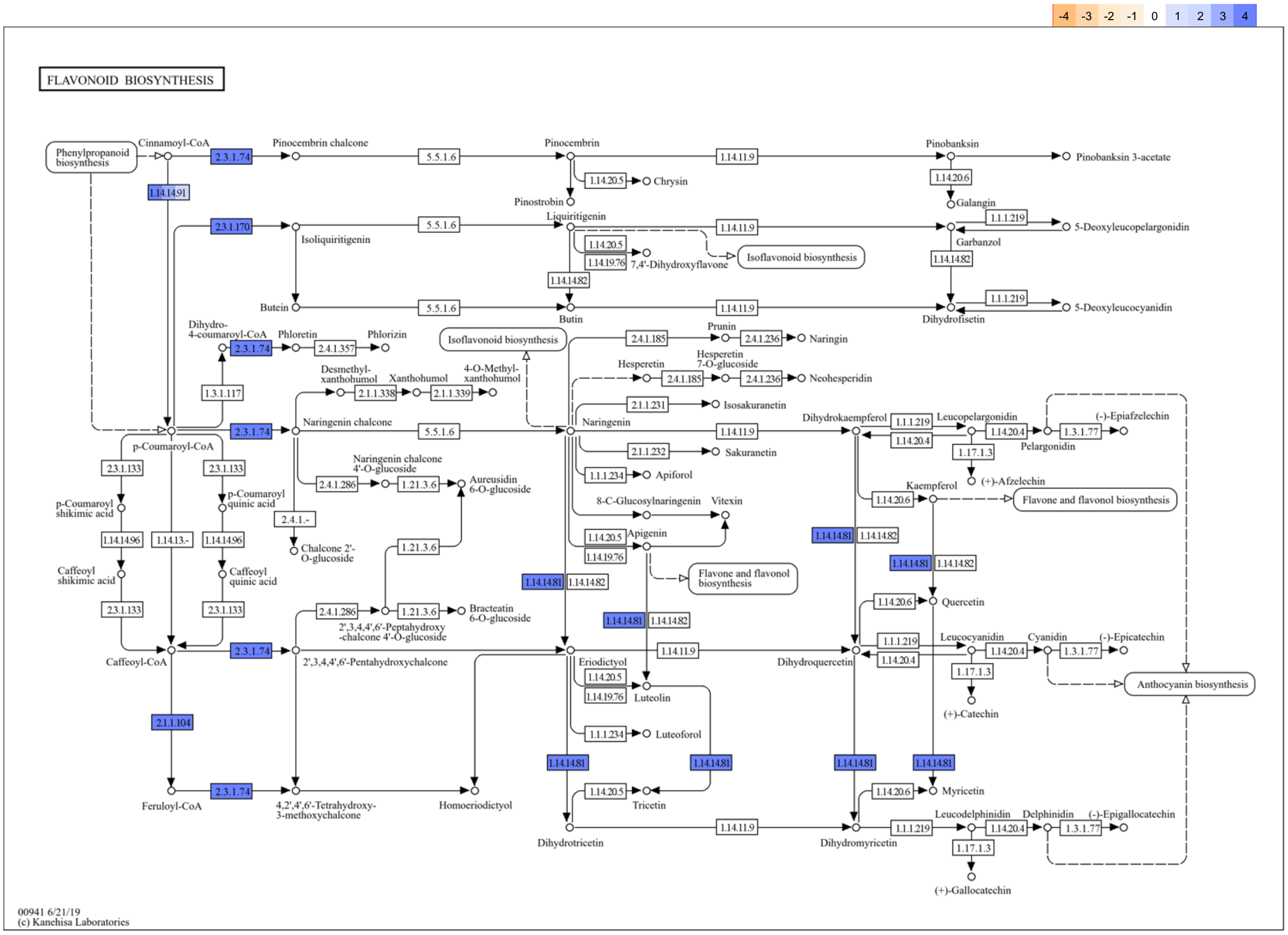


**Fig. S16 KEGG predicted PQQ effects on flavonoid biosynthesis regulations.**

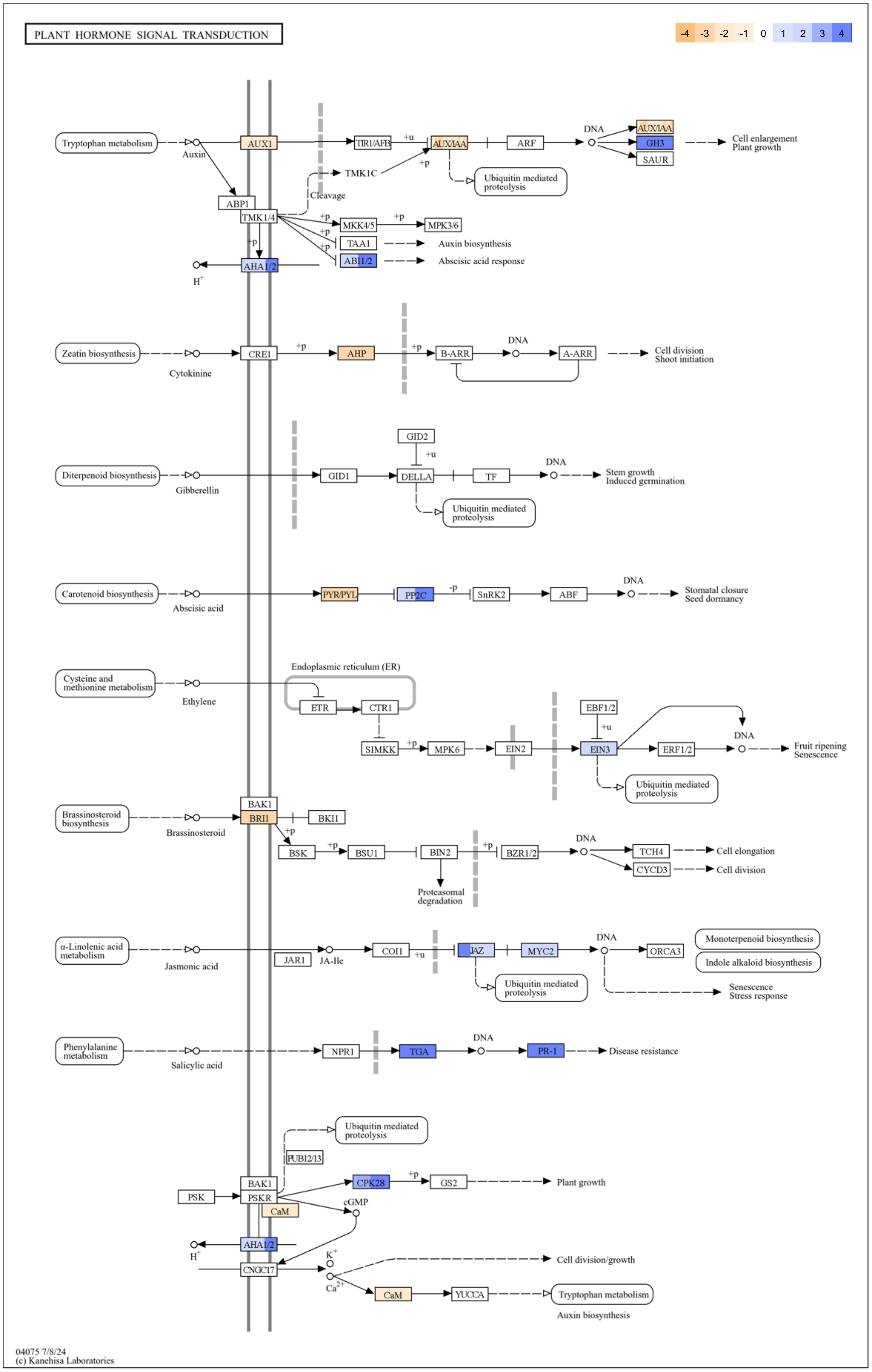


**Fig. S17 KEGG predicted PQQ effects on plant hormone signal transduction regulations.**

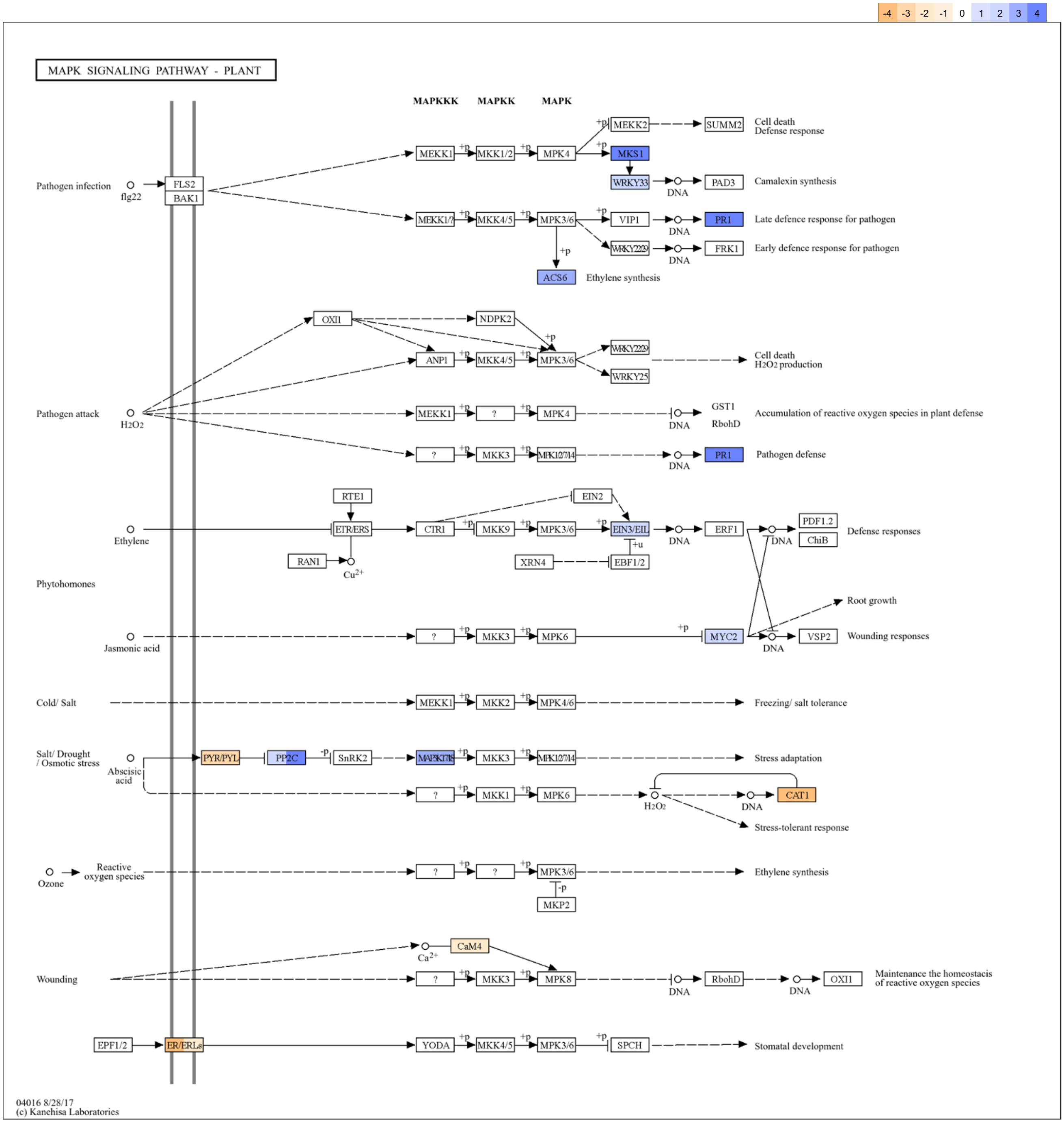


**Fig. S18 KEGG predicted PQQ effects on MAPK signalling pathway – plant regulations.**

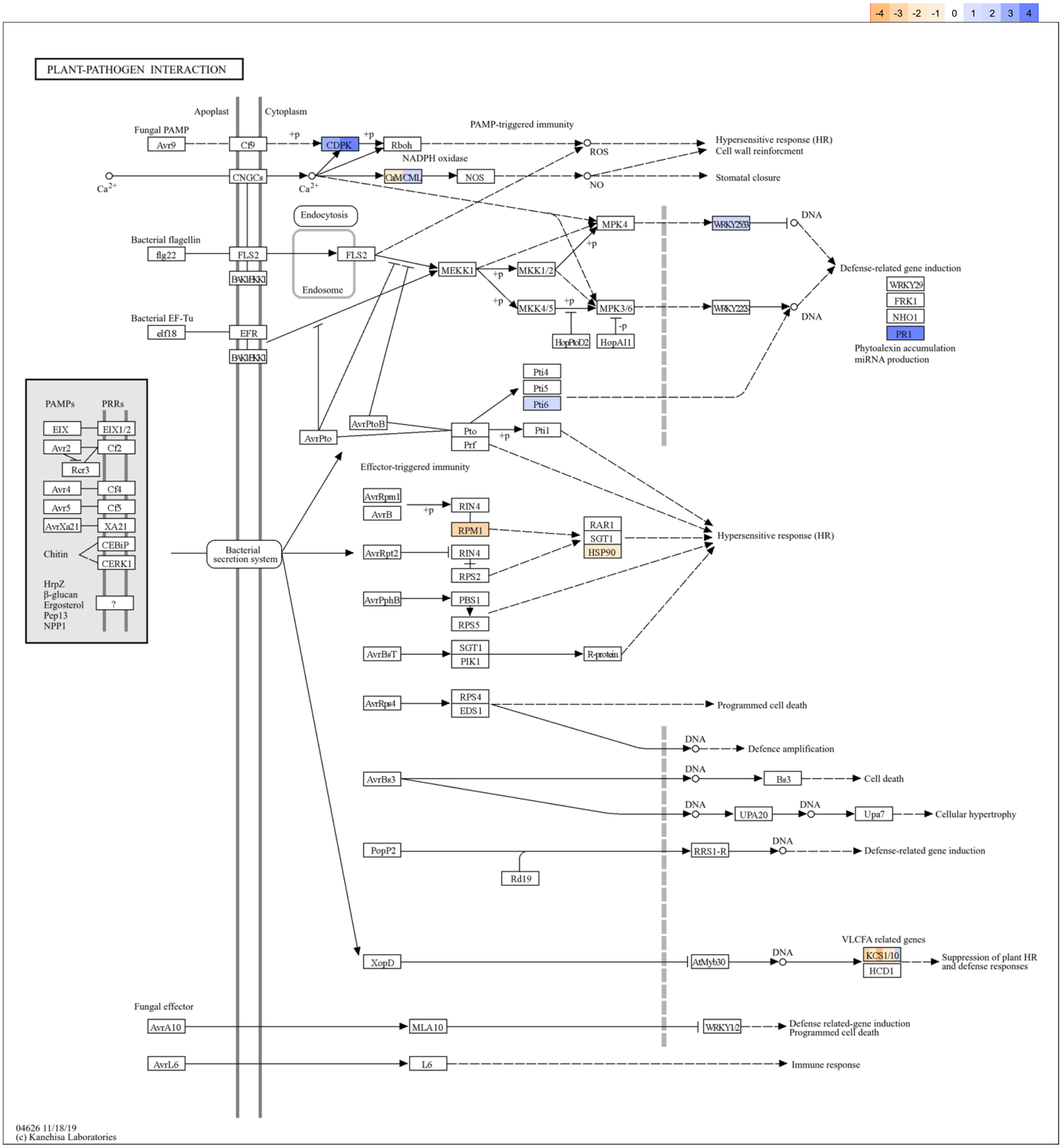


**Fig. S19 KEGG predicted PQQ effects on plant-pathogen interaction regulations.**

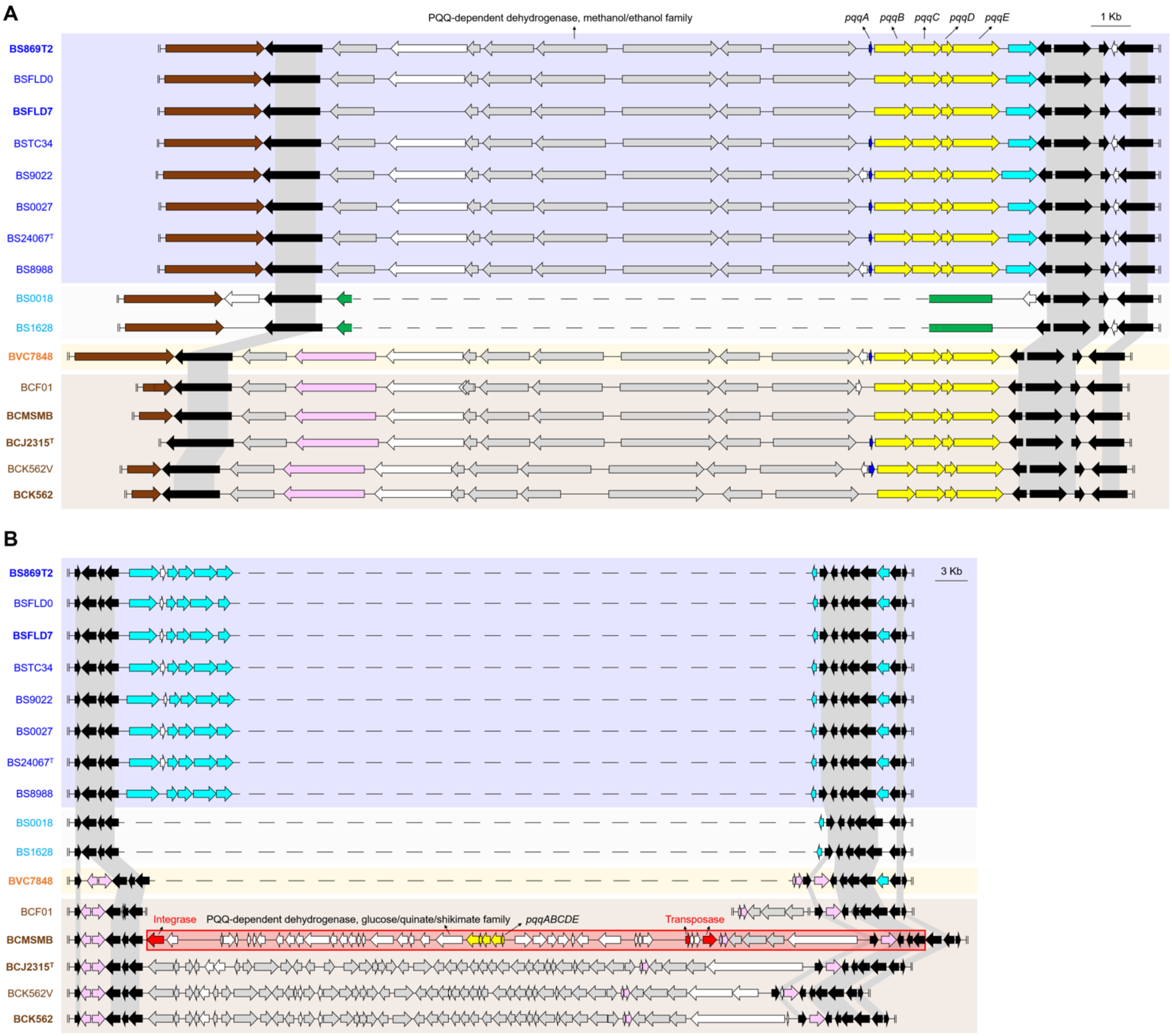


**Fig. S20 Organisation of the PQQ operon among *Burkholderia* species.** The analysed genomes were presented in the same order as the maximum likelihood phylogeny shown in Fig. 1, excluding *Paraburkholderia phytofirmans* PsJN. Distribution of **A** primary and **B** additional PQQ operons (*pqqABCDE*) and their neighbour genes within analysed genomes. The PQQ biosynthesis genes are highlighted in yellow (pqqA was highlighted in dark blue in **A**). Genes uniquely found within *B. seminalis* and *B. cenocepacia* genomes (type strains clades) were shown in fluorescent blue and pink arrows, respectively. Conserved genes at the proposed PQQ operon's neighbouring cassette boundaries were coloured black and linked by grey shadow. The red box and arrows indicated a predicted genomic island and integrase/transposase identified within the genome, respectively.

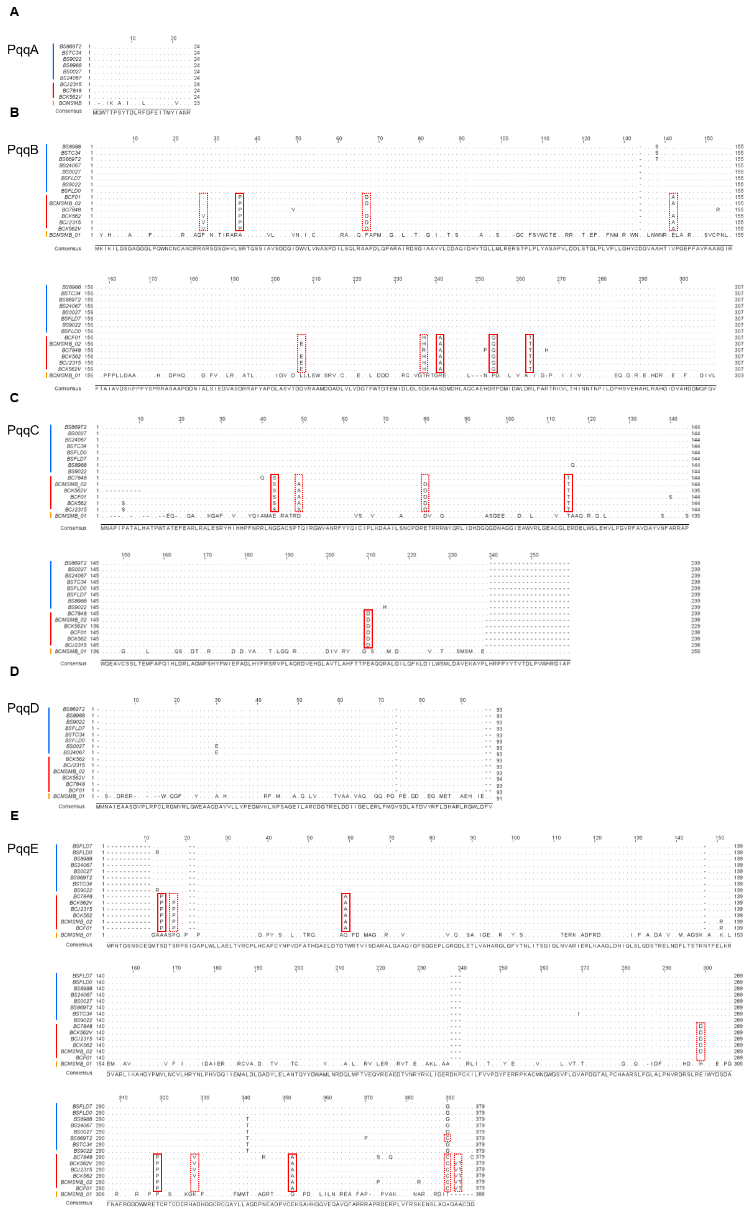


**Fig. S21 Multiple protein sequence alignment of PQQ biosynthesis genes.** Multiple sequence alignment results of **A** PqqA, **B** PqqB, **C** PqqC, **D** PqqD and **E** PqqE were presented. *B. seminalis* and *B. cenocepacia* strains are labelled by blue and red lines along with IDs, respectively; a distinct PQQ operon of the strain MSMB384WGS is shown in orange. Fully conserved single amino acid polymorphisms (SAPs) in analysed *B. cenocepacia* strains are highlighted by boxes with red solid-line; boxes with red dashed-line indicate partially conserved SAPs.

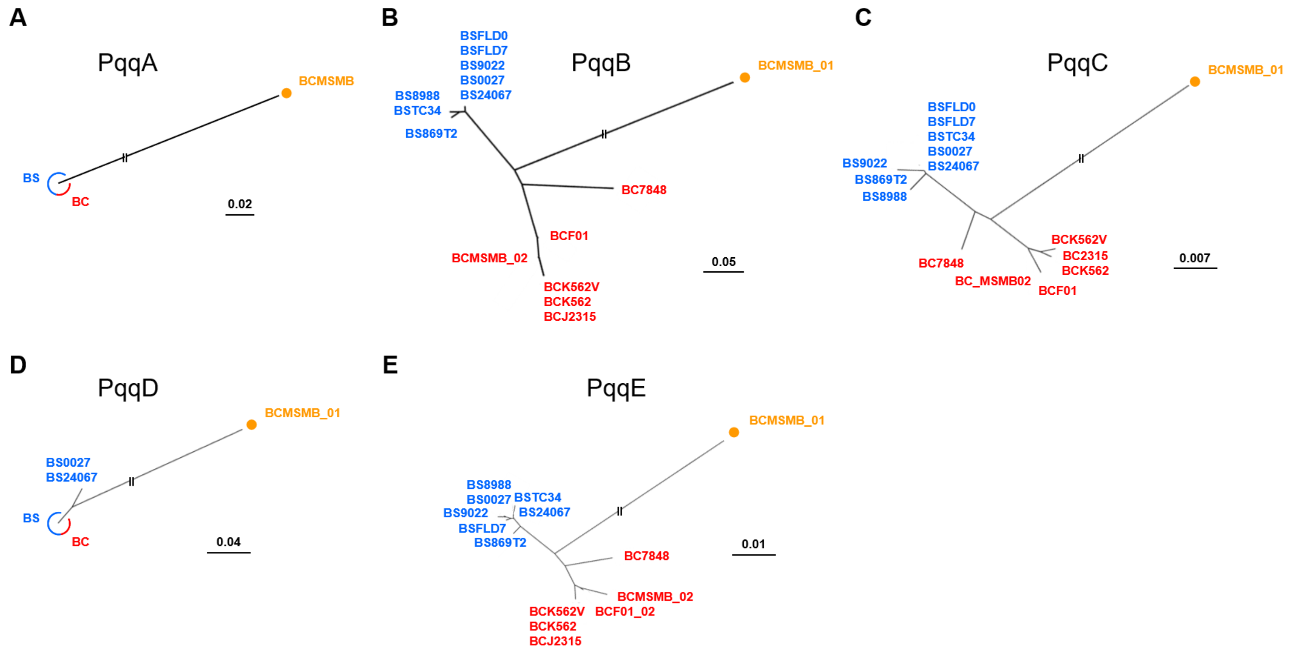


**Fig. S22 Evolutionary divergence of PQQ operon among *Burkholderia* species.** Unrooted maximum likelihood trees of **A** PqqA, **B** PqqB, **C** PqqC, **D** PqqD and **E** PqqE were generated by using multiple protein sequences alignments data in Fig. 13. BS, *Burkholderia seminalis*, shown in blue; BC, *Burkholderia cenocepacia*, shown in red; BCMSMB in **A** and BCMSMB_01 in **B-E** are together embedded a PQQ operon that located on a predicted gene island within *B. cenocepacia* MSMB384WGS.
